# Transthyretin Levels and Instability in Alzheimer’s Disease: Correlations with AD Biomarkers in a Cohort Study

**DOI:** 10.1101/2025.01.24.634710

**Authors:** Tiago Gião, Miguel Tábuas-Pereira, Inês Baldeiras, Joana Saavedra, Alexandre Dias, Maria João Saraiva, Maria Rosário Almeida, Isabel Santana, Isabel Cardoso

## Abstract

**Background:** In the last few decades, Transthyretin (TTR) has gained attention due to its implication in brain functions and neurodegenerative conditions, such as Alzheimer’s Disease (AD) by modulating amyloid-beta (Aβ) pathology. This study aims to assess TTR levels and tetrameric instability in both plasma and cerebrospinal fluid (CSF) across the clinical continuum of AD, and to establish associations between TTR and key AD biomarkers, to enhance our understanding of the role of TTR in AD pathogenesis.

**Methods:** We conducted an evaluation of TTR levels and tetrameric instability in plasma and CSF in mild cognitive impairment (MCI-AD, n=29) and Dementia-AD (n=37) patients and examined the association with clinical, biochemical and genetic data.

**Results:** Key findings revealed a substantial decrease in plasma TTR levels in the Dementia-AD patients compared to the MCI-AD group, with a pronounced gender-specific effect identified in women, while no differences were detected in the CSF. The search for associations identified several correlations amongst MCI-AD patients where CSF TTR levels inversely correlated with markers of amyloid pathology and neurodegeneration such as Aβ40 (r=-0.43, p < 0.02), p-Tau181 (r=-0.54, p < 0.03), t-Tau (r=-0.57, p< 0.001) and NfL (r=-0.49, p<0.006). CSF TTR instability also correlated negatively with CSF Aβ42 (r=-0.6, p<0.001) and Aβ42/Aβ40 ratio (r=-0.58, p<0.001), indicating an association with increased amyloid burden. In the Dementia-AD group, plasma TTR levels correlated negatively with GFAP (r=-0.4, p<0.014), reflecting potential links with neuroinflammation. Additionally, plasma TTR instability correlated with CSF TTR instability (r=0.42, p < 0.009) and tau pathology markers (p-Tau181 (r=0.38, p < 0.02) and t-Tau (r=0.42, p<0.09)). Given the strong association between CSF TTR instability and Aβ42, we investigated the effect of the peptide on TTR tetrameric instability and showed a clear increase in TTR instability in samples incubated with Aβ42 for 24 hours (p<0.05).

**Conclusions:** This study emphasizes several associations between TTR and other key indicators of AD and suggests TTR is associated with Aβ clearance, neurodegeneration, and neuroinflammation across different stages of the disease. We propose Aβ42 and its increase in the brain as a factor destabilizing the TTR tetrameric fold, resulting in TTR impaired function.

## Background

Mild cognitive impairment (MCI) represents a clinical and transitional stage between normal aging and dementia, particularly Alzheimer’s Disease (AD) type, accounting for 60-80% of all dementia cases (1,2). Consequently, MCI stage is a target for dementia prevention and risk reduction research. Approximately 15-26 % of elderly people develop MCI (3–7) with the rates of conversion from MCI to dementia ranging from 10% to 13% annually (7–10). In fact, most of these patients have already neuropathological hallmarks of AD, which lead to the concept of MCI-AD as a prodromic or pre-dementia state of dementia due to AD (Dementia-AD) in the clinical continuum of AD.

In recent years, the scientific community has focused on identifying early biomarkers for AD that are surrogates of neuropathology to facilitate early and accurate diagnosis of neurodegeneration. While current treatments do not reverse pathological changes, the timely implementation of appropriate and personalized interventions can potentially delay the progression of cognitive and functional decline. Early diagnosis could also help in the ongoing search for truly effective disease-modifying therapies for AD, beyond anti-amyloid therapies, which have been relatively unsuccessful in recent decades (11,12).

The most reliable biomarkers for detecting AD are highly discriminative and have significantly advanced our understanding of the disease. These biomarkers are typically identified through neuroimaging methods or by measuring levels of total tau (t-tau), Phosphorylated-tau at threonine 181 (p-tau181), amyloid-β 42 (Aβ42), amyloid-β 40 (Aβ40), and the Aβ42/β40 ratios in cerebrospinal fluid (CSF) (13,14). Promising plasma candidates, including Aβ42, Aβ40, different p-Tau phosphoforms, t-tau, neurofilament light chain (NfL), and glial fibrillary acidic protein (GFAP), provide faster, more accessible testing. This makes continuous monitoring and broader clinical use more feasible, significantly improving early detection and disease management (15).

Transthyretin (TTR) is a homotetrameric protein that serves as a transporter of thyroxine (T4) and retinol-binding protein in both blood and CSF. TTR is primarily synthesized in the liver and choroid plexus, from where it is secreted into the plasma and CSF, respectively (16). This protein constitutes approximately 5% of the protein content in CSF and comprises less than 1% of plasma proteins (17,18). Beyond its transport functions, TTR acts as a neuromodulator (19,20) and plays a critical role in preserving memory function with aging (21,22), demonstrated in rodent models.

In the 1990s, Schwarzman and colleagues identified TTR as the primary sequestering Aβ protein in CSF. It is hypothesized that TTR binds to Aβ, preventing its aggregation and subsequent oligomer formation, thereby playing a protective role against AD pathology (23). Further supporting these findings, *ex vivo* and *in vitro* studies show that TTR binds Aβ, preventing both its aggregation and toxicity (24–29), and cleaves Aβ, generating smaller, less amyloidogenic peptides (30,31). This neuroprotective effect of TTR in AD is supported by *in vivo* studies that show that AD mouse models with reduced TTR expression exhibit increased Aβ production and deposition (32,33). Conversely, overexpression of TTR in AD mice has been shown to reduce both AD neuropathology and Aβ deposition, while improving memory function (34).

At the vascular level, TTR facilitates Aβ clearance from the brain across the blood-brain barrier (BBB) (35) and its absence is associated with vascular impairment, evidenced by thicker basement membranes and reduced vascular density in AD models (36), suggesting that TTR also plays a protective role against vascular alterations associated with AD.

Despite its protective functions, clinical studies report lower plasma/serum TTR levels in AD compared to controls (Ribeiro et al., 2012; Elovaara et al., 1986; Han et al., 2011; Velayudhan et al., 2012) whereas CSF TTR levels yield inconsistent results, confounding the interpretation of the role of TTR in AD pathophysiology.

The underlying cause of decreased TTR levels in AD remains unclear, but it is hypothesized to be linked to tetrameric instability, which accelerates TTR clearance (41) and impairs its interaction with Aβ (24,42,43). Plasma TTR instability has been previously found to be increased in AD patients compared to controls, as indicated by the decreased ability of plasma TTR to carry T4 (42), and higher monomer/folded TTR ratio (43). TTR mutations, such as those seen in TTR amyloidosis, are responsible for TTR destabilization and aggregation, although no TTR mutations are reported in AD patients (44). Thus, other factors may be responsible for TTR instability in AD.

Given these insights, in the current work, we evaluated TTR levels and instability in the plasma and CSF of MCI-AD and Dementia-AD patients to monitor changes as the disease progressed. Demographic and clinical data for each participant were correlated with TTR measurements to explore factors influencing TTR dynamics in AD, thereby enhancing our understanding of the role of TTR in AD pathogenesis.

## Methods

### Subjects

This study included a cohort of 66 subjects (29 MCI-AD and 37 Dementia-AD patients) that were recruited at the Neurology Department of Coimbra University Hospital (HUC), Coimbra, Portugal. Patients were diagnosed in accordance with standard criteria and based on CSF-AD biomarkers: the framework for MCI due to AD and Dementia due to AD, proposed by NIA-AA criteria (45,46). The baseline study and follow-up protocol have already been published elsewhere (47,48).

Patients enrolled had biannual clinical observations and annual neuropsychological and functional evaluations. All patients underwent a thorough biochemical, neurological and imaging (CT, MRI and SPECT) evaluation and performed a lumbar puncture for CSF-AD biomarkers determination and genetic studies (APOE). Amyloid-PET and mutation analysis were more restricted, although considered respectively in patients without inconclusive CSF-biomarkers and younger patients. At baseline, a neurologist completed a medical history with the patient and the caregiver, and conducted a general physical, neurological and psychiatric examination as well as a comprehensive diagnostic battery protocol, including: cognitive instruments such as the Mini Mental State Examination (MMSE) (49) Portuguese version (50), The Montreal Cognitive Assessment (MoCA) (51) Portuguese version (52), the Alzheimer Disease Assessment Scale—Cognitive (ADAS-Cog) (53,54) Portuguese version (55) and a comprehensive neuropsychological battery with normative data for the Portuguese population (Lisbon Battery for Dementia Assessment (BLAD)) (56) exploring memory (Wechsler Memory Scale subtests) and other cognitive domains (including language, praxis, executive functions and visuoconstructive tests); and standard staging scales which provide objective information about subject performance in various domains, including the Clinical Dementia Rating scale (CDR) (57) for global staging, the Disability Assessment for Dementia (DAD) (58,59) for evaluation of functional status and the Neuropsychiatric Inventory (NPI) (60,61) to characterize the psychopathological profile, including the presence of depression. All of the available information (baseline cognitive test, staging scales, clinical laboratory and imaging studies) was used to reach a consensus research diagnosis, supported by CSF-AD biomarkers.

Dementia-AD participants were diagnosed according to the Diagnostic and Statistics Manual for Mental Disorders—fourth edition text review (DSM-IV-TR) criteria, and AD according to the 2011 NIA-AA criteria (46). These cases were classified as probable AD dementia according to clinical, CSF-AD biomarkers and neuroimaging features already referred. Severity of dementia (mild, moderate or severe) was based on the Clinical Dementia Rating scale (CDR) (57) for global staging.

MCI patients included in this study were of the amnestic type and the diagnosis was made in accordance with the framework for MCI due to AD proposed by NIA-AA criteria (45).

As exclusion criteria for enrollment of participants, besides being in a unstable condition namely by acute comorbidities, we considered a significant underlying medical or neurological illness revealed by laboratory tests or imaging besides AD; a relevant psychiatric disease, including major depression, suggested in the medical interview and confirmed by the GDS; CT or MRI demonstration of significant vascular burden (62).

### Samples collection

CSF and blood samples were collected from patients as part of their regular clinical diagnostic evaluation. Pre-analytical and analytical procedures were done according to BIOMARKAPD guidelines for CSF-AD biomarkers (63).

CSF Samples were collected in sterile polypropylene tubes, immediately centrifuged at 1800 g (10 min at 4°C), aliquoted into polypropylene tubes and stored at –80°C until analysis.

Blood samples were collected into plasma separation and EDTA tubes on the same day as the lumbar puncture, centrifuged at 1800 g for 10 min at 4°C, aliquoted into polypropylene tubes, and stored at −80°C.

### Apolipoprotein E (APOE) genotyping

For Apolipoprotein E (APOE) genotyping. DNA was isolated from whole EDTA blood using a commercial kit (Roche Diagnostics GmbH, Manheim, Germany), as described by the manufacturer. The analysis of the two polymorphisms at codons 112 and 158 of the APOE gene (rs429358 and rs7412) was performed by PCR-RFLP assay, as described previously (64).

### Analyses of AD core biomarkers

CSF Aβ42, Aβ40, t-Tau and p-Tau181 were measured separately by commercially available fully automated chemiluminescence enzyme immunoassays (LUMIPULSE, Fujirebio, Japan), as previously described (65). External quality control of the assays was performed under the scope of Alzheimer’s Association Quality Control Program for CSF Biomarkers (66).

Previously determined laboratory specific cut-offs for LUMIPULSE (65) were used to dichotomize markers as normal (-) or abnormal (+) and to classify samples according to the ATN scheme (67): the Aβ42/Aβ40 ratio for evidence of amyloid deposition (A), p-Tau181 for evidence of Tau aggregation (T) and t-Tau for evidence of neurodegeneration (N). According to this scheme, all samples used on this study were classified as being A+T+N+, therefore as having biological AD.

### Analyses of NfL and GFAP

NfL and GFAP CSF levels determinations were done in duplicate, using different aliquots than the one used for CSF-AD biomarkers. For NfL, CSF was diluted 1:1 and NfL was quantified using an ELISA assay (NF-light; Uman Diagnostics, Sweden), as previously described (68). For GFAP, samples were diluted 40 times and measurements were made by single molecule array in a SR-X platform (Quanterix) using the GFAP discovery kit (Quanterix) in accordance with the manufacturer’s instructions and sample type specifications.

### Analysis of the CSF/serum albumin quotient (QAlb)

Albumin plasma levels were determined in duplicate using bromocresol purple (BCP Albumin Assay Kit (Sigma-Aldrich), while albumin CSF levels were measured by sandwich ELISA kits (Proteintech), both following the manufacturer’s instructions. The CSF and serum albumin from 66 paired samples were used to calculate the QAlb using the formula QAlb *=* [albumin (mg/mL)] CSF/[albumin (mg/mL)] plasma × 1000, and used as blood-brain barrier (BBB) measure.

### Transthyretin measurement

Plasma and CSF TTR levels were measured in duplicate by commercially available sandwich ELISA (Abcam), following the manufacturer’s instructions.

### Evaluating Plasma and CSF TTR instability

To assess TTR instability, 4 μL of plasma or CSF was added to SDS reducing gel loading buffer (SDS final concentration = 0.5%) and samples were heated to 55°C for 5 minutes. Proteins were then separated using 15% SDS-polyacrylamide gel electrophoresis (PAGE) and transferred to a PVDF membrane (Thermo Fisher) using a dry system (iBlot, Thermo Fisher). The membranes were then blocked for 1 hour at room temperature with 5% powdered skimmed milk in PBS containing 0.05% Tween-20 (PBS-T). Immunoblotting was then performed using a mouse anti-TTR antibody (69) to detect monomer and dimer TTR. The blots were developed using Clarity™ Western ECL substrate (Bio-Rad), and proteins were detected and visualized using a chemiluminescence detection system (ChemiDoc, Bio-Rad). Images were analyzed and band intensities were quantified using Image Lab (Bio-Rad, version 4.1). A ratio of TTR monomer/dimer intensities was used to determine TTR instability.

### Preparation of soluble Aβ42

Synthetic Aβ42 (Genscript) was dissolved in hexafluoro-2-propanol (HFIP) (Sigma-Aldrich) and kept at RT over weekend. After, the HFIP was removed under a stream of nitrogen and the powder was dissolved in DMSO at 2 mM. Soluble Aβ42 was obtained by diluting the peptide in Ham’s F12 medium. To confirm the presence of soluble Aβ42 species, samples were analysed by transmission electron microscopy (TEM) and visualized by negative staining using uranyl acetate. Briefly, 5 μl aliquots were adsorbed into carbon coated collodion film supported on 300-mesh copper grids (Electron Microscopy Sciences, PA, USA) and negatively stained twice with 1% (m/v) uranyl acetate (Electron Microscopy Sciences, PA, USA). Grids were visualized with a JEOL (Tokyo, Japan) JEM1400 transmission electron microscope equipped with an Orious (CA, USA) Sc1000 digital camera, and exhaustively observed.

### Assessment of tetrameric TTR instability when incubated with Aβ42

To assess the instability of tetrameric TTR in the presence of Aβ42, 10 µM of recombinant wild-type TTR (WT TTR, AlexoTech) was incubated at 37°C overnight, either alone or with 40 µM of Aβ42 (Genscript), in duplicate. The samples were then subjected to semi-denaturing electrophoresis and Western Blot, as previously described for evaluating plasma and CSF) TTR instability.

### Statistical analysis

Categorical variables were expressed as count (%) and compared with Chi-square test or Fischer’s Exact test, as appropriate. Continuous variables were expressed as mean (standard deviation) and compared using Student’s t-test or Mann-Whitney test, as appropriate. Correlations were calculated by the Spearman’s correlation test. The statistical significance threshold was set at P<0.05, two-sided. The analyses and figures were made using SPSS Statistics version 29 (IBM) and Graphpad Prism 8.

## Results

### Demographics and clinical description of the participants

A total of 66 participants with a biological diagnosis of AD supported by CSF biomarkers were included in this study, comprising 29 with Mild Cognitive Impairment (MCI-AD) and 37 with dementia (Dementia-AD). Demographic and clinical information is summarized in Table 1, categorized by clinical staging MCI-AD versus Dementia-AD. No statistically significant differences were found between the groups regarding sex or education level.

**Table 1.**
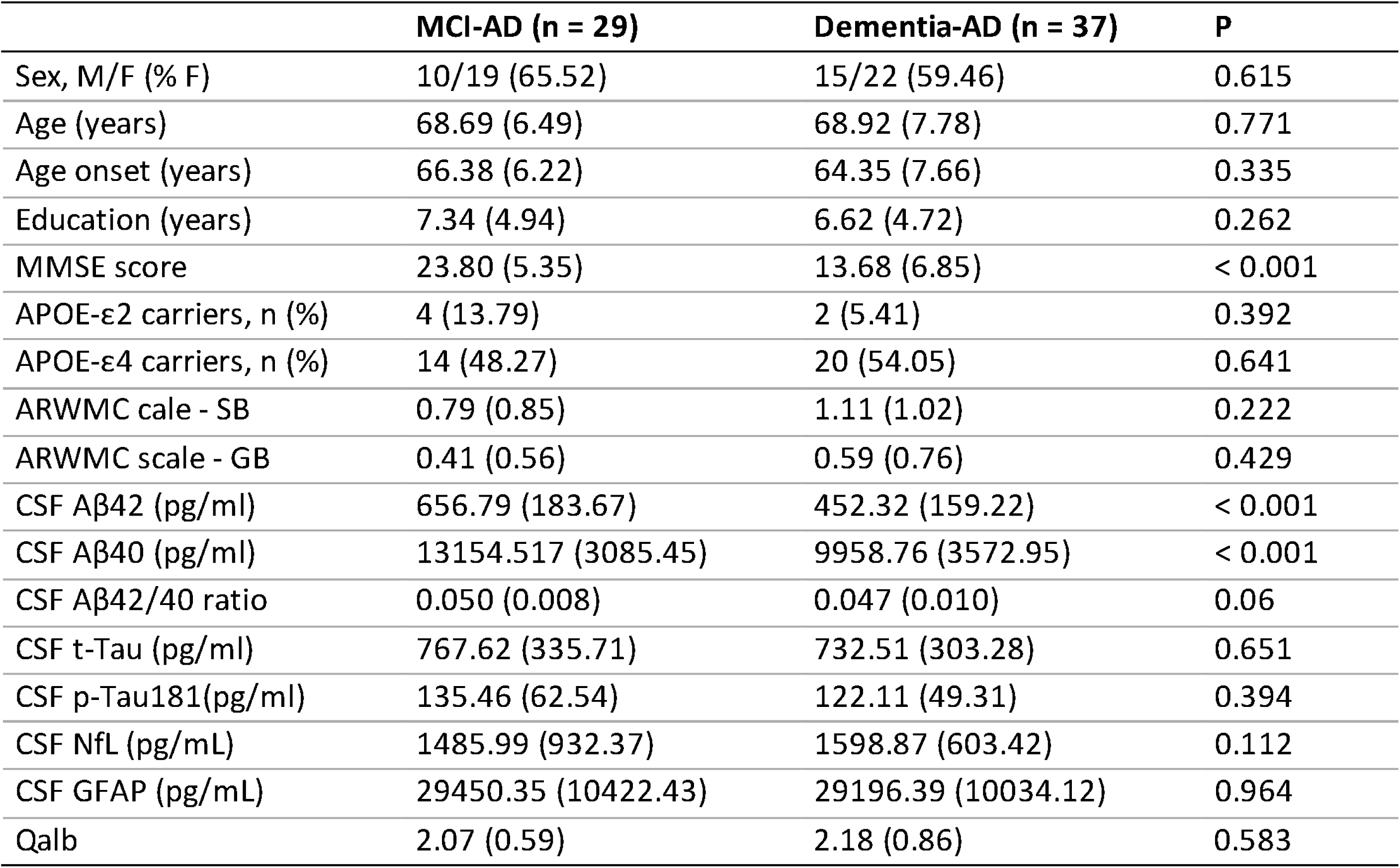
Demographic and clinical data from the cohort.

Several comorbidities were documented among participants, with significant decreases observed in plasma and CSF glucose levels in Dementia-AD patients (Supplement, Table S1).

### Evaluation of CSF and plasma TTR levels and instability

Plasma TTR levels were measured, revealing lower levels in Dementia-AD subjects compared to those with MCI-AD (p=0.024) (Fig. 1A). Stratification by gender showed that this difference was predominantly influenced by females, who exhibited a similar pattern of lower TTR levels in MCI-AD versus Dementia-AD patients (p=0.036) (Fig. 1B), while no significant differences were observed among males (Fig. 1C). CSF analysis revealed no significant differences in TTR levels between MCI-AD and Dementia-AD groups (Fig 1. D), even when divided by gender (Fig. 1E, F). The analysis revealed no significant differences in TTR instability between the diagnostic groups in both fluids (Fig. 1G, H, I, J), although, there was a trend indicating that CSF TTR may be less stable in Dementia-AD patients.

**Figure 1.**
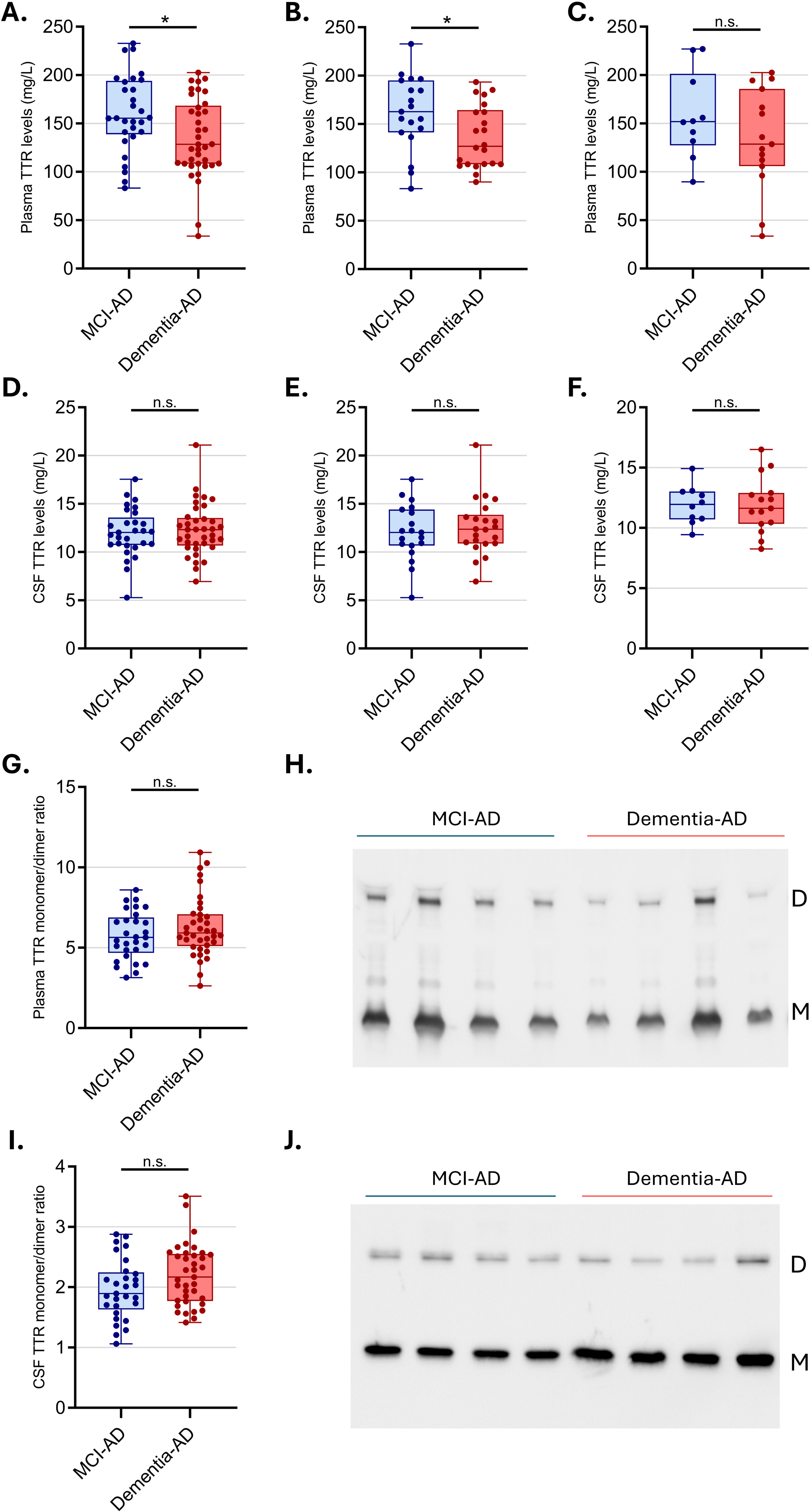
Evaluation of TTR levels and instability in plasma and CSF according to clinical diagnosis. Box-plots comparing plasma TTR levels in **A.** MCI-AD (n=29) and Dementia-AD (n=37) groups; **B.** MCI-AD (n=19) and Dementia-AD (n=22) in females; and **C.** MCI-AD (n=10) and Dementia-AD (n=15) in males. Box-plot comparing CSF TTR levels in **D.** MCI-AD (n=29) and Dementia-AD (n=37) groups; **E.** MCI-AD (n=19) and Dementia-AD (n=22) in females; and **F.** MCI-AD (n=10) and Dementia-AD (n=15) in males. Box-plots comparing TTR instability in **G.** plasma and **I.** CSF, in MCI-AD (n=29) and Dementia-AD (n=37) groups, by measuring the monomer/dimer ratio. The boxplots depict the median (horizontal line within each box), interquartile range (IQR, box ends) and 1.5 × IQR (whiskers). Asterisks indicate significant differences, where * *p*⍰<⍰0.05 by two-tailed Mann-Whitney test. Western blot of TTR in **H.** plasma and **J.** CSF samples from the groups analyzed, exhibiting monomer (M) and dimer bands (D) used for analysis.

### Correlation of TTR levels and instability with CSF AD biomarkers and other CSF candidates

The availability of clinical data on CSF core AD biomarkers, as well as CSF NfL, and GFAP— indicators of neurodegeneration and glial activation, respectively, enabled the establishment of correlations between TTR levels and instability in both plasma and CSF.

As shown in Fig. 2A, in the combined cohort of MCI-AD and Dementia-AD patients, there was a negative correlation between plasma TTR levels and CSF GFAP (r=-0.33, p<0.007). Additionally, plasma TTR instability showed a modest correlation with CSF p-Tau181 (r=0.34, p<0.006), CSF t-Tau (r=0.34, p<0.006) and CSF TTR instability (r=0.26, p<0.03). CSF TTR instability showed a weak correlation with CSF Aβ42 levels (r = −0.28, p<0.02).

**Figure. 2.**
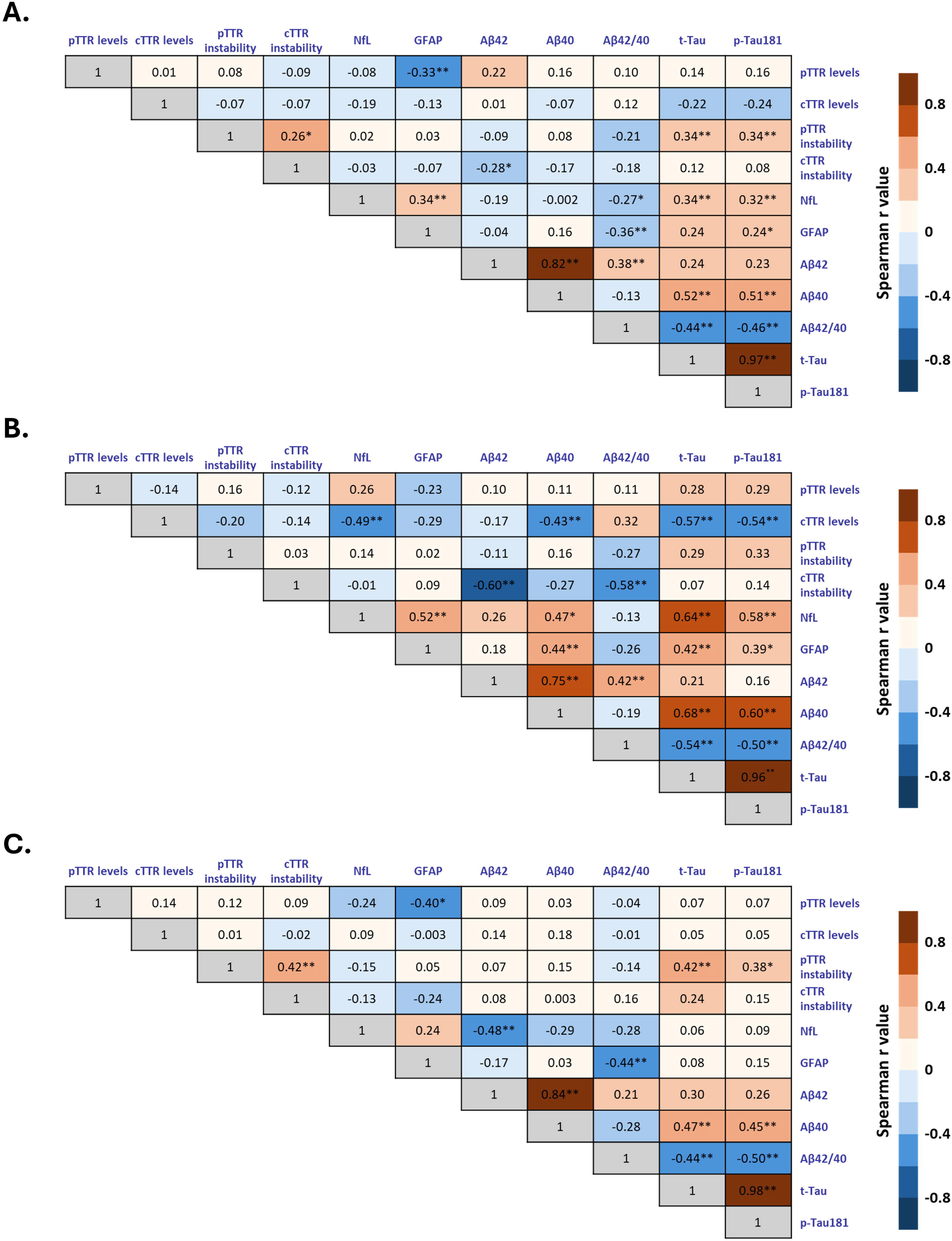
Correlation between plasma (pTTR) and CSF TTR (cTTR) instability and levels with core AD biomarkers in A. MCI-AD and Dementia-AD (n=66) group; **B.** MCI-AD (n=29) group; **C.** Dementia-AD (n=37) group. The correlation matrix graphically represents the Spearman’s correlation coefficient (r) for each pairwise comparison among the biomarkers. According to the color scale on the right side of the matrix, positive and negative correlations are indicated in shades of red and blue, respectively; an r value of +1 indicates a perfect positive relationship, r = 0 indicates no relationship, and an r value of −1 indicates a perfect negative relationship. Asterisks indicate significant differences, where *p⍰<⍰0.05 and **p⍰<⍰0.01. Abbreviations: Aβ - amyloid-β, AD - Alzheimer’s disease, CSF - cerebrospinal fluid, GFAP - Glial fibrillary acidic protein, MCI - mild cognitive impairment, NfL - neurofilament light chain, p-tau - phosphorylated tau, t-tau - total tau.

In the MCI-AD group (Fig. 2B), CSF TTR levels inversely correlated with CSF p-Tau181 (r=-0.54, p<0.03), t-Tau (Fig 2G, r=-0.57, p<0.001), Aβ40 (r=-0.43, p<0.02), and CSF NfL (r=-0.49, p<0.006). CSF TTR instability correlated with CSF Aβ42 (r=-0.6, p<0.001) and Aβ42/Aβ40 ratio (r=-0.58, p<0.001).

A similar analysis was conducted for the Dementia-AD group (Fig. 2C), revealing that plasma TTR levels associated with CSF GFAP (r=-0.4, p<0.014). Additionally, plasma TTR instability correlated moderately with CSF TTR instability (r=0.42, p<0.009), and CSF p-Tau181 (r=0.38, p<0.02) and t-Tau (r=0.42, p<0.09). Other correlations between TTR levels and instability have been identified across several demographic and clinical variables (Supplementary Fig. S1), of which we highlight the positive relation between plasma TTR instability with women in the combined MCI-AD patient cohort and in the Dementia-AD group (r = 0.29, p<0.02, Fig. S1A) and in the MCI group (r = 0.51, p<0.005, Fig. S1B). Furthermore, plasma TTR instability was negatively correlated with the age of disease onset in the MCI-AD group (r=-0.38, p<0.045, Fig. S1B).

### Evaluating TTR tetramer destabilization by Aβ

To address the question of whether or not TTR instability can be influenced by Aβ peptide, recombinant TTR (10 μM) was incubated with or without Aβ42 (40 μM) for 24 hours. TTR instability was assessed using semi-denaturing gel electrophoresis, followed by Western blot analysis. Results showed a clear decrease in TTR instability in samples incubated with Aβ42 for 24 hours (Fig. 3).

**Figure 3.**
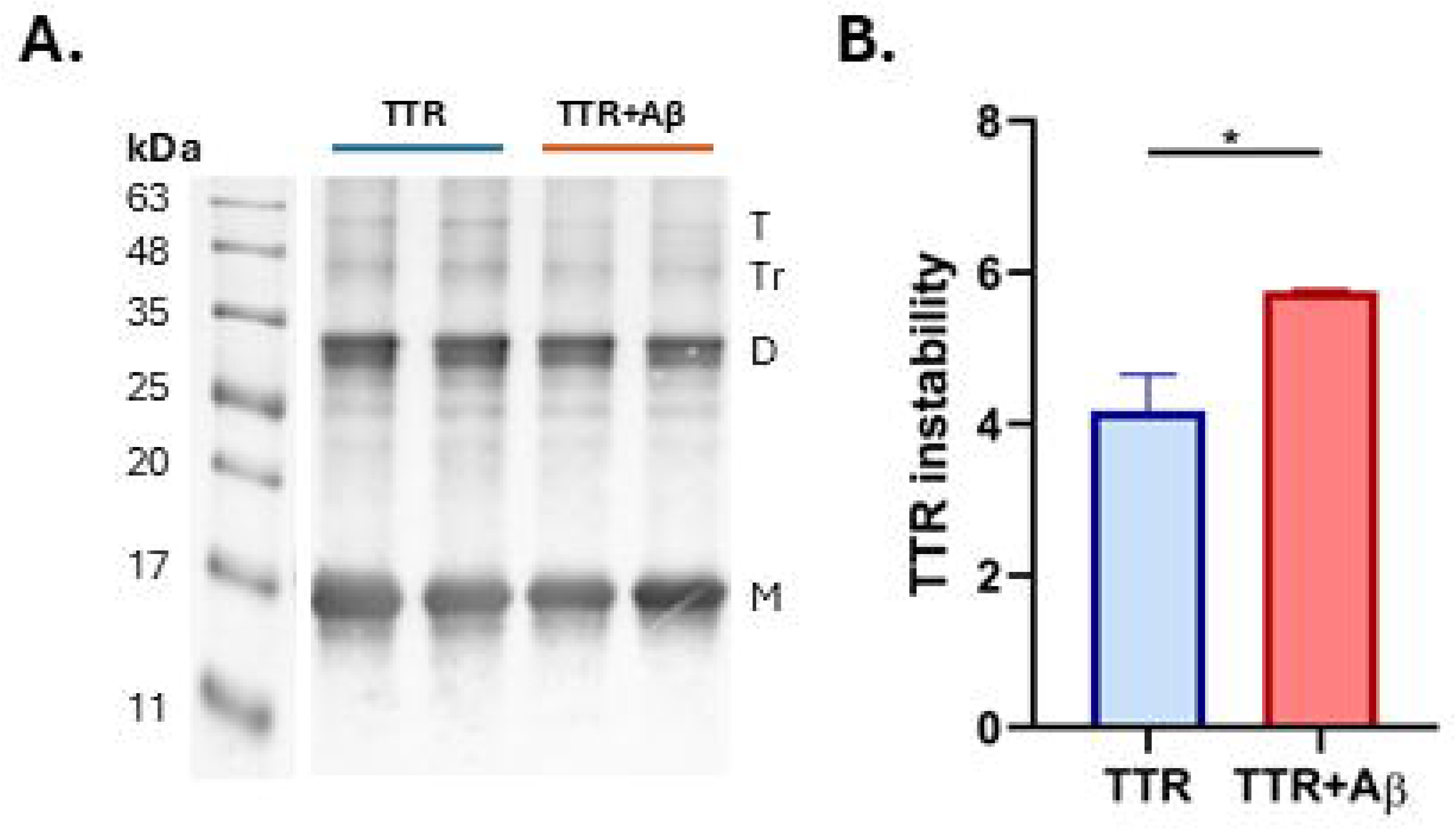
Effect of TTR binding to Aβ42 peptide and consequences in TTR tetrameric instability. **A.** Western blot analysis of TTR (10 μM) alone or in combination with Aβ42 (40 μM) after incubation at 37°C for 24 hours, detected using an anti-TTR antibody. Samples were subjected to semi-denaturing gel electrophoresis. The positions of TTR monomer (M), dimer (D), trimer (Tr), and tetramer (T) are indicated on the right side of the blot, with molecular weights in kDa marked on the left. **B.** Graphical analysis quantifying TTR instability, represented by the monomer+dimer/tetramer intensity ratio. Data are presented as mean ± SD. Asterisk indicate significant differences, where * p⍰<⍰0.05 by unpaired Student’s t test.

## Discussion

Our study found significantly reduced plasma TTR levels in Dementia-AD patients compared to MCI-AD subjects. These findings are in line with earlier studies (70,71), although some research has reported no significant differences between these groups (72). Our result suggests a progressive decrease in TTR levels along the AD continuum, as supported by several studies that have documented lower plasma TTR levels in AD patients relative to controls (37–40,70). Interestingly, our study is the first to report sex-specific differences within the AD continuum, where female Dementia-AD subjects exhibited lower TTR levels compared to their MCI-AD counterparts. Previous studies have detected significant decreases in TTR levels among women with AD compared to controls, although differences were not found between with MCI and AD. This decrease during the disease progression supports the hypothesis that women are at a higher risk of developing AD, potentially due to reduced levels of estradiol (73), which regulates TTR expression in the liver and choroid plexus (74,75). This gender-specific observation is complemented by findings from animal models where estradiol treatment increased TTR levels in the brain, along with reduced APP metabolism, lower Aβ42 levels, and decreased plaque burden (76).

Concerning CSF TTR levels, our study found no changes with disease progression. CSF TTR levels have been reported to be lower in AD compared to MCI patients (77,78). However, results across studies have been inconsistent, with some reporting lower CSF TTR levels in AD compared to controls (78–81), and others finding no difference or even increased levels (82–84). This variability is further complicated by the different methodologies used across studies, such as ELISA, radial immunodiffusion, nephelometry, and mass spectrometry employed in different settings, highlighting the urgent need for standardized measurement techniques to ensure reliable and reproducible results.

Regarding TTR instability, a key factor in the TTR/Aβ interaction, it is interesting to note that, although our study did not detect significant differences in protein instability between the groups, we observed a trend toward increased instability of TTR in the CSF of Dementia-AD patients compared to those with MCI-AD. This suggests a potential alteration in TTR dynamics as the disease progresses, justifying further investigation into the role of TTR stability in the pathophysiology of AD. Previously, our laboratory observed that reduction in plasma TTR levels in AD patients compared with controls was associated with increased instability, suggesting a link between TTR instability and its clearance rates (35,72). Furthermore, we demonstrated that Aβ can destabilize the recombinant TTR tetrameric structure, aligning with the observed trend of peptide accumulation during disease progression. These results are consistent with previously reported data that verified TTR destabilization after 24 hours in the presence of Aβ (85). This observation is fundamental as it links TTR dynamics directly to the pathological processes underlying AD.

To better understand the complex dynamics of TTR in relation to AD pathology, TTR levels and instability were correlated with core AD CSF biomarkers and CSF NfL and GFAP, offering insights into proteins related to TTR dynamics in AD pathology. Correlation analysis in the MCI-AD group revealed that CSF TTR levels were negatively associated with CSF Aβ40, p-Tau181, t-tau, and NfL. Additionally, CSF TTR instability showed a strong negative correlation with CSF Aβ42 levels and the Aβ42/40 ratio.

It can be speculated that in the early stages of AD pathogenesis, CSF TTR may sequester Aβ42 and inhibit amyloid formation. Conversely, impaired TTR tetrameric stability, induced by this interaction, may lead to Aβ42 accumulation, aggregation, and amyloid formation and, consequently, reduced CSF Aβ42 levels. Interestingly, Aβ40, which is less amyloidogenic than Aβ42, correlates with CSF TTR levels, suggesting that the interaction with Aβ42 may depend on TTR stability, while Aβ40 is related to its levels. Although we did not find correlations between TTR levels and Aβ in Dementia-AD group in our study, previous research has shown that CSF TTR levels negatively correlate with disease severity and senile plaque abundance (78,80), and positively correlate with Aβ38, Aβ40, and Aβ42 levels in AD (81).

Markers such as NfL, t-tau protein, and its hyperphosphorylated form, p-Tau181, indicative of neuronal injury show an inverse correlation with CSF TTR levels and suggests that TTR may have a neuroprotective role. This relationship may, therefore, reflect neuronal degeneration, potentially due to amyloid accumulation.

Higher TTR levels in MCI-AD patients compared to those with Dementia-AD may represent an early neuroprotective response to counter Aβ accumulation and its neurotoxic effects. Supporting evidence from AD mouse models reveals increased TTR mRNA in the cortex and hippocampus prior to plaque formation, with a subsequent decline post-deposition (86,87). Similarly, higher plasma TTR levels correlate with increased Aβ deposition in MCI patients, suggesting TTR upregulation as an adaptive response to initial pathological changes (88). At this stage, maintaining TTR levels and stability is crucial for neuronal homeostasis.

Although TTR displays neuroprotective properties in regulating Aβ levels in MCI-AD, this effectiveness declines in Dementia-AD patients. Nonetheless, TTR maintains a consistent inverse relationship with neuronal damage and also begins to show an inverse correlation with markers of glial activation in Dementia-AD patients, as evidenced by a negative correlation with CSF GFAP levels. GFAP, a protein highly specific to the brain that reflects astroglial response and gliosis near Aβ plaque sites (89) is strongly linked with cerebral Aβ pathology (90,91). Our results demonstrate a negative correlation between peripheral TTR levels and GFAP, suggesting the involvement of TTR in AD-related inflammation. Previous work using a model with diminished heat-shock response, reported increased GFAP levels in the hippocampus of TTR KO animals compared to WT, indicating that TTR mitigates astrogliosis under these conditions (92). Changes in TTR levels may directly or indirectly impact astrocyte function and neuronal health.

Interestingly, in AD patients experiencing rapid cognitive decline and severe cognitive impairment, plasma TTR levels are significantly lower, suggesting a critical role in disease progression (37). This decrease in peripheral TTR could be attributed to its regulation in the liver, where TTR synthesis can be modulated independently of the CP. For instance, TTR is downregulated in the liver during the acute phase response to inflammatory conditions, a process that occurs without corresponding transcriptional changes in the CP (93–95). The implications of this independent regulation within the context of AD, however, require further investigation to elucidate the impact on disease progression. This decline in TTR might also be linked to dysfunction of the CP in AD, the primary site of TTR synthesis in the brain, which leads to diminished secretion of TTR into the CSF (96).

TTR can cross the BBB from the brain to the blood, but not significantly in the reverse direction (35). Disruption in TTR efflux from the brain, necessary for TTR turnover and degradation in peripheral tissues (97), may result in its accumulation in blood vessels and senile plaques. Indeed, several studies have shown that TTR colocalizes with Aβ in senile plaques and blood vessels in AD patients, as opposed to non-demented controls, possibly impacting plasma TTR levels (34,86,98,99). Nevertheless, the presence of TTR in these plaques could be beneficial, as interactions between TTR monomers and soluble Aβ aggregates might lead to the formation of non-fibrillar amorphous precipitates, considered a more favorable outcome than fibrillar Aβ (100). Furthermore, recent research suggests that peripheral TTR accumulation might also contribute to cerebral Aβ pathology (101), highlighting the complex role of TTR in the pathogenesis of AD.

### Strengths and Limitations

The primary strength of the study lies in its clinical setting, reflecting real-life patients seeking medical attention in hospital environments. All participants underwent comprehensive diagnostic evaluations with overlapping measures to enhance diagnostic consistency. Although the confirmation of neurodegenerative diagnoses via postmortem autopsies was not possible, analyses of well-validated CSF biomarkers were performed as an alternative. Additionally, the paired analysis of plasma and CSF samples from the same individuals provided insights into dynamic changes in transthyretin (TTR). The paired analysis of plasma and CSF samples offered insights into dynamic transthyretin (TTR) changes, and the inclusion of CSF biomarkers such as NFL and GFAP further enriched the analysis. Vascular and metabolic factors, including white matter and basal ganglia hyperintensities, BBB integrity, diabetes, hypertension, and glucose levels, were also assessed, offering a multifaceted view of their impact on disease progression and TTR.

Despite its strengths, the study has some limitations. The modest sample size limits statistical power and generalizability. Additionally, the higher proportion of female participants (approximately 62%) might have influenced the findings, introducing a potential bias toward female-related results.

The absence of a control group is a significant limitation. The neurological controls that were available were individuals undergoing routine diagnostic lumbar punctures for conditions such as acute or chronic headaches or peripheral polyneuropathy. Although their CSF cytochemical evaluations were normal, and no major CNS diseases were detected. Additionally, their average age was relatively low compared to the patient cohorts. This discrepancy may affect the comparability of findings and limit the broader applicability of the results.

## Conclusions

We propose that TTR acts as a neuroprotective protein against early Aβ accumulation during the MCI-AD stage, but its plasma levels decrease in later Dementia-AD stage for reasons not yet understood. Our results indicate that plasma TTR concentration decreases in Dementia-AD, particularly in women, and shows TTR is associated with key AD markers, such as Aβ pathology, neuroinflammation, and neurodegeneration. These associations differ depending on the stage of the disease, highlighting the complex role of TTR in AD progression.

## Supporting information

Uncroped Blots

## List of abbreviations

Aβ: Amyloid-β
AD: Alzheimer’s disease
APOE: Apolipoprotein E
ARWMC: Age-related white matter changes
CSF: Cerebrospinal fluid
GFAP: Glial fibrillary acidic protein
MCI: Mild cognitive impairment
MMSE: Mini-Mental State Examination
NfL: Neurofilament light chain
p-Tau181: Phosphorylated-tau at threonine 181
Qalb: CSF/plasma albumin quotient
SD: Standard deviation
T4: Thyroxine
t-Tau: Total Tau
TTR: Transthyretin

## Declarations

### Ethics approval and consent to participate

The present research complied with the ethical guidelines for human experimentation stated in the Declaration of Helsinki and was approved by the Ethics Board of Coimbra University Hospital (OBS.SF.088.2022) and Committee for Ethical and Responsible Conduct of Research of i3S (N 16/CECRI/2023). All subjects or responsible caregivers, whichever appropriate, gave their informed consent.

### Consent for publication

Not applicable.

## Declarations

### Consent for publication

Not applicable. informed consent.

### Availability of data and materials

The datasets used and/or analysed during the current study are available from the corresponding author on reasonable request.

### Authors’ Contributions

IC and IS designed and supervised the study. IS and MT-P coordinated clinical assessments and IB, MT-P and IS managed the collection of samples and clinical data from the recruited participants. TG performed TTR- and albumin-related experimental work with assistance from JS, and IB performed overall biomarker quantification. TG and AD performed data analysis and data interpretations with support from MT-P, MRA, MJS, IS and IC. TG, MT-P, IS and IC wrote the manuscript. All authors contributed and critically reviewed the final version of the manuscript and approved the final manuscript.

## Acknowledgements

The authors would like to thank all patients and their caregivers who participated in the study.

