## Supplementary files for "Transthyretin Levels and Instability in Alzheimer’s Disease: Correlations with AD Biomarkers in a Cohort Study"

**Table S1.** Comorbidities and health status of the cohort.

|  | **MCI-AD (n = 29)** | **Dementia-AD (n = 37)** | **P** |
| --- | --- | --- | --- |
| Diabetes mellitus, n (%) | 7 (24.14) | 4 (10.81) | 0.191 |
| Dyslipidemia, n (%) | 14 (48.28) | 17 (45.95) | 0.851 |
| Obesity, n (%) | 4 (13.79) | 5 (13.51) | 0.194 |
| Hypertension, n (%) | 16 (56.67) | 22 (59.46) | 0.727 |
| Psychiatric issues, n (%) | 9 (31.03) | 12 (32.43) | 1 |
| Plasma glucose (mg/dL) * | 104.94 (21.72) | 94.46 (18.73) | 0.042 |
| CSF glucose (mg/dL) # | 69.69 (13.59) | 60.61 (5.59) | 0.031 |
| Plasma Albumin (mg/mL) | 46.31 (4.81) | 45.43 (6.87) | 0.531 |
| CSF Albumin (mg/mL) | 0.095 (0.027) | 0.097 (0.032) | 0.872 |
| Qalb | 2.07 (0.59) | 2.18 (0.86) | 0.583 |

Differences between groups were assessed using Mann-Whitney U for continuous variables and chi-square test ( for dyslipidemia, hypertension and psychiatric issues) or Fisher’s exact test (for diabetes mellitus and obesity). Data are presented as mean (standard deviation) or number of participants (percentage), as appropriate. P values < 0.05 were considered statistically significant.

Note:

* Sample size differs due to incomplete data for the Glycemia parameter: NC, n=20; MCI-AD, n=17; Dementia-AD, n=28.

\# Sample size differs due to incomplete data for the CSF glucose parameter: NC, n=20, MCI-AD, n=16; Dementia-AD, n=28.

Abbreviations used: Qalb - CSF/plasma albumin quotient

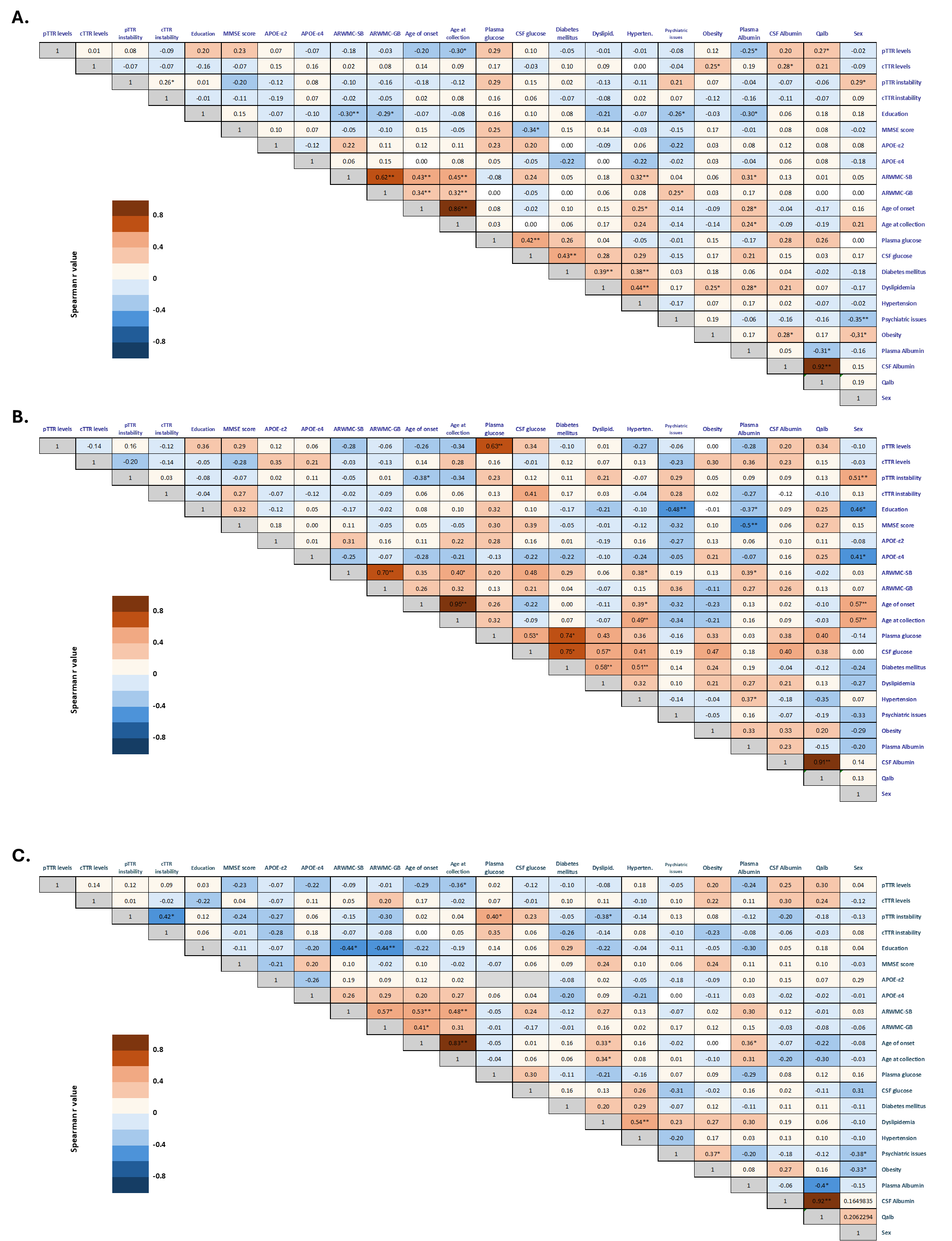

**Figure S1.** **Correlation between plasma (pTTR) and CSF TTR (cTTR) instability and levels with demographic and clinical variables in** **A.** MCI-AD and Dementia-AD (n=66) group; **B.** MCI-AD (n=29) group; **C.** Dementia-AD (n=37) group. Sample size differs due to incomplete data for the plasma glucose parameter (MCI-AD and Dementia-AD, n=45; MCI-AD, n=17; Dementia-AD, n=28) and CSF glucose parameter (MCI-AD and Dementia-AD, n=44; MCI-AD, n=16; Dementia-AD, n=28).

The correlation matrix graphically represents the Spearman's correlation coefficient (r) for each pairwise comparison among the biomarkers. According to the color scale on the right side of the matrix, positive and negative correlations are indicated in shades of red and blue, respectively an r value of +1 indicates a perfect positive relationship, r = 0 indicates no relationship, and an r value of -1 indicates a perfect negative relationship. Significant differences are denoted by * (p < 0.05) and ** (p < 0.01).

Abbreviations used: APOE - apolipoprotein E, ARWMC - age-related white matter changes, CSF - cerebrospinal fluid, MCI - mild cognitive impairment, MMSE - Mini-Mental State Examination, Qalb - CSF/plasma albumin quotient
