## Supplementary material for "Transthyretin Levels and Instability in Alzheimer’s Disease: Correlations with AD Biomarkers in a Cohort Study": Uncroped Blots

1. **Evaluation of plasma instability**

All plasma samples were analyzed in two independent sets of Western blot experiments, with one replicate per sample. A total of 12 gels were run for each set, and the results were visualized in six images, each showing two gels. The arrangement of the samples in each gel is detailed in a corresponding template displayed above the respective blot. For each gel, a molecular weight marker (MW marker) and recombinant TTR (recTTR) were included as controls. Each gel contains eight plasma samples from patients with Mild Cognitive Impairment-AD (MCI-AD) or AD Dementia (Dem.-AD). Some blots also include non-demented Control (NC) samples, which were not analyzed within the scope of this study. Monomers and dimers are represented in the images, and the ratio of the band intensities of monomers to dimers was used to evaluate plasma instability. All images were acquired with a 1 second exposure time. The general scheme of the blots is as follow:

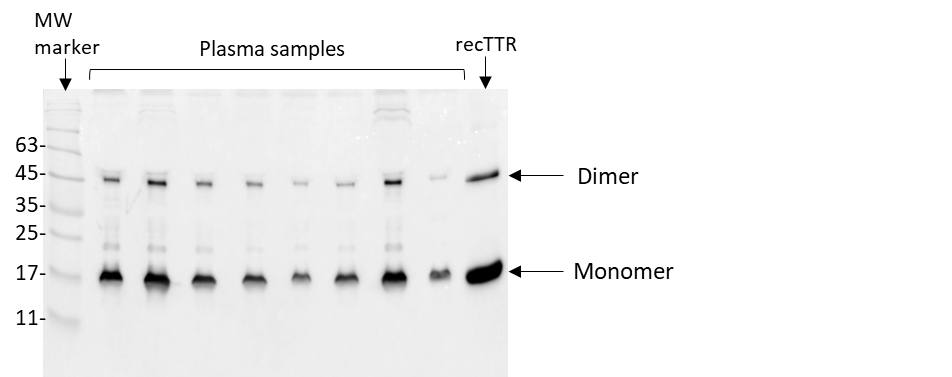

**1.1. Evaluation of plasma instability – 1st set of blots (12 blots)**

**Blot 1 and 2**

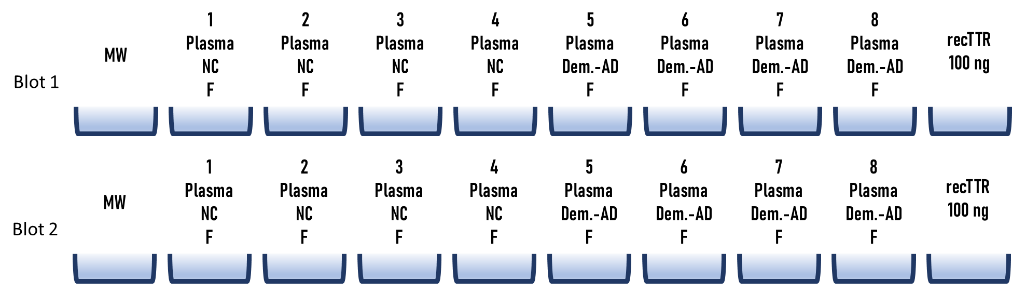

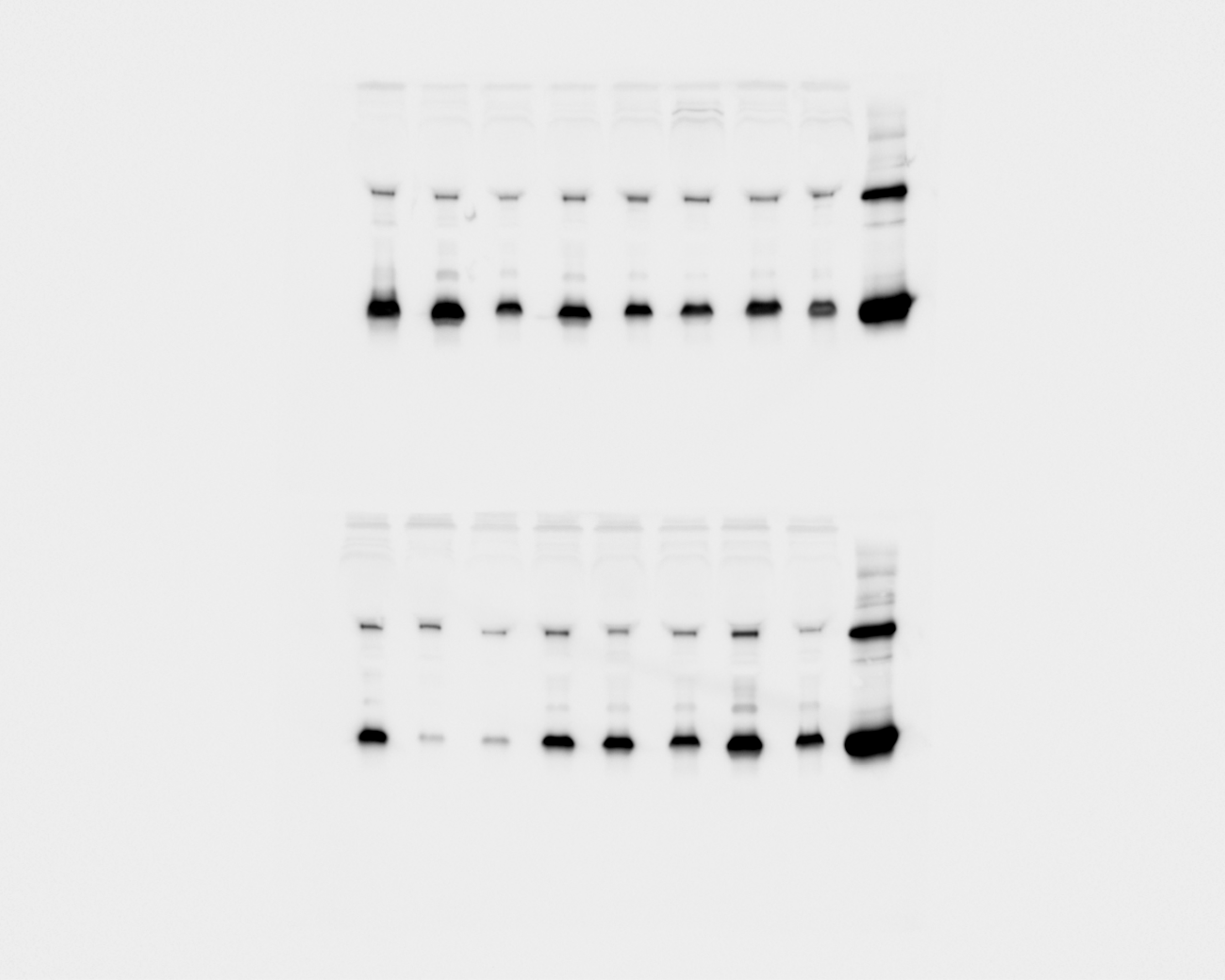

**Blot 3 and 4**

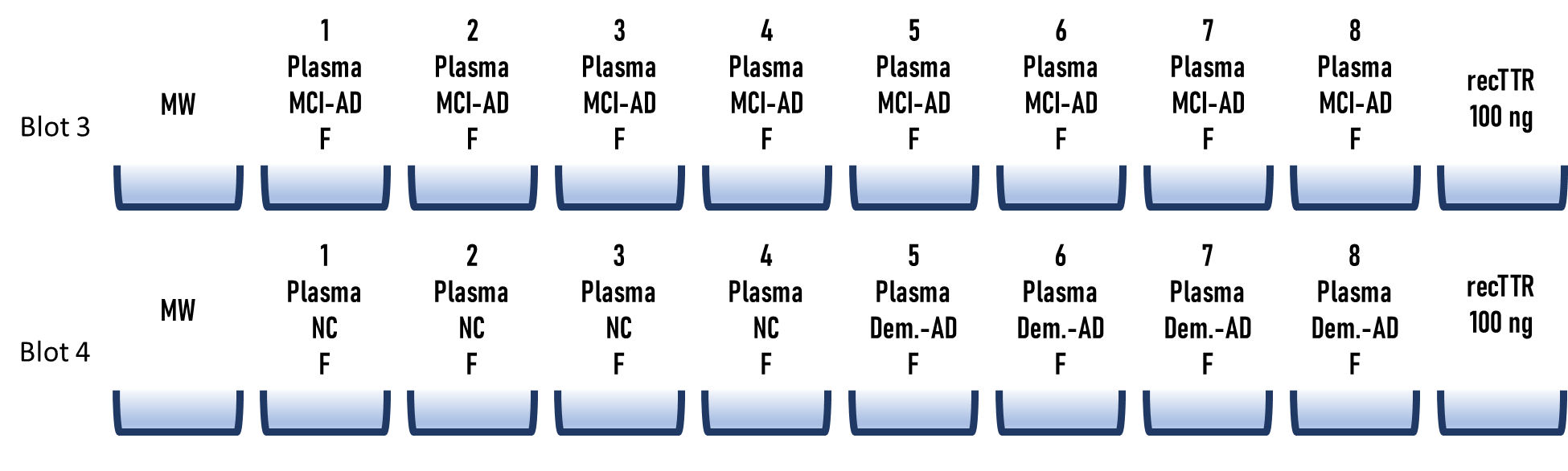

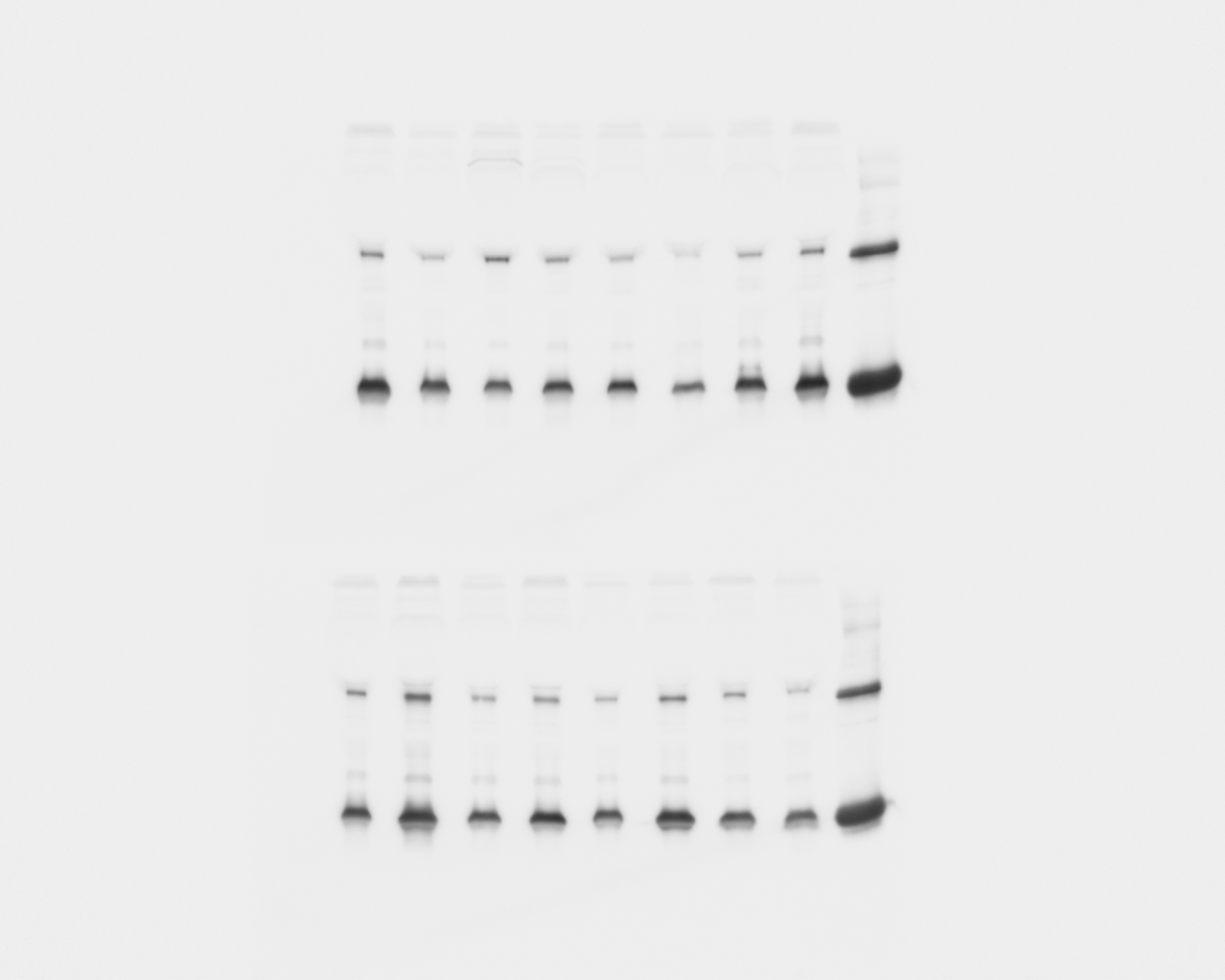

**Blot 5 and 6**

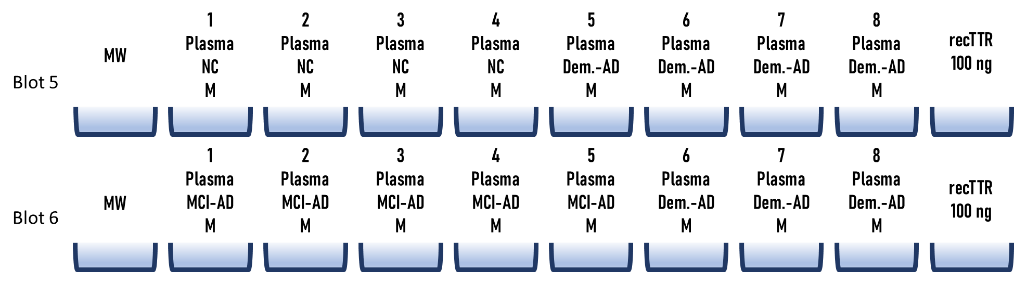

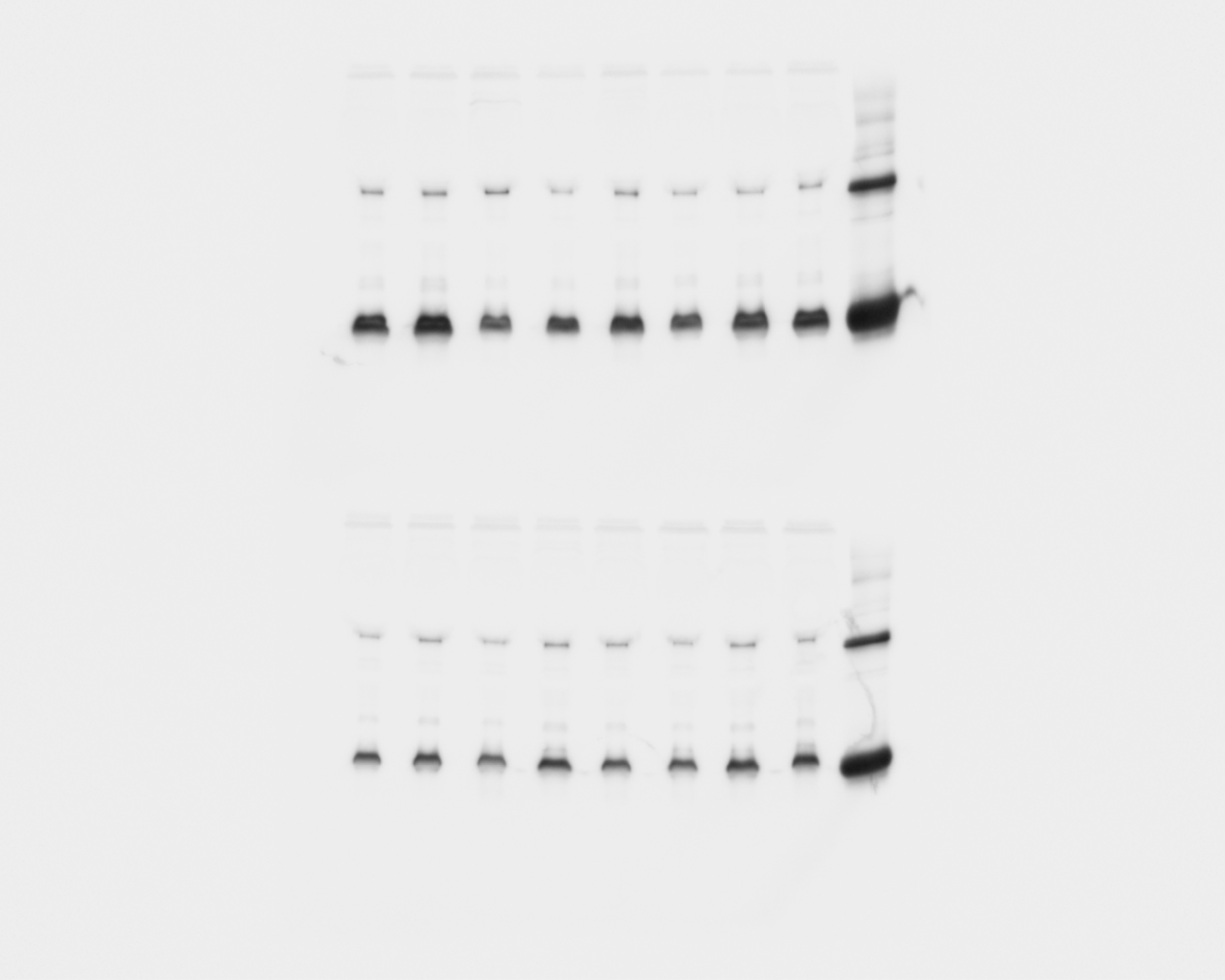

**Blot 7 and 8**

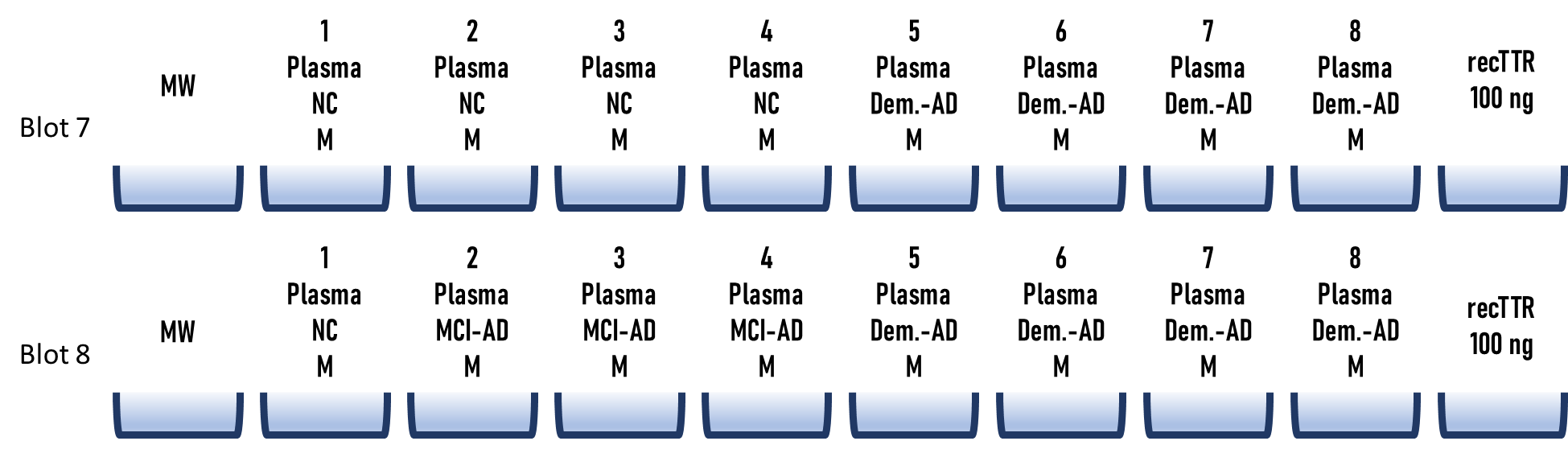

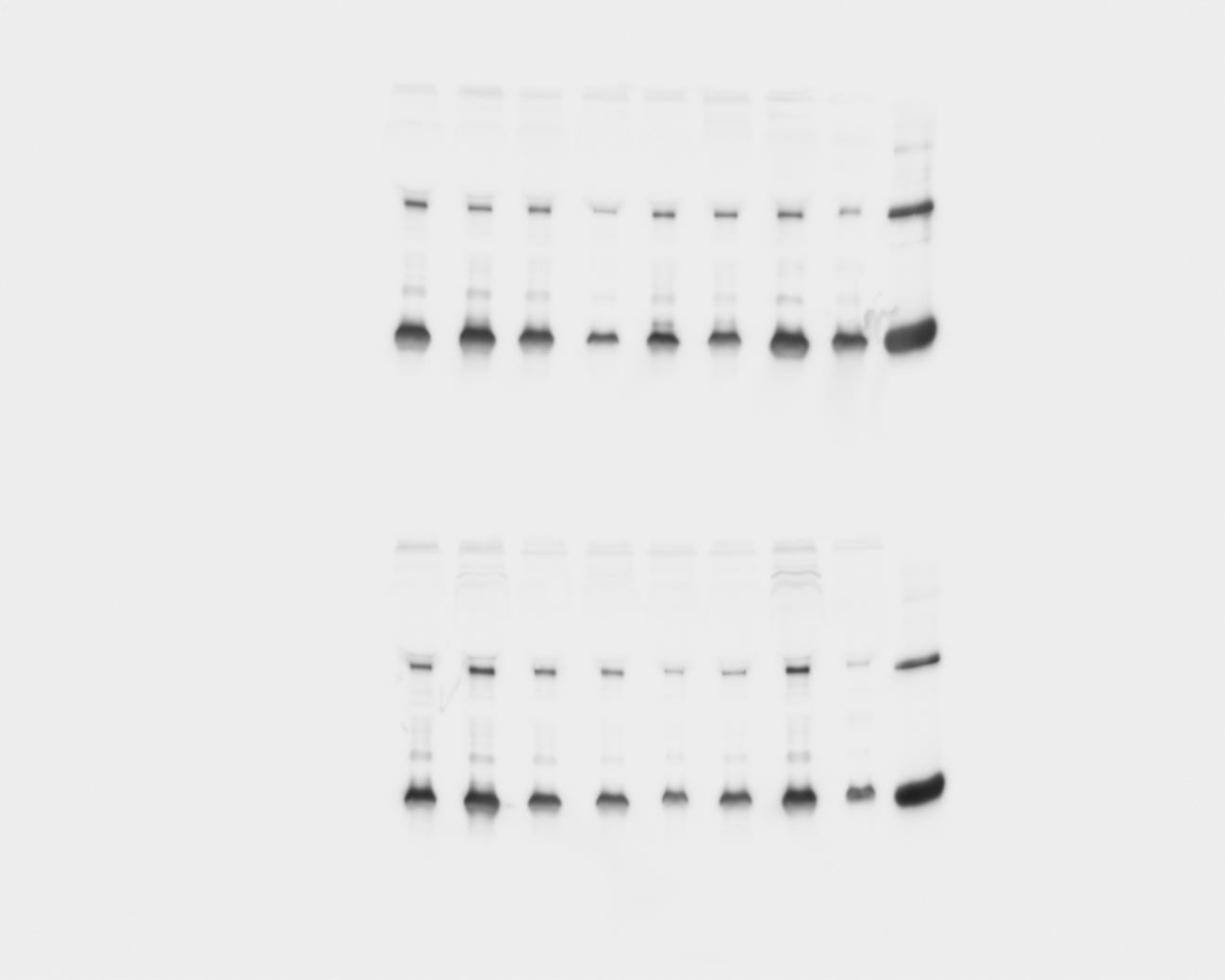

**Blot 9 and 10**

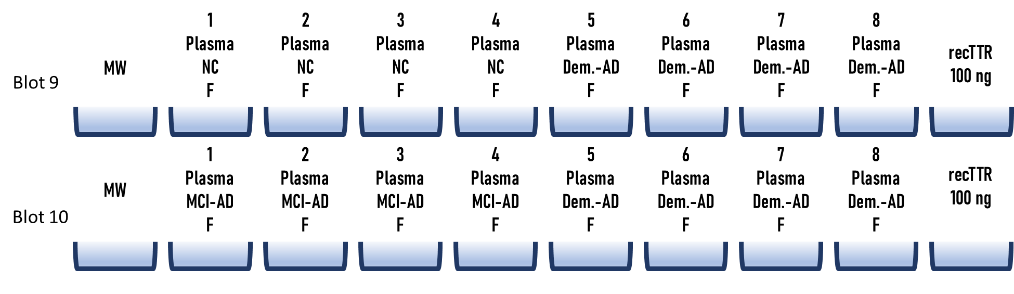

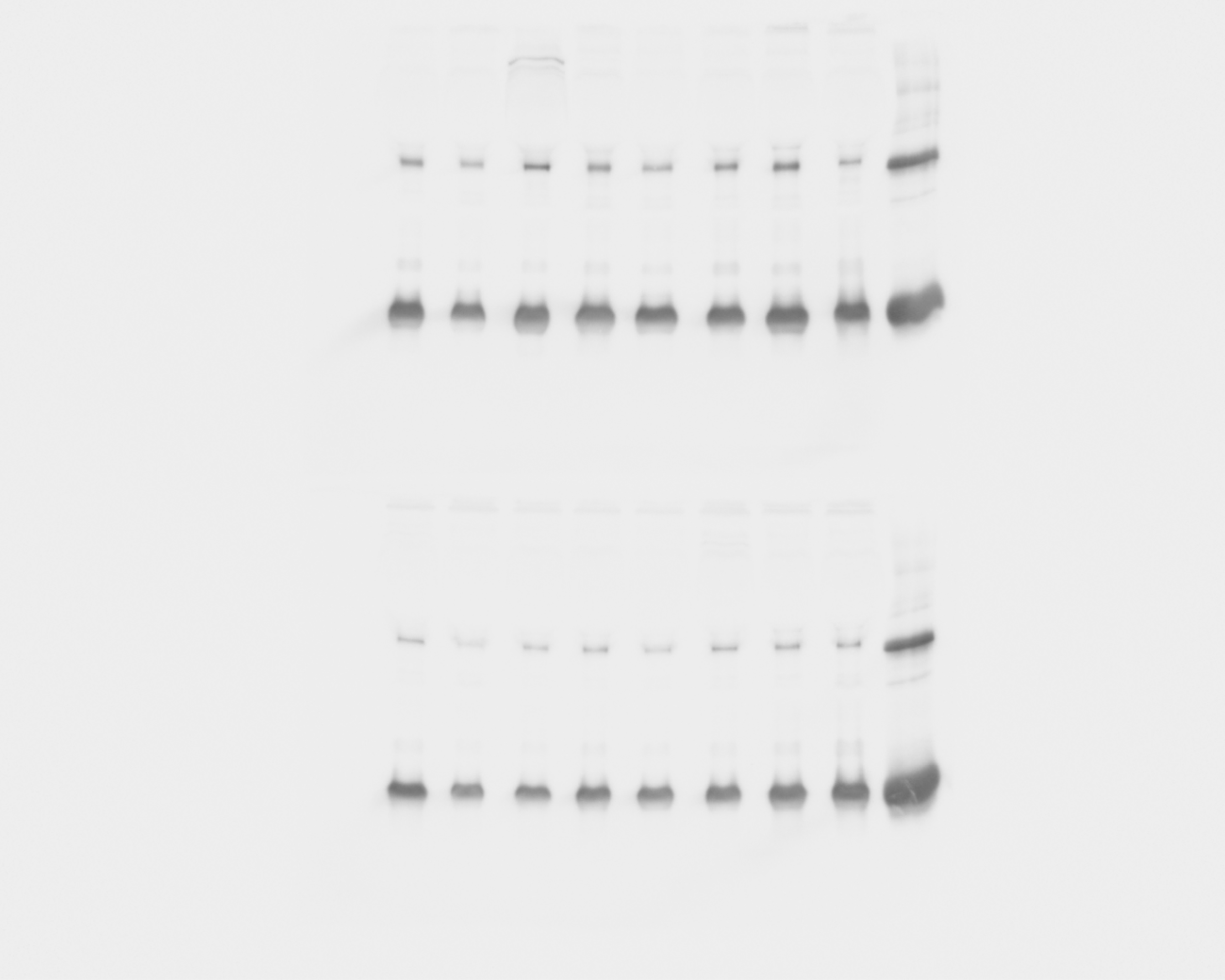

**Blot 11 and 12**

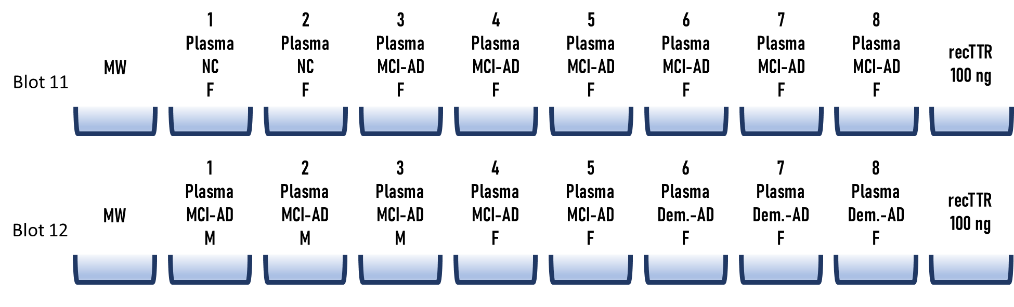

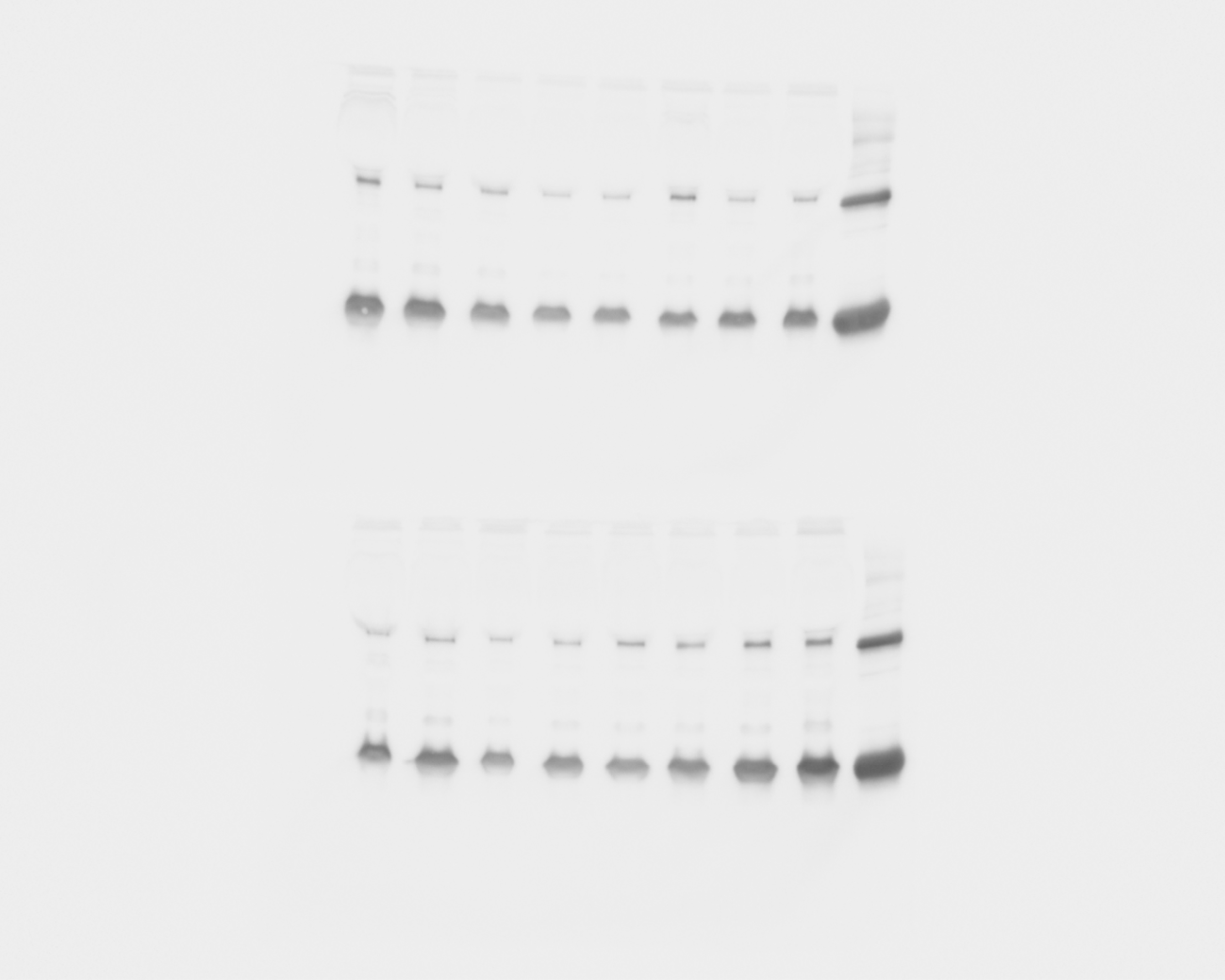

**1.2. Evaluation plasma instability – 2nd set of blots (12 blots)**

**Blot 1 and 2**

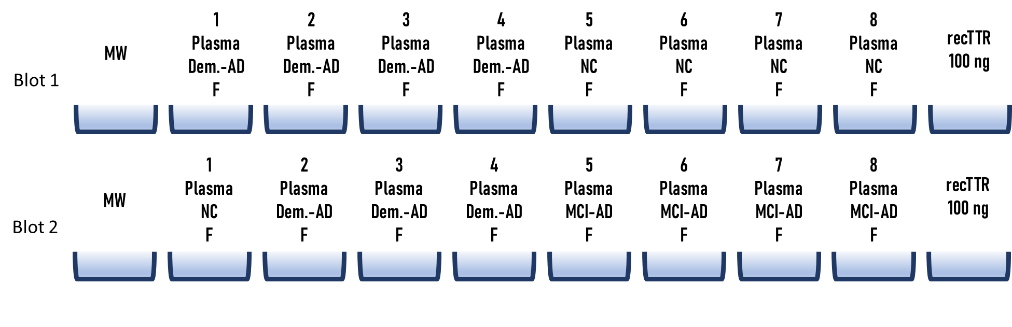

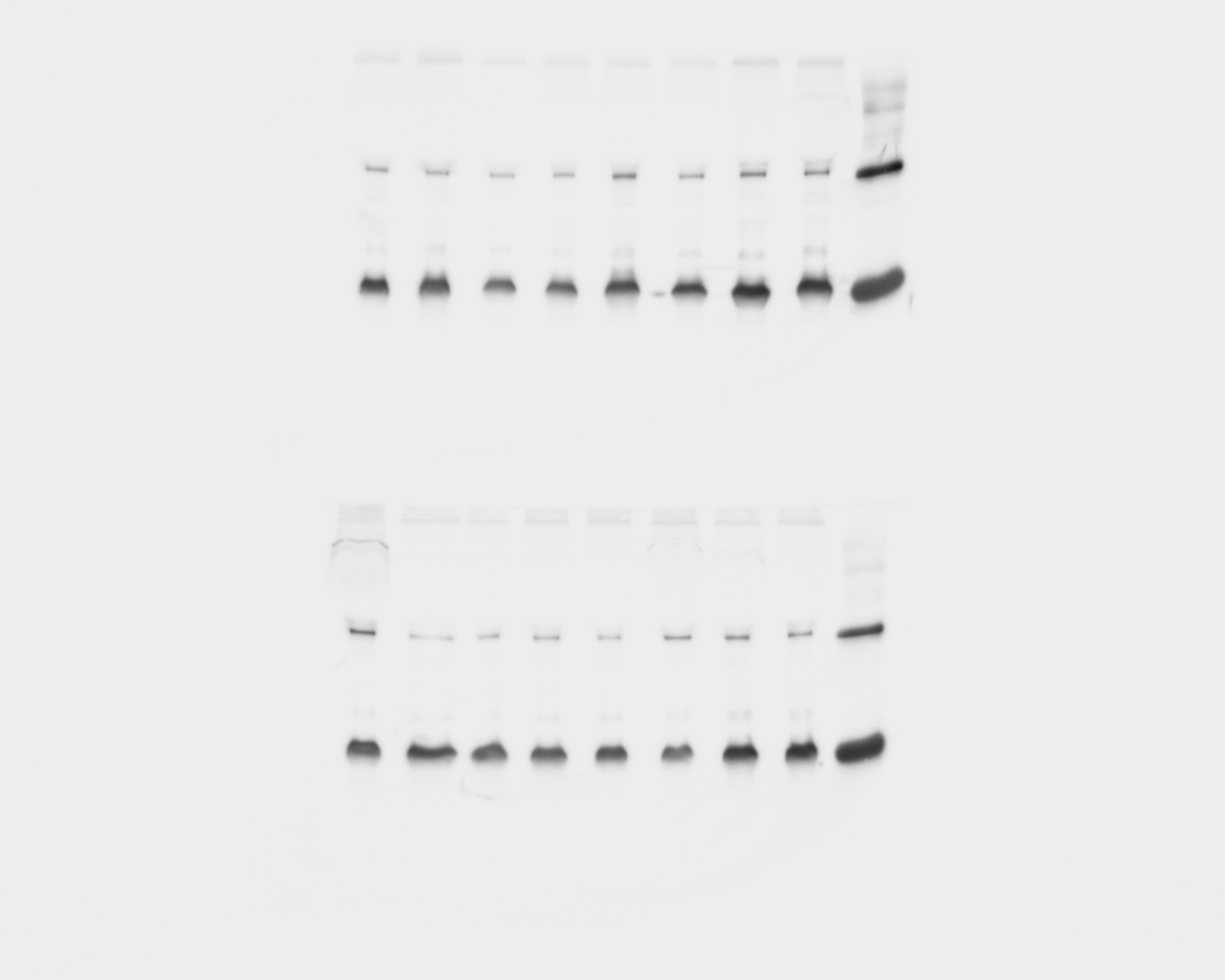

**Blot 3 and 4**

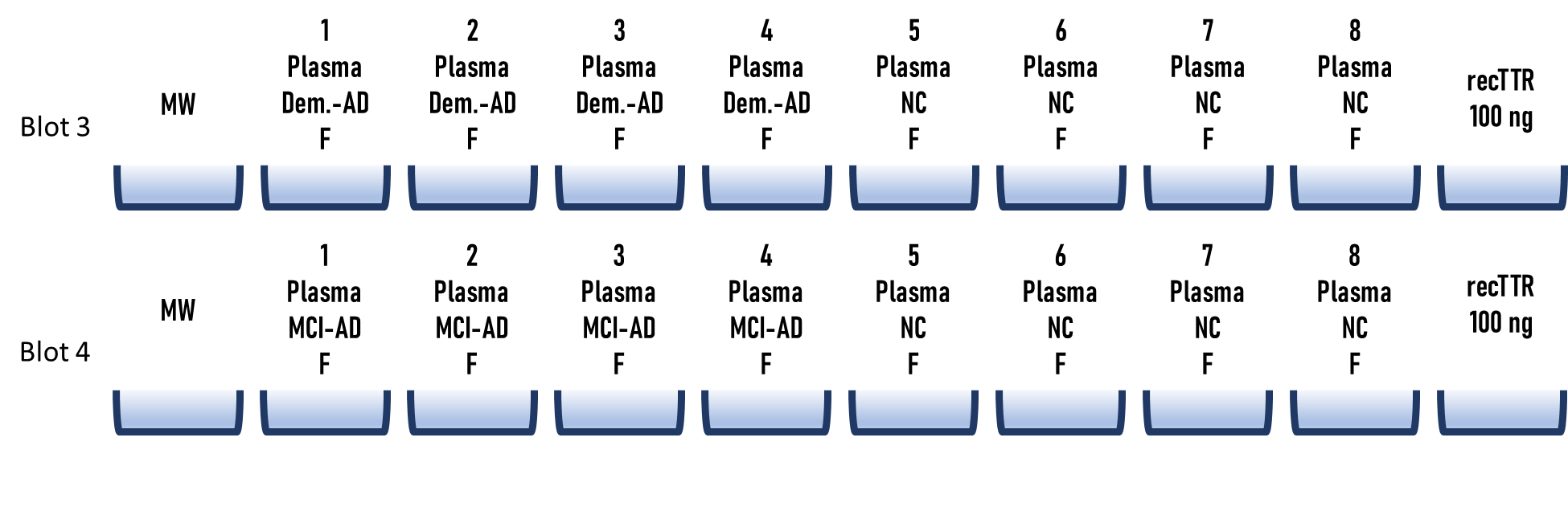

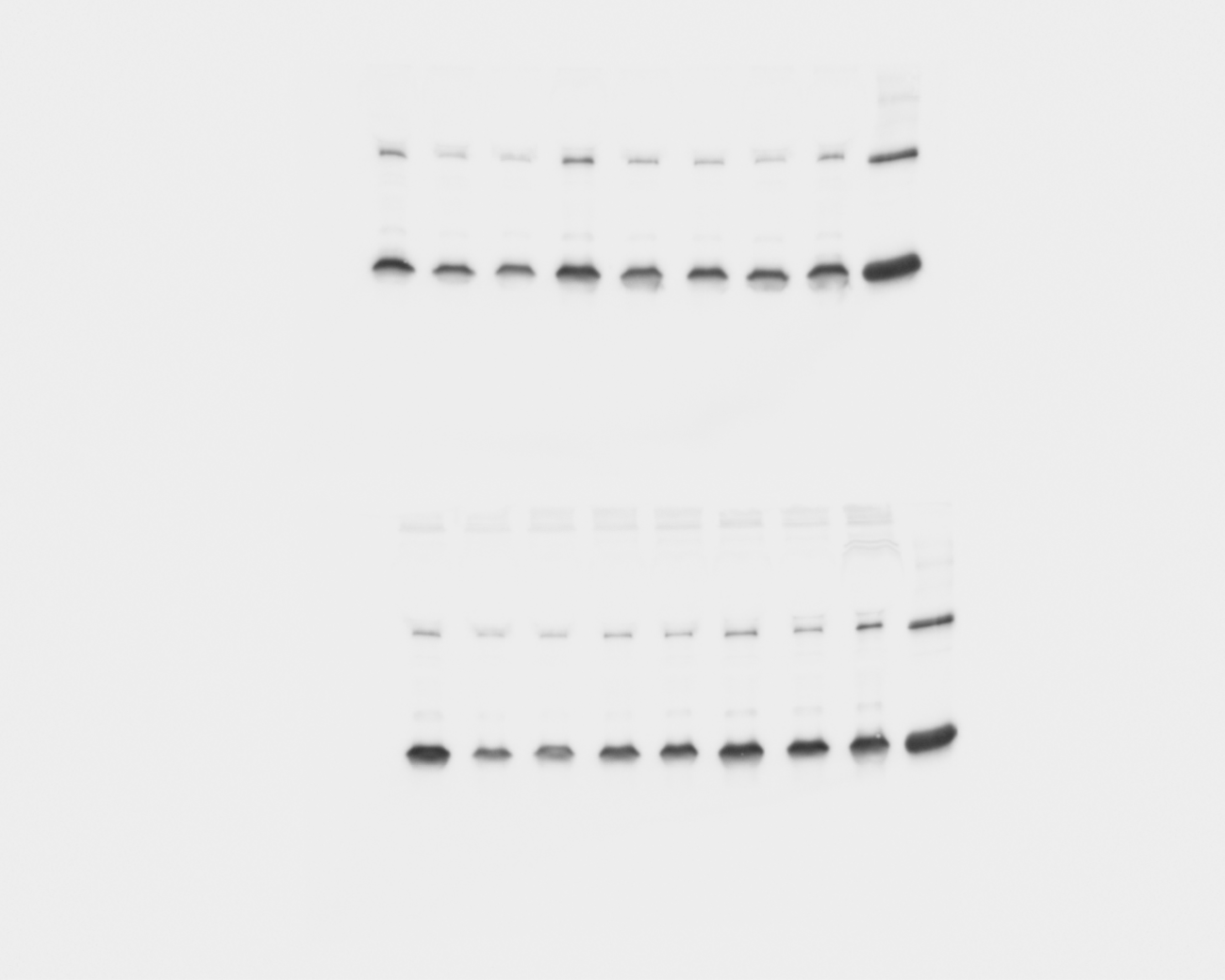

**Blot 5 and 6**

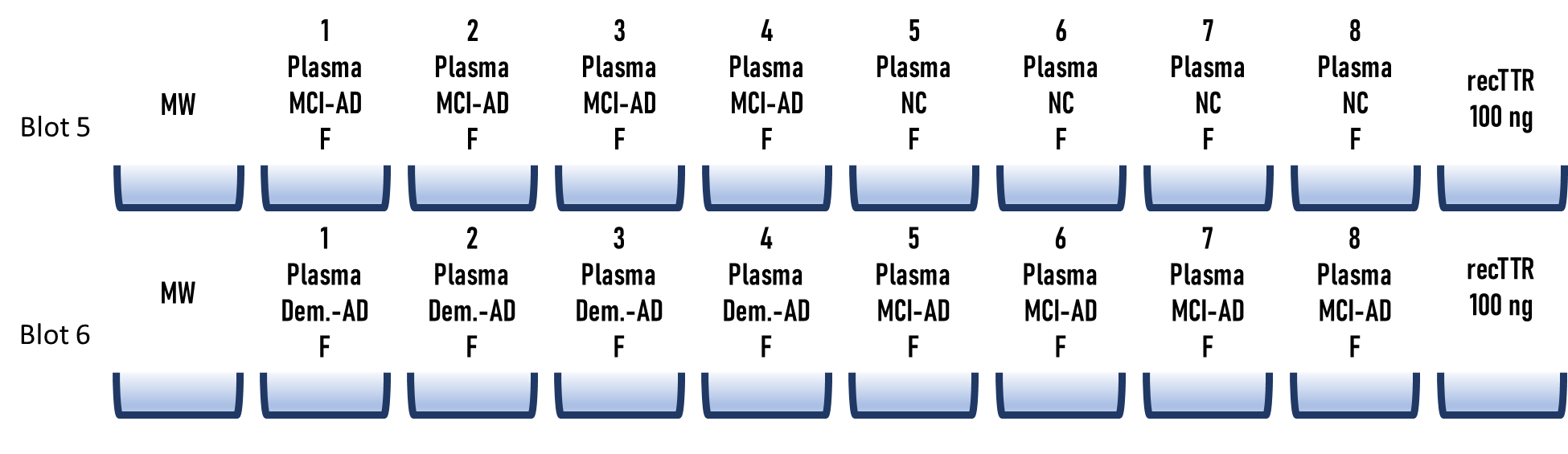

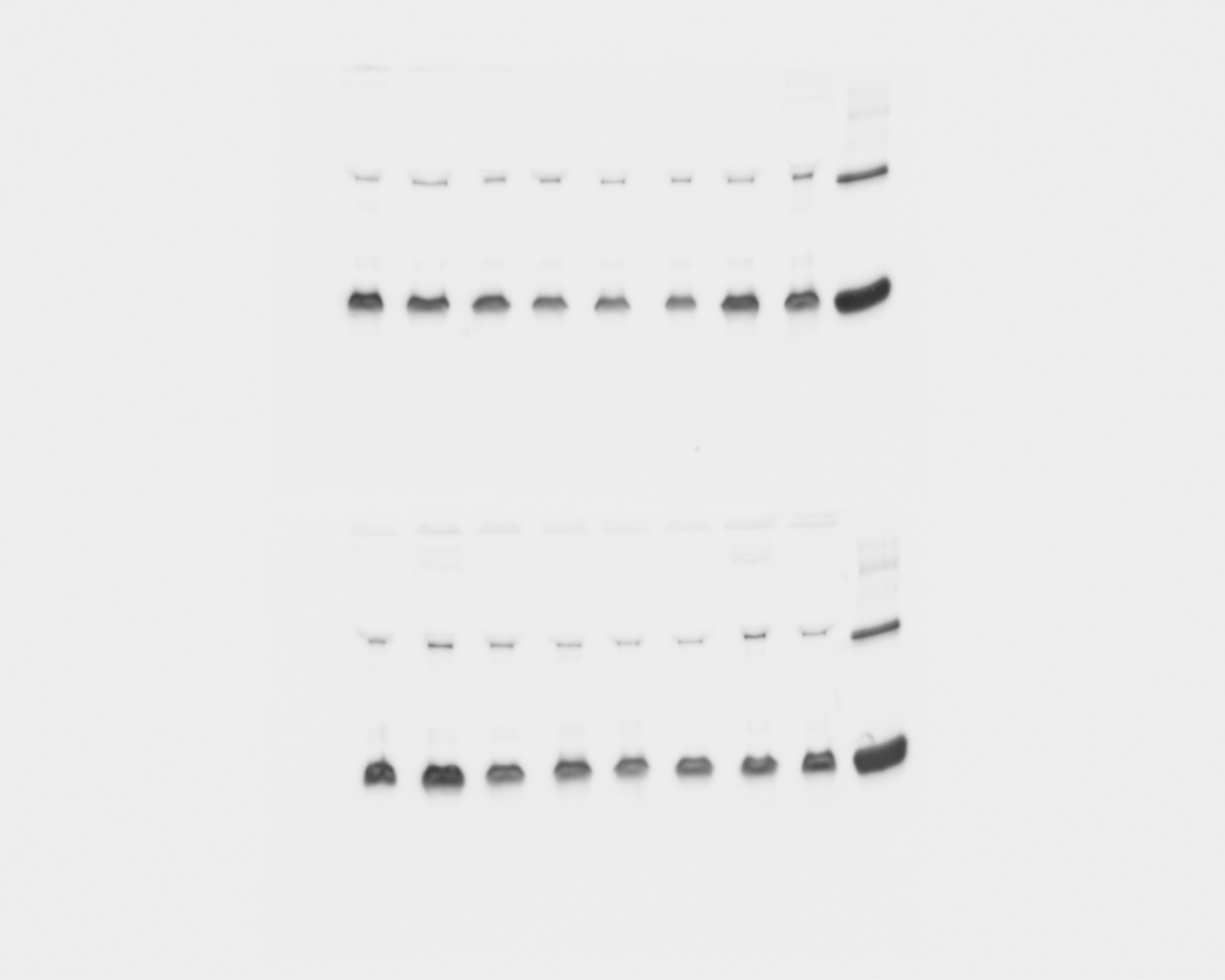

**Blot 7 and 8**

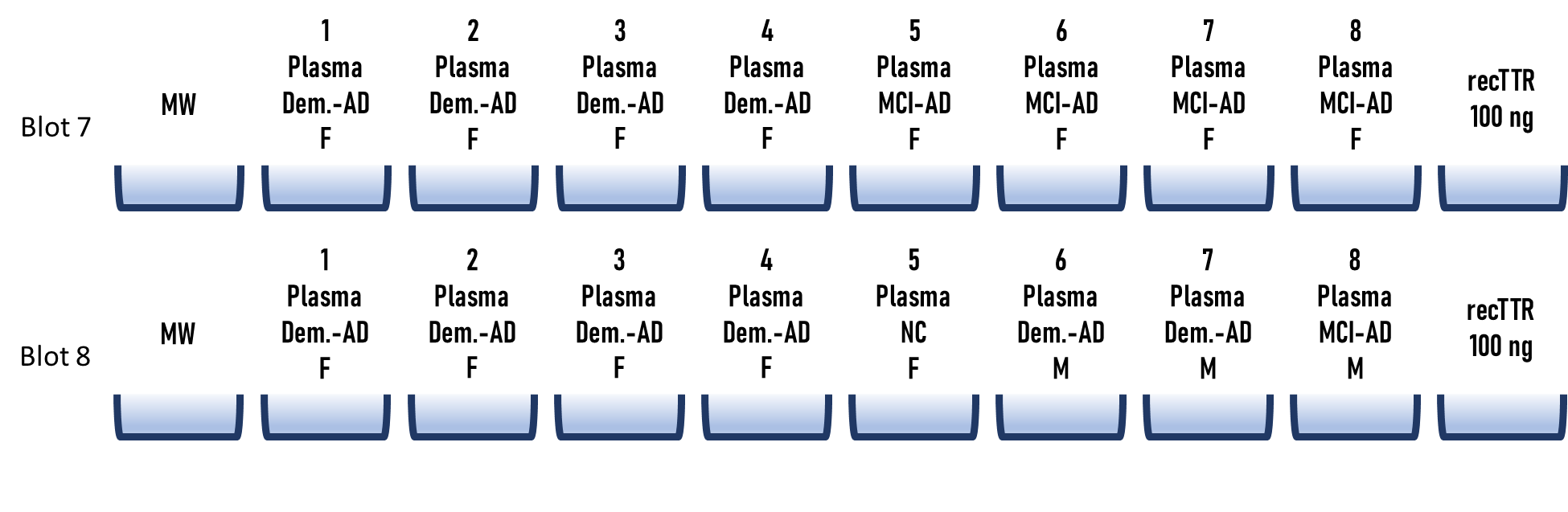

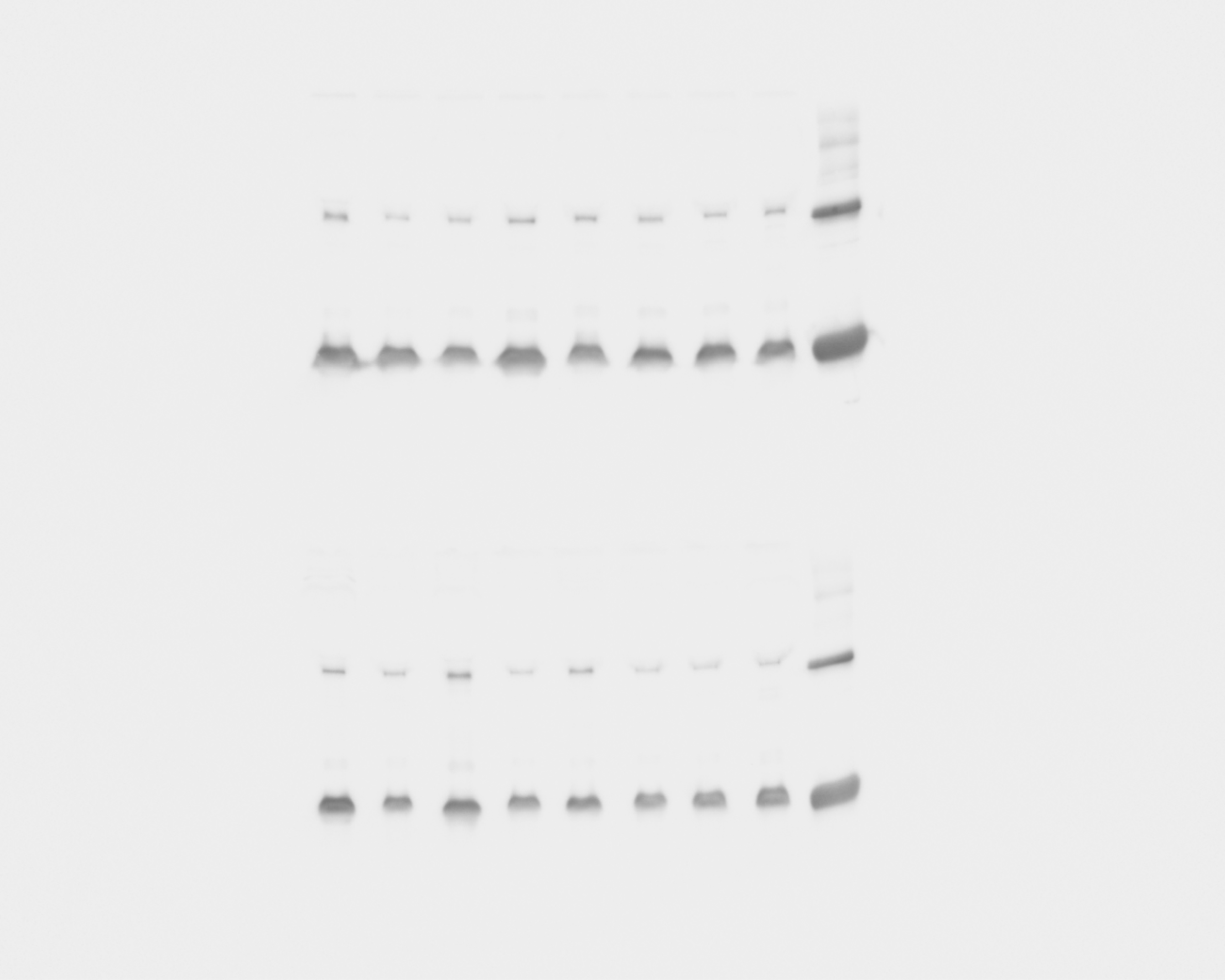

**Blot 9 and 10**

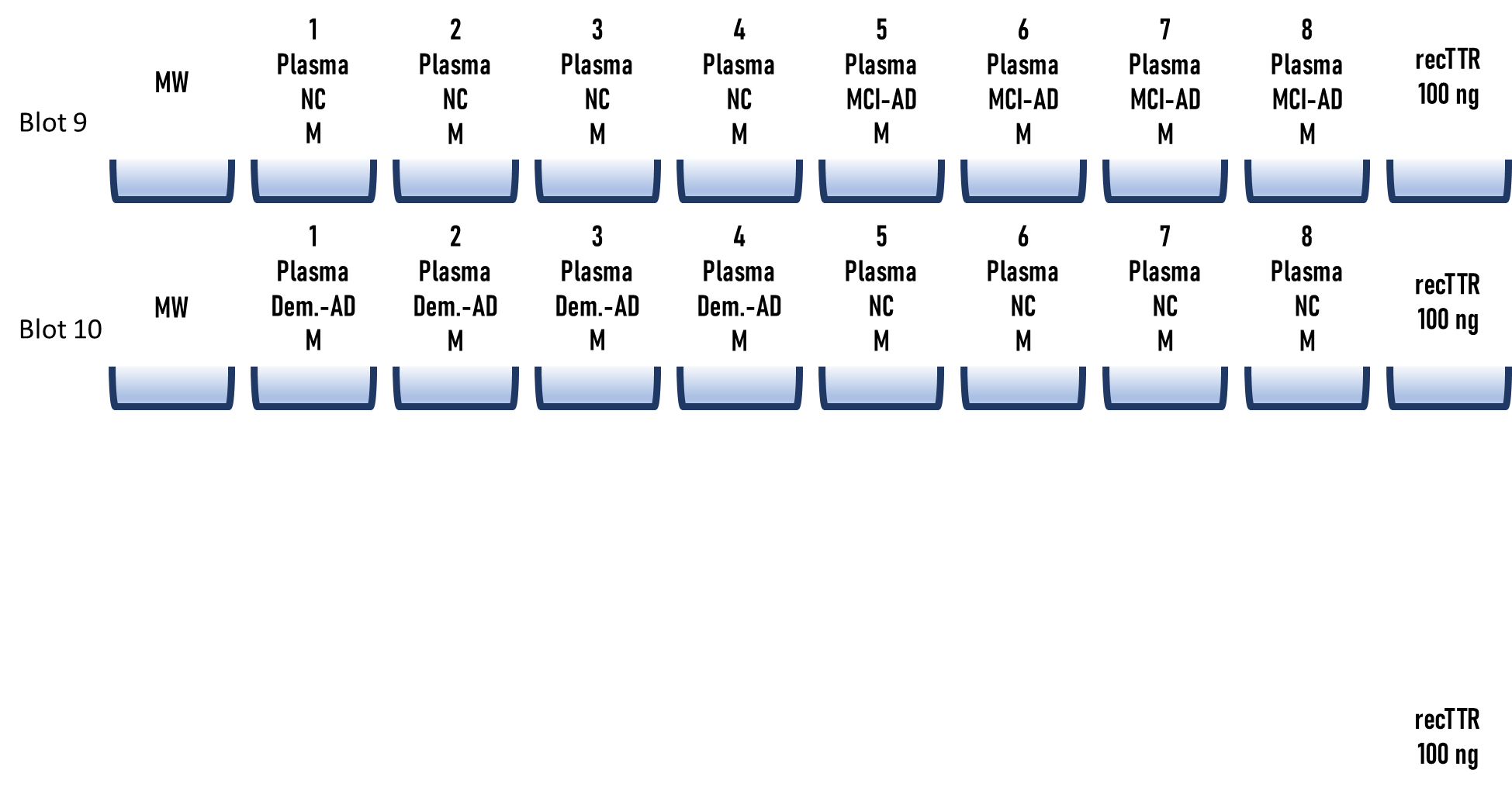

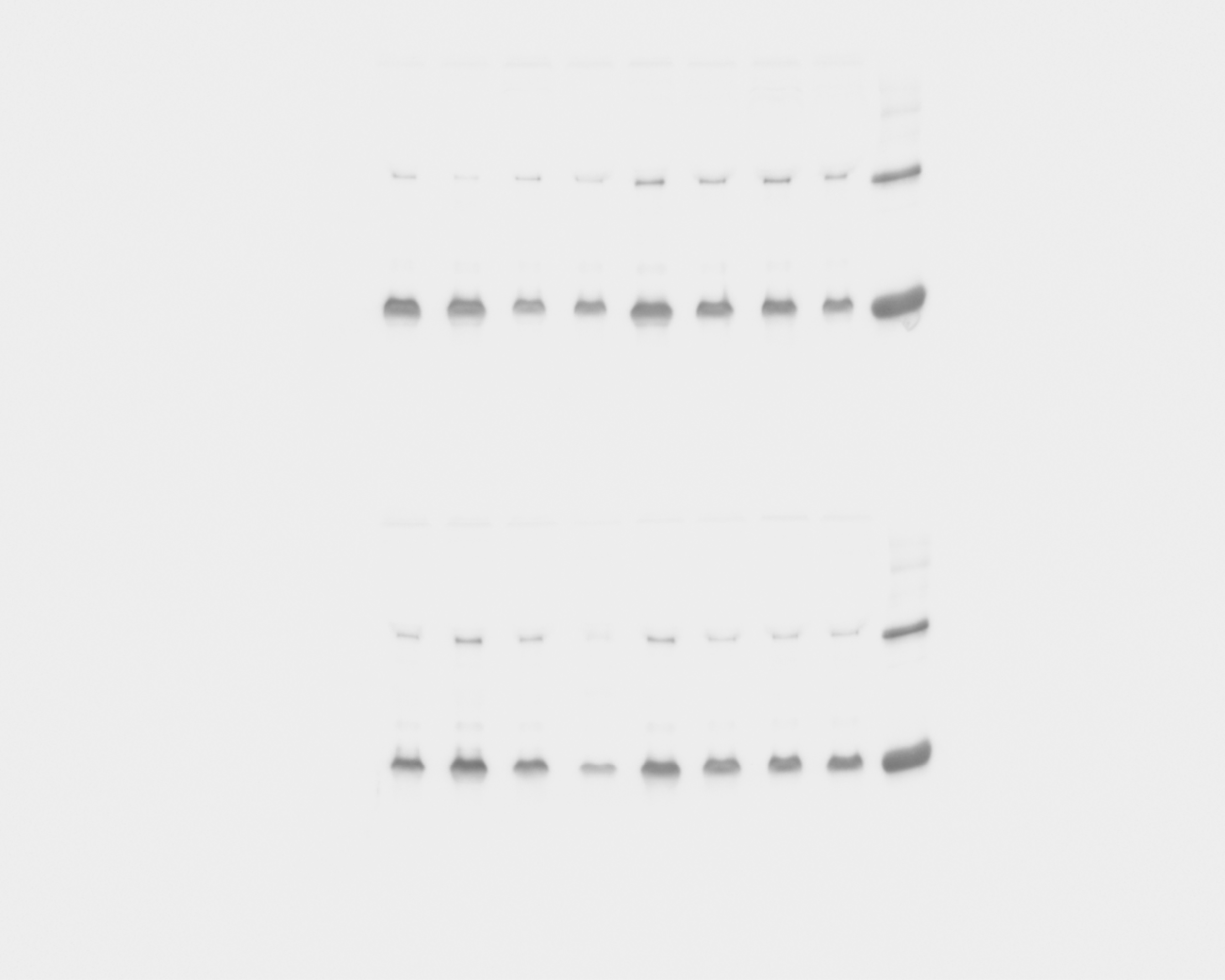

**Blot 11 and 12**

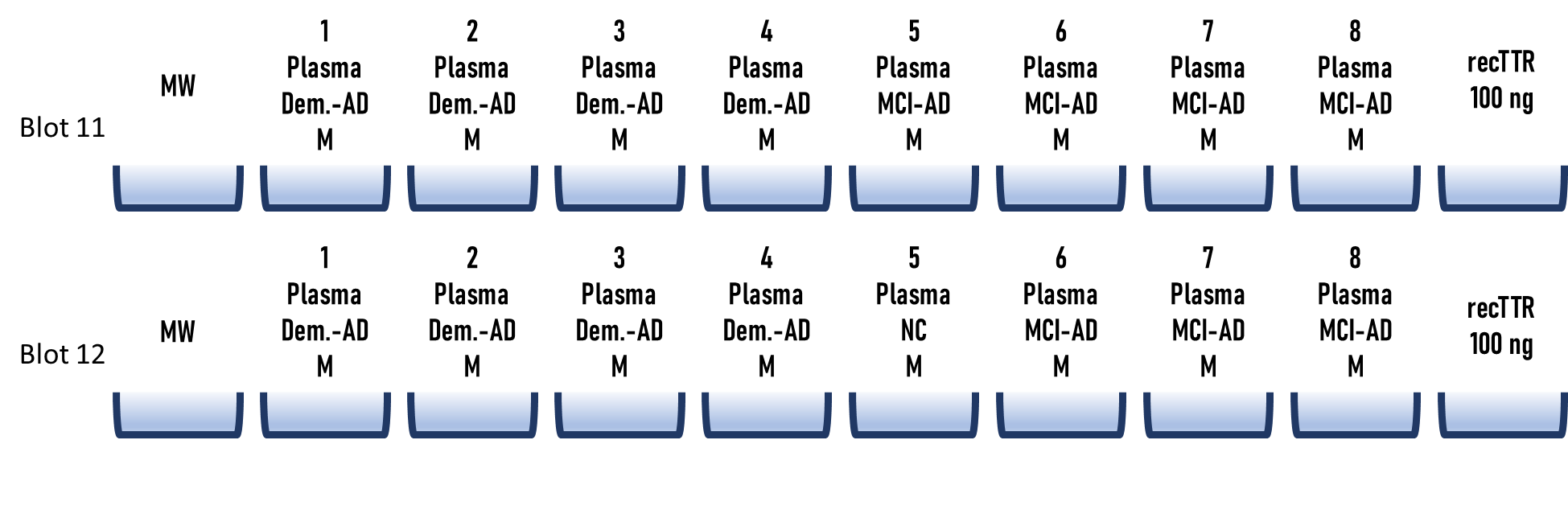

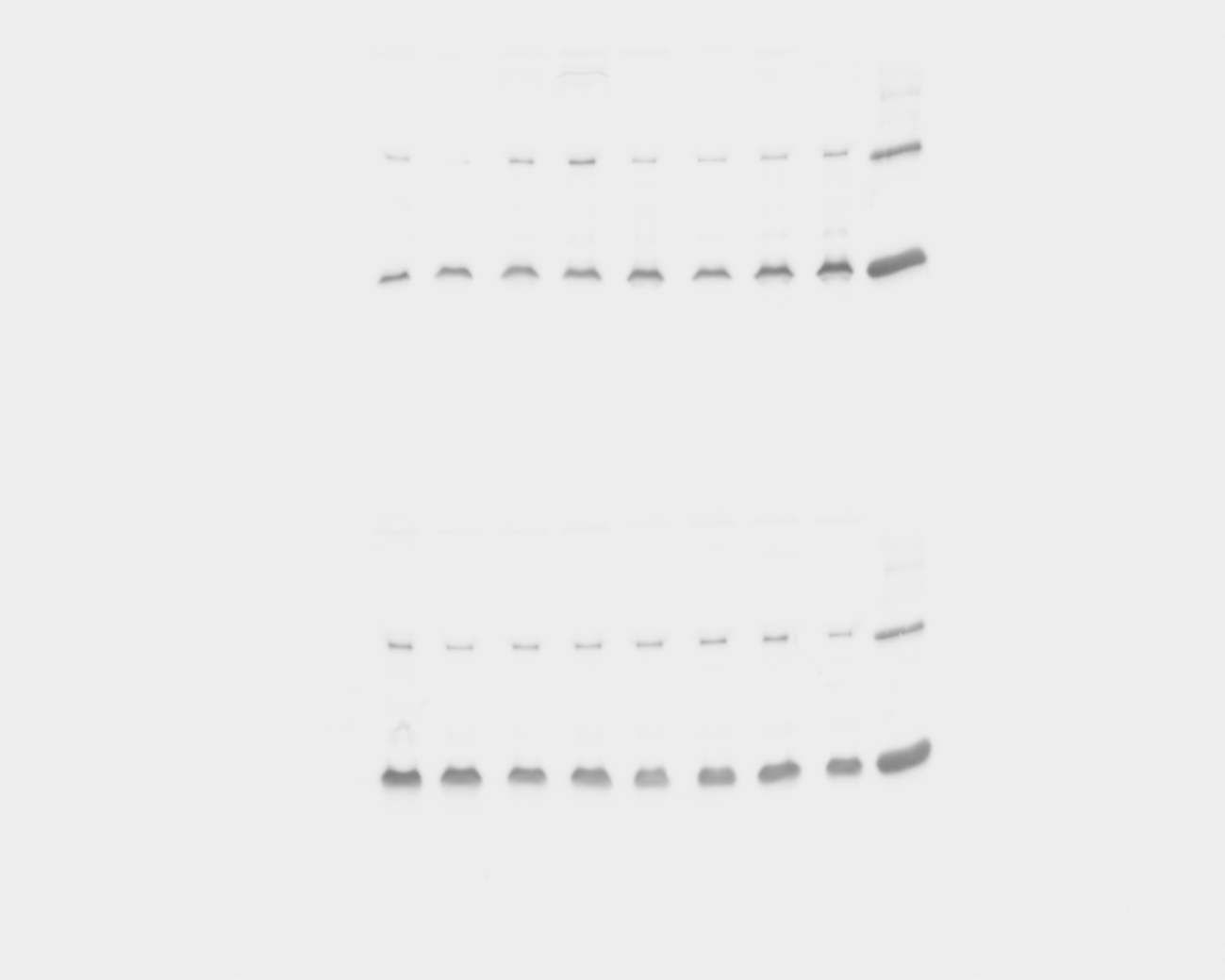

2. **Evaluation of CSF instability**

All CSF samples were analyzed in two independent sets of Western blot experiments, with one replicate per sample. A total of twelve gels were run for each set, and the results were visualized in six images, each showing two gels. The arrangement of the samples in each gel is detailed in a corresponding template displayed above the respective blot. For each gel, a molecular weight marker (MW marker) and recombinant TTR (recTTR) were included as controls. Each gel contains eight CSF samples from patients with Mild Cognitive Impairment -AD (MCI-AD) or Dementia-AD (Dem.-AD). Some blots also include non-demented control (NC) samples, which were not analyzed within the scope of this study. Monomers and dimers are represented in the images. Monomers were quantified using images acquired with a 1 second exposure time, while dimers were quantified using images acquired with an 11 second exposure time. The ratio of the band intensities of monomers to dimers was used to evaluate CSF instability. The general scheme of the blots is as follow:

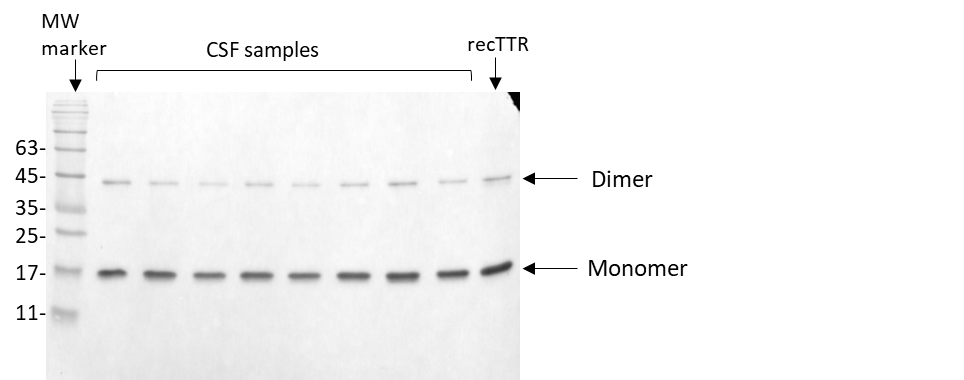

**2.1. Evaluation of CSF instability – 1st set of blots (12 blots)**

**Blot 1 and 2**

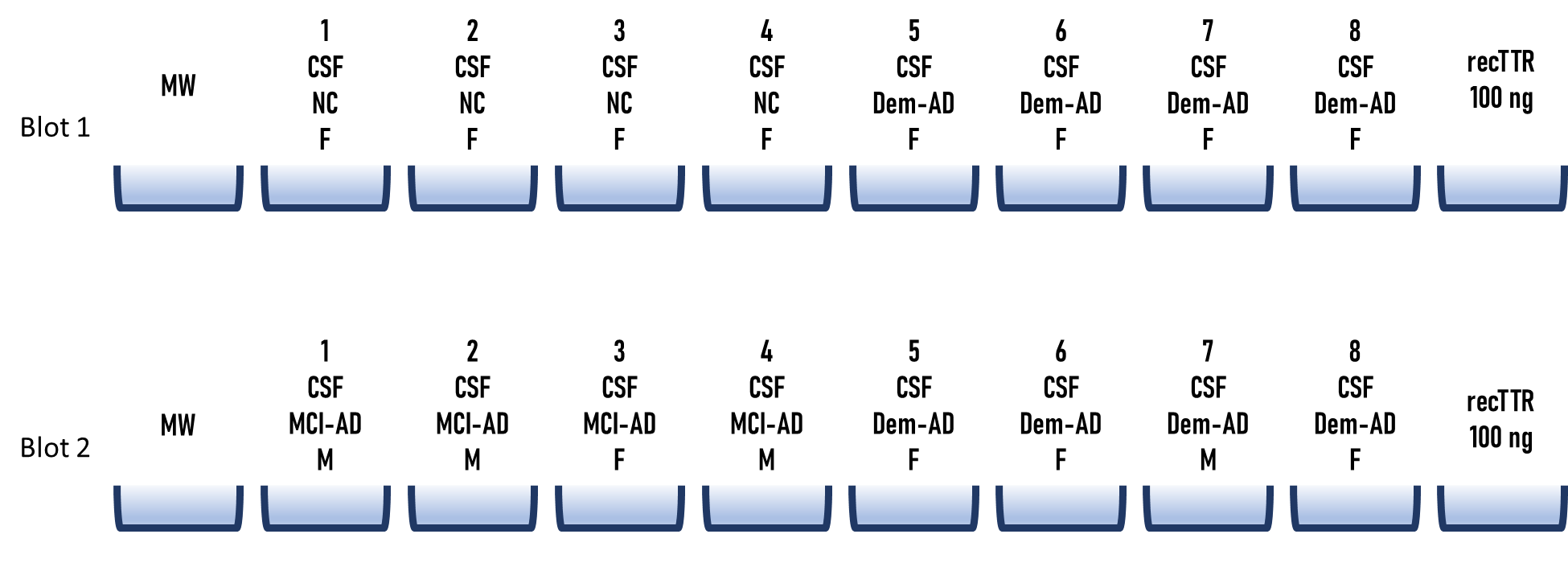

Exposition time = 1 second

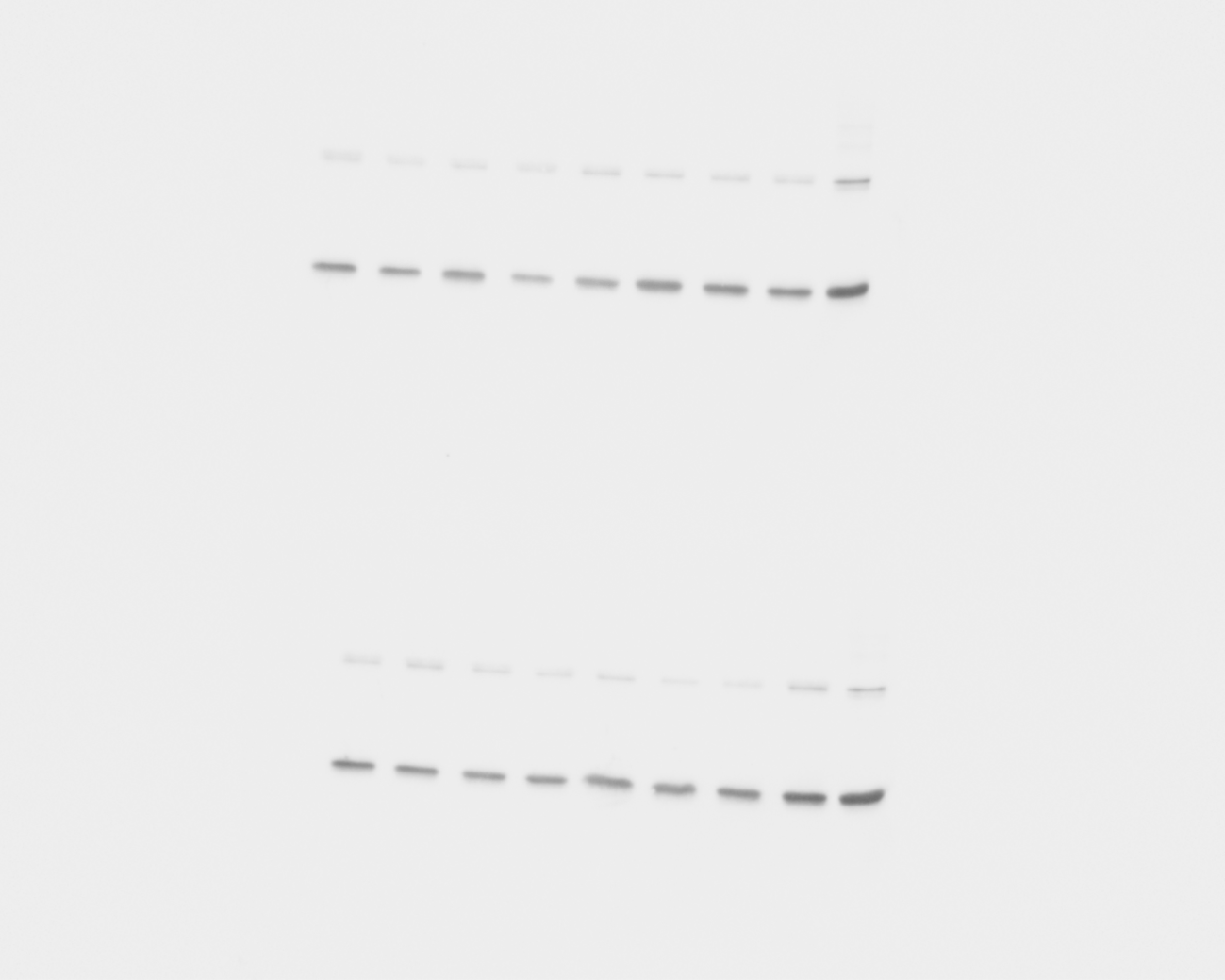

Exposition time = 11 seconds

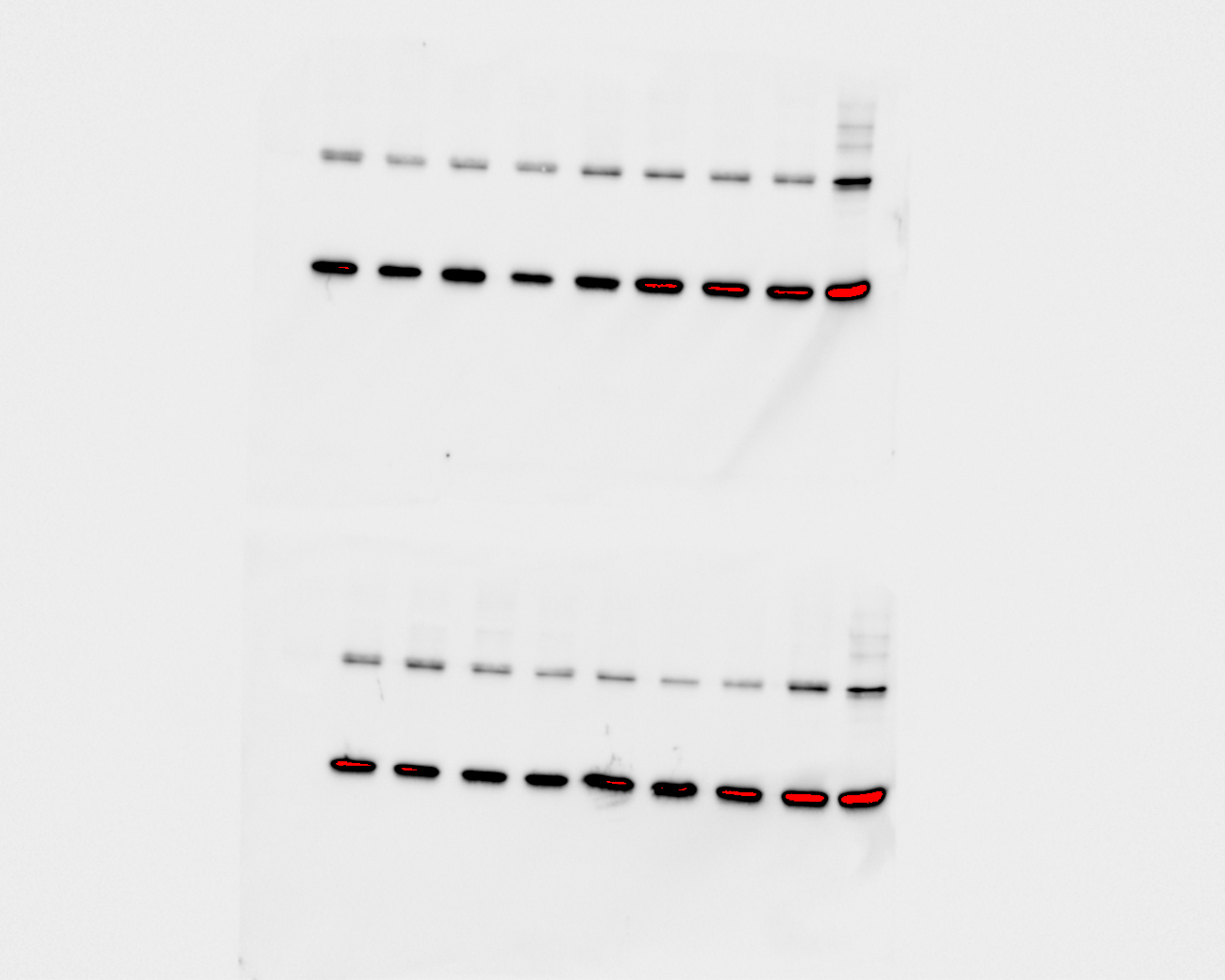

**Blot 3 and 4**

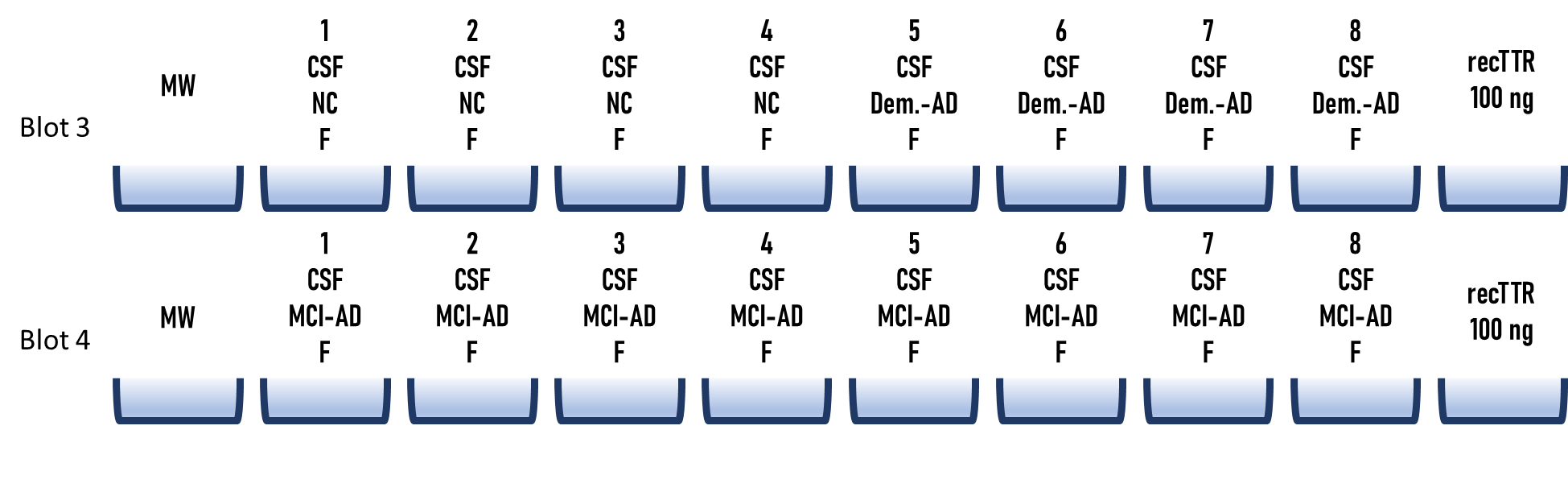

Exposition time = 1 second

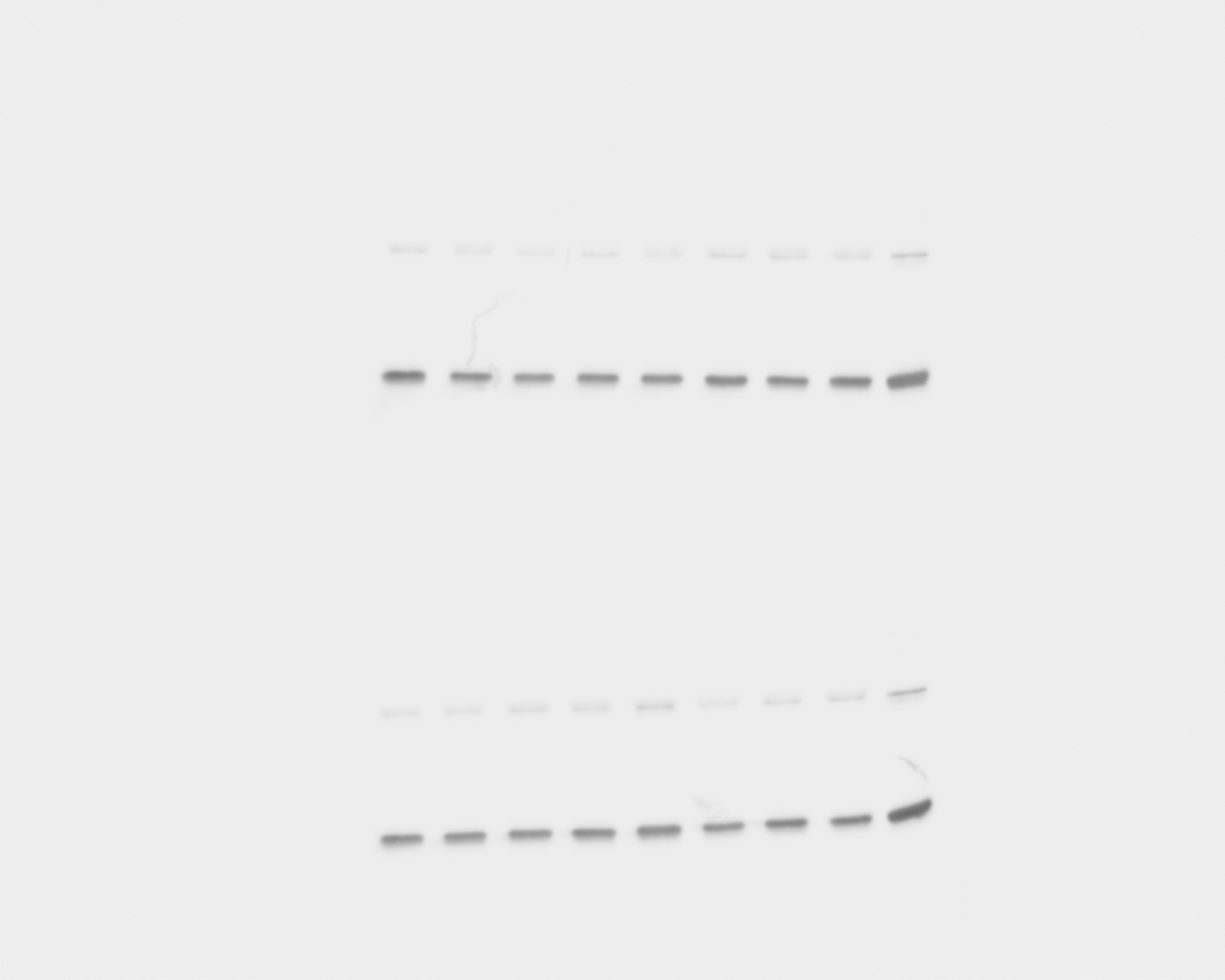

Exposition time = 11 seconds

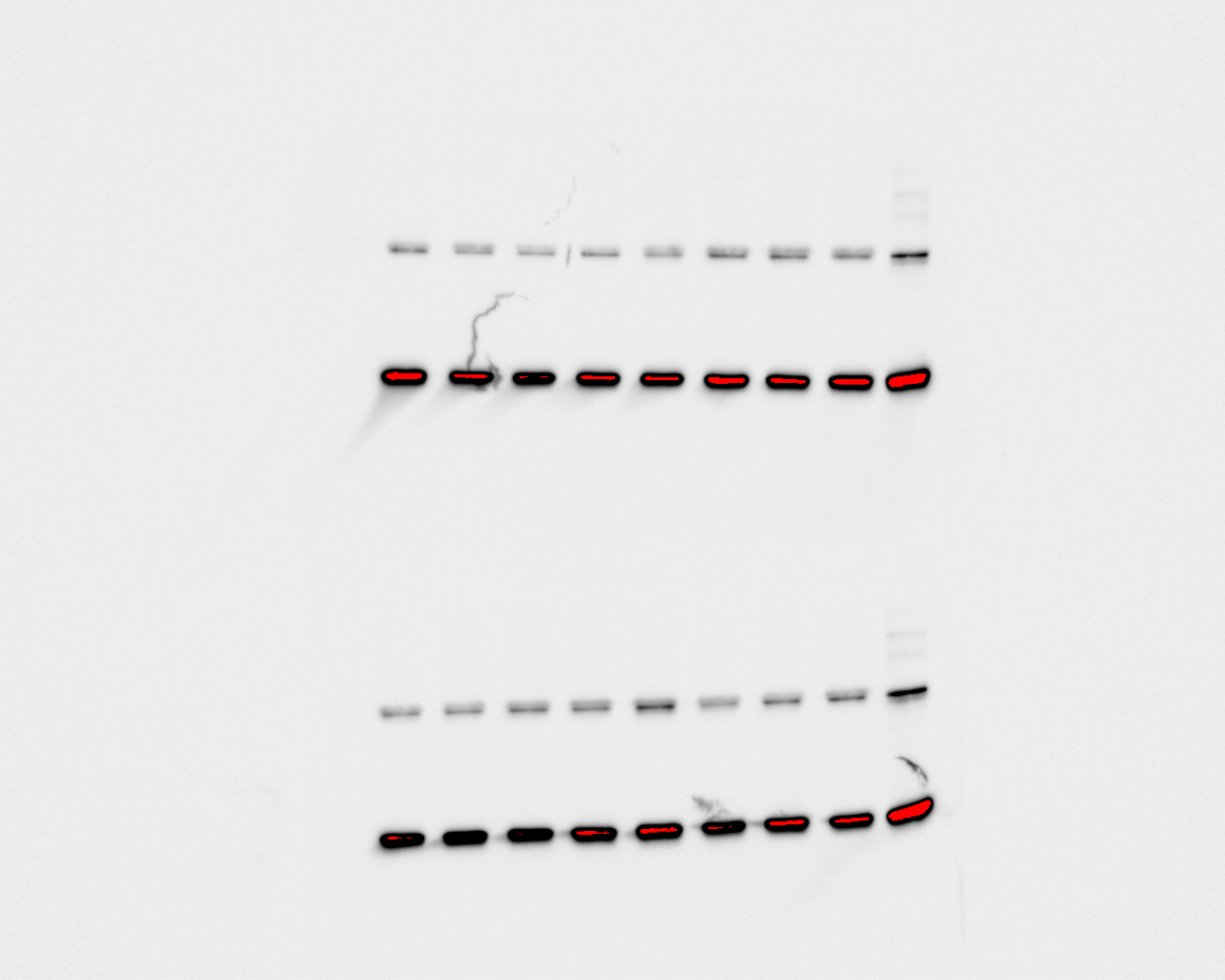

**Blot 5 and 6**

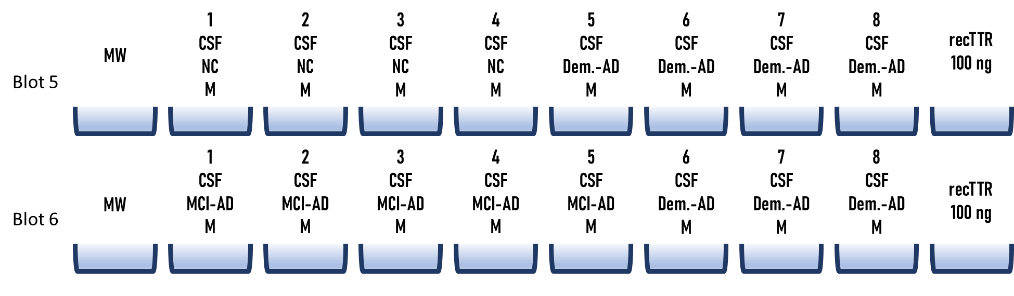

Exposition time = 1 second

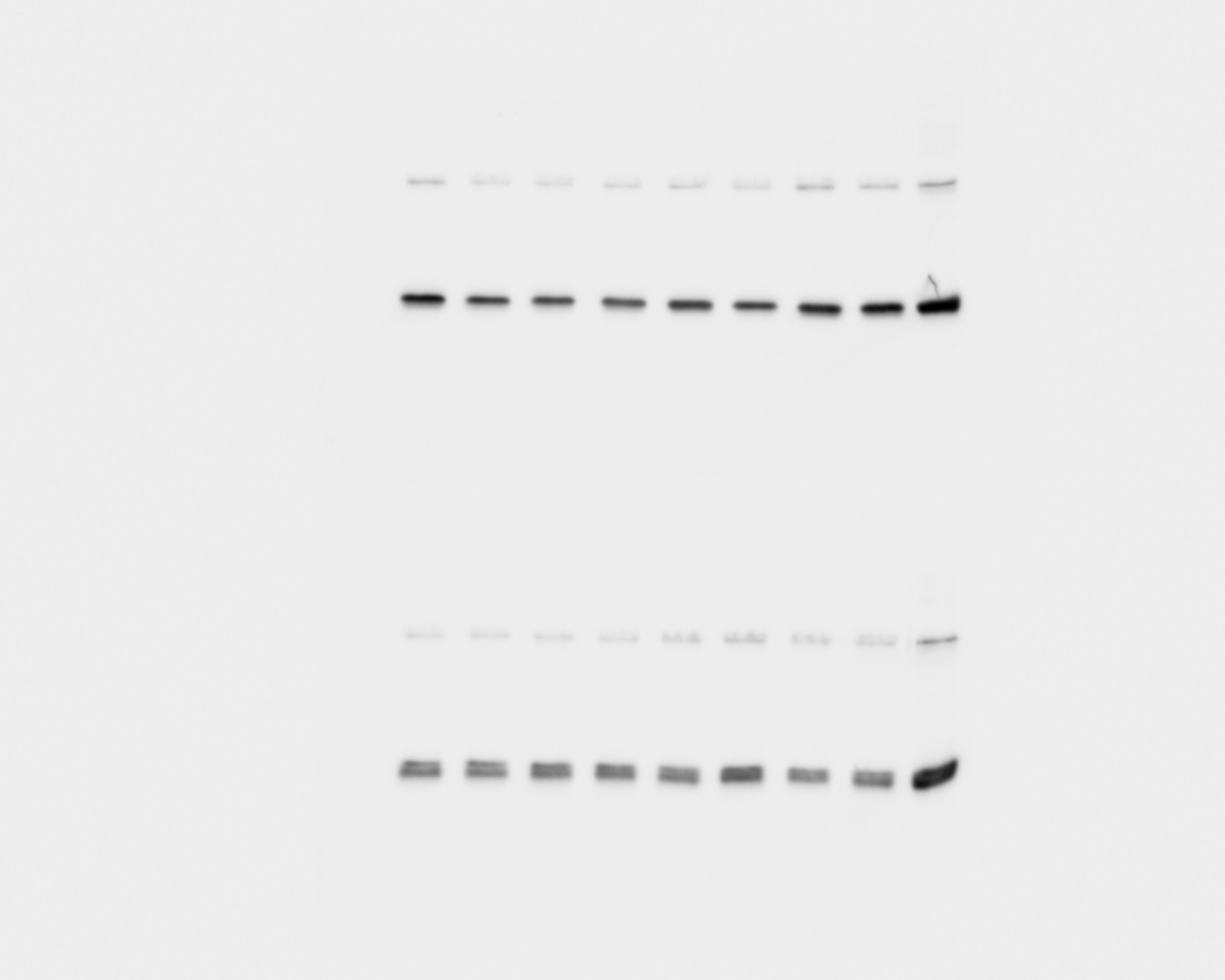

Exposition time = 11 seconds

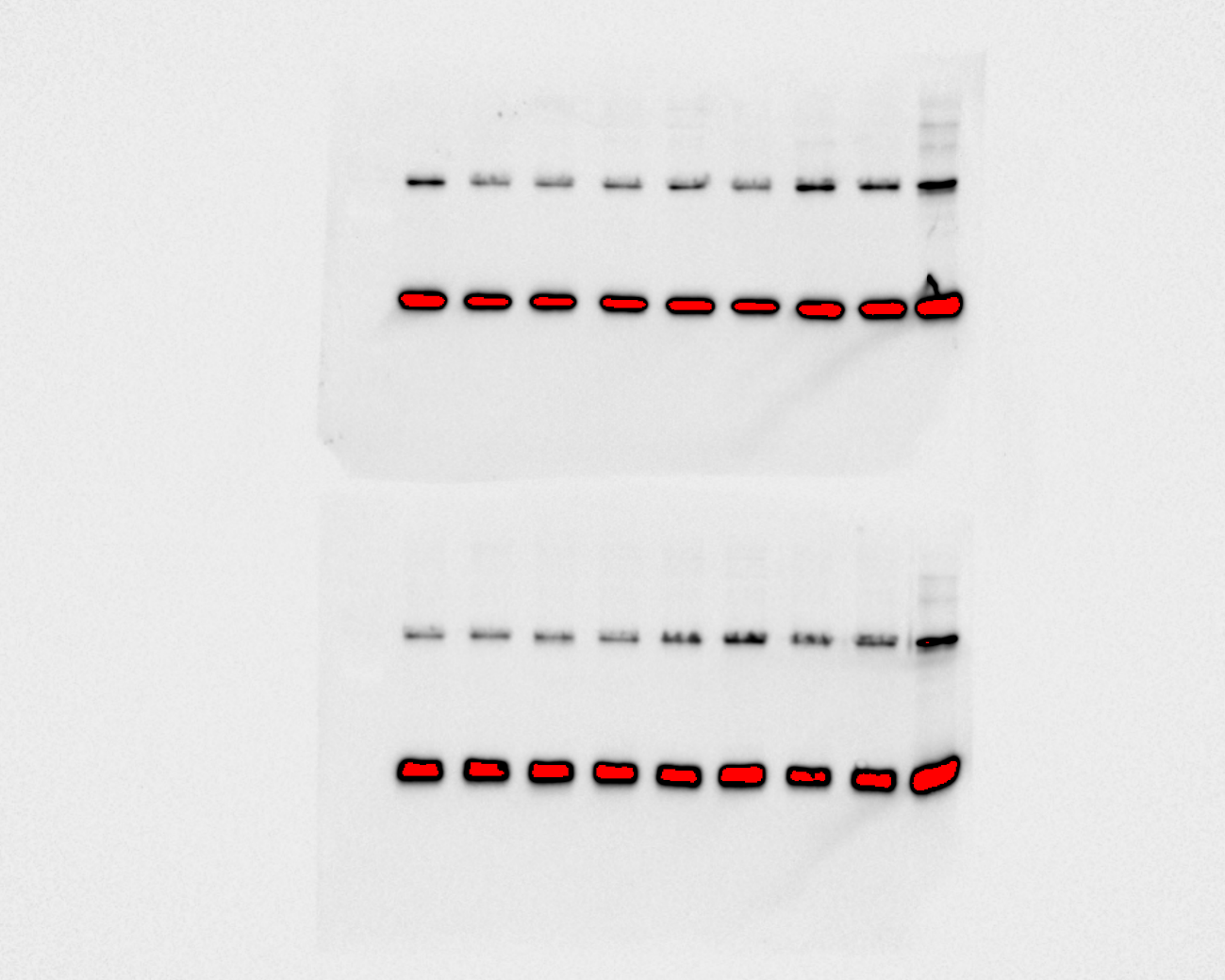

**Blot 7 and 8**

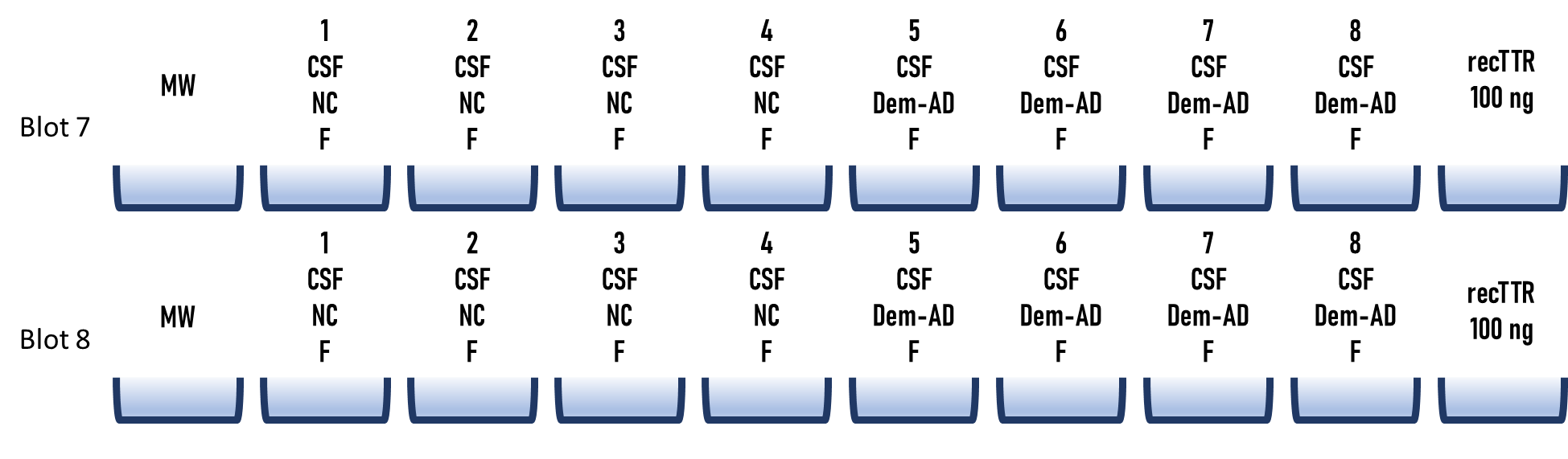

Exposition time = 1 second

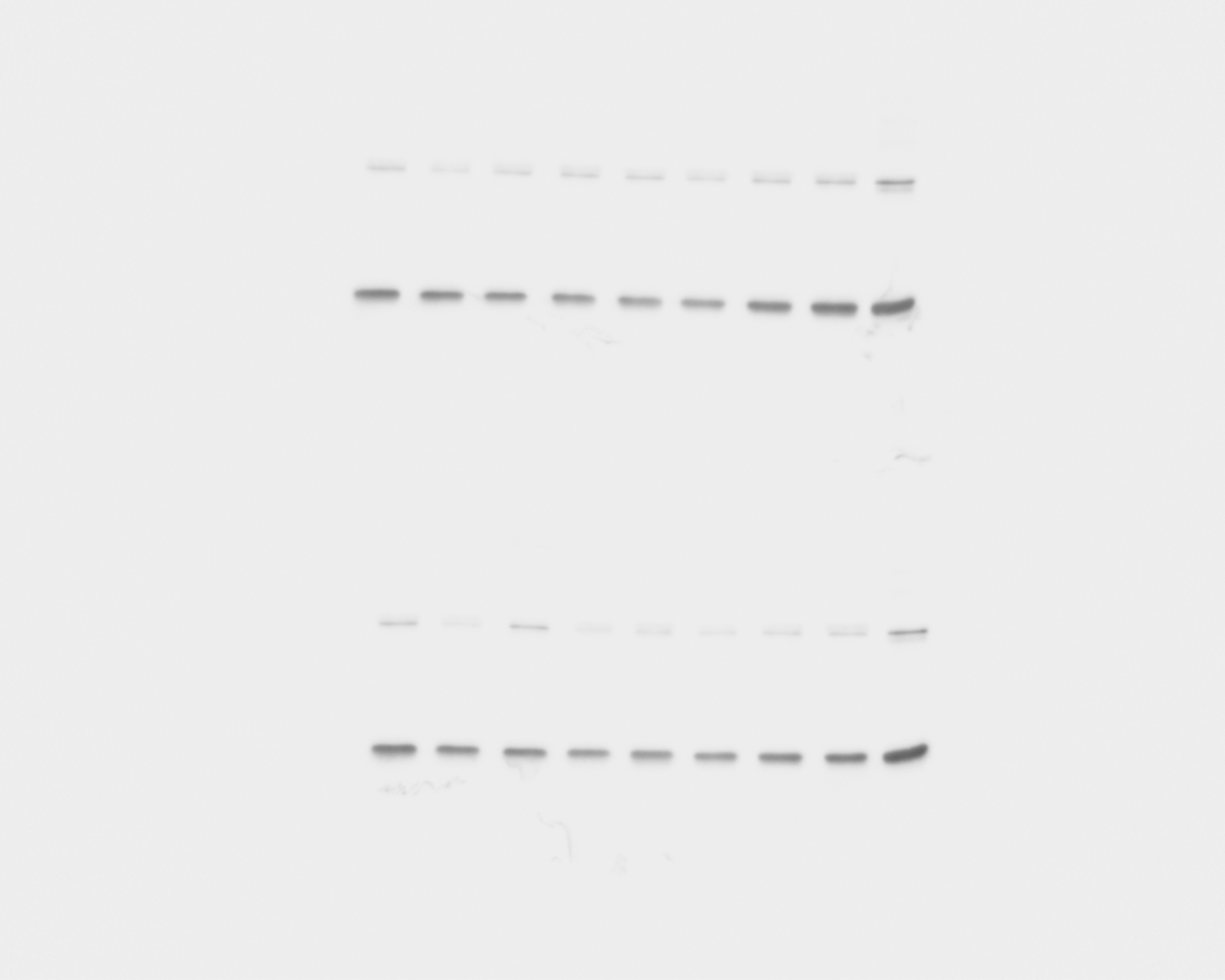

Exposition time = 11 seconds

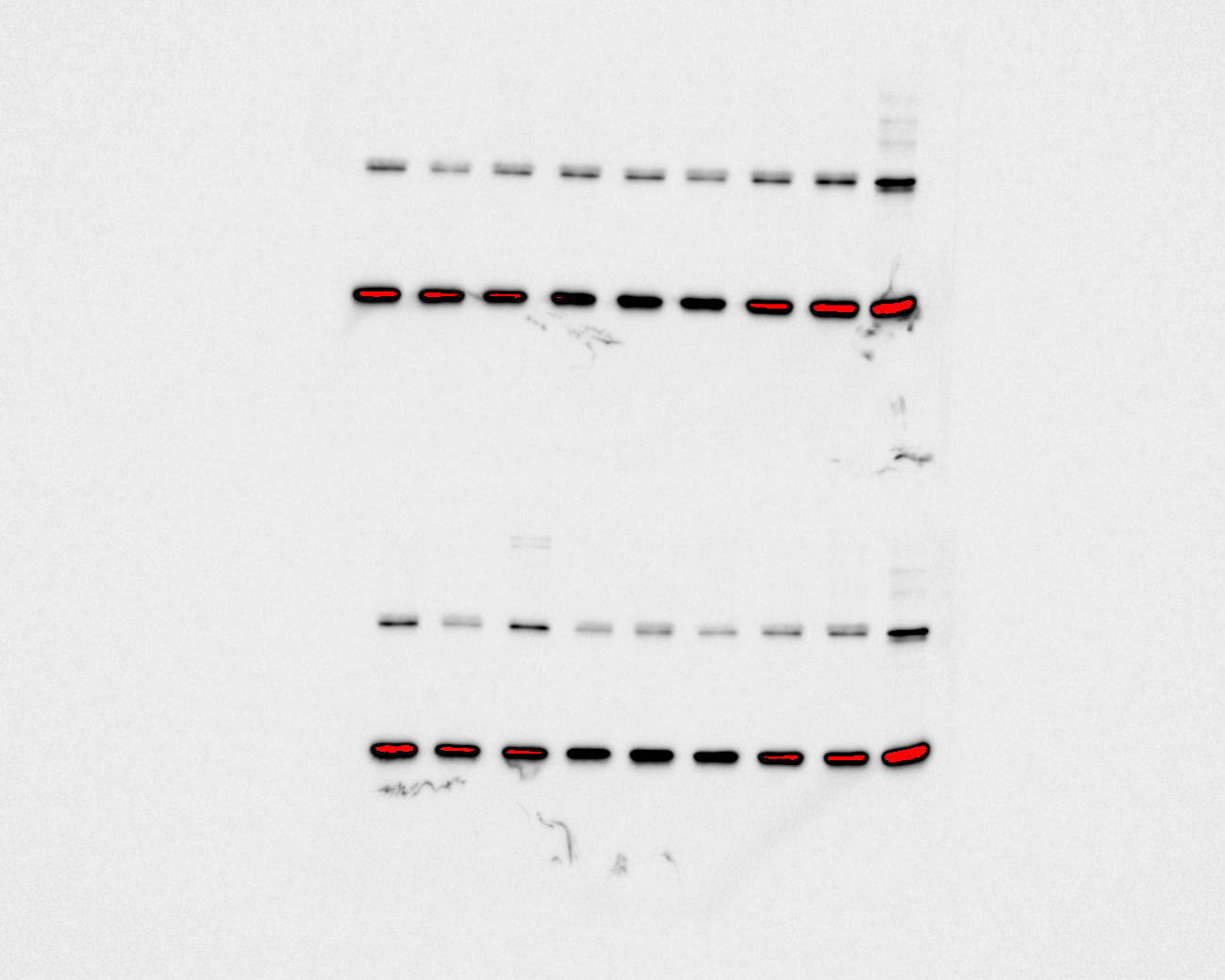

**Blot 9 and 10**

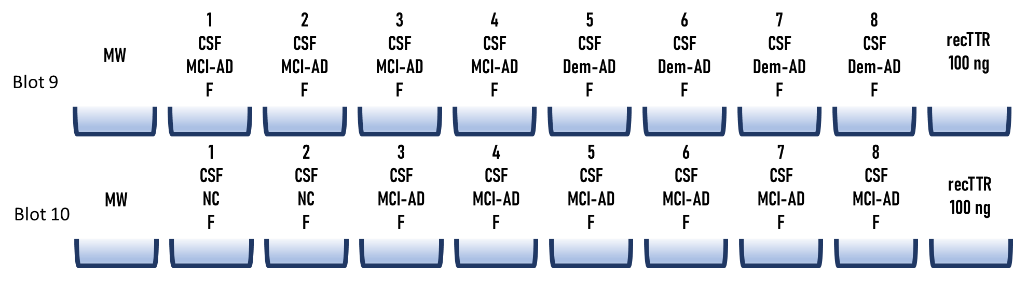

Exposition time = 1 second

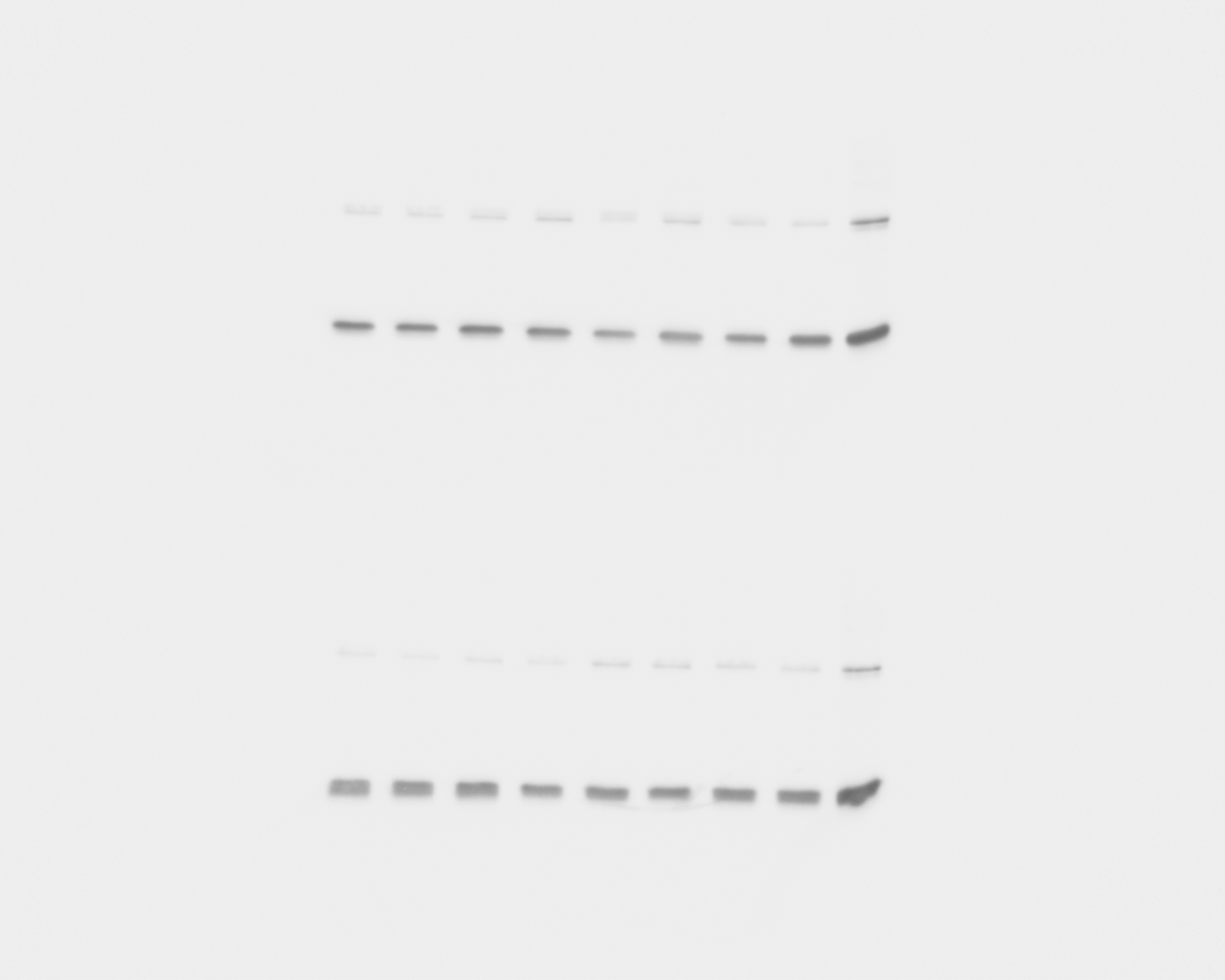

Exposition time = 11 seconds

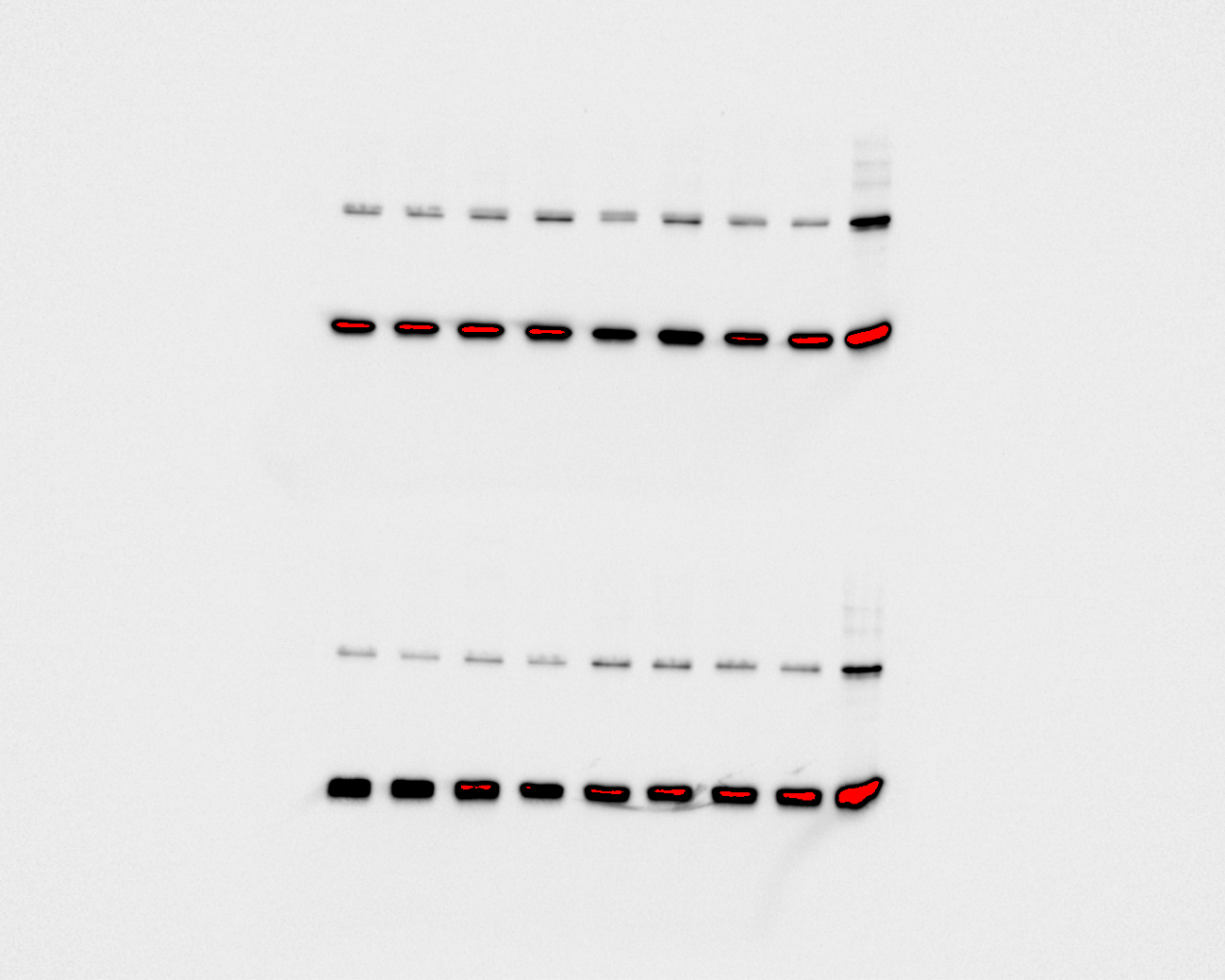

**Blot 11 and 12**

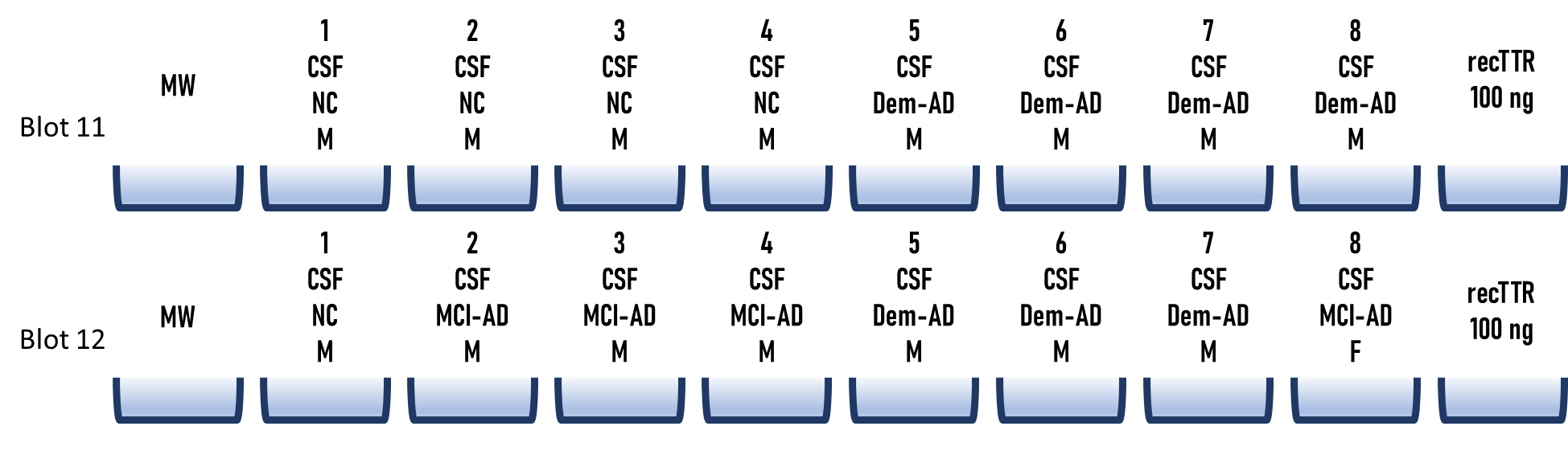

Exposition time = 1 second

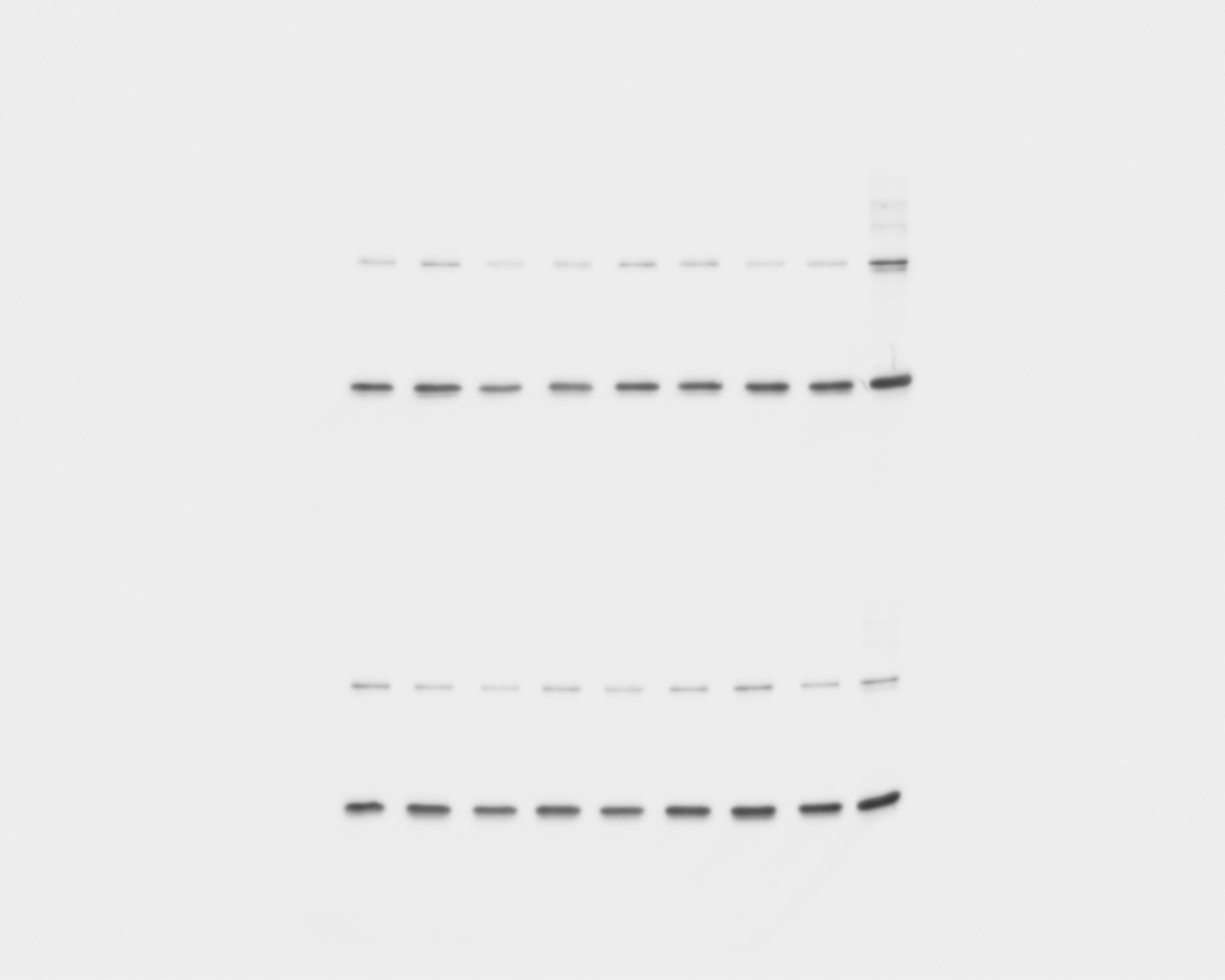

Exposition time = 11 seconds
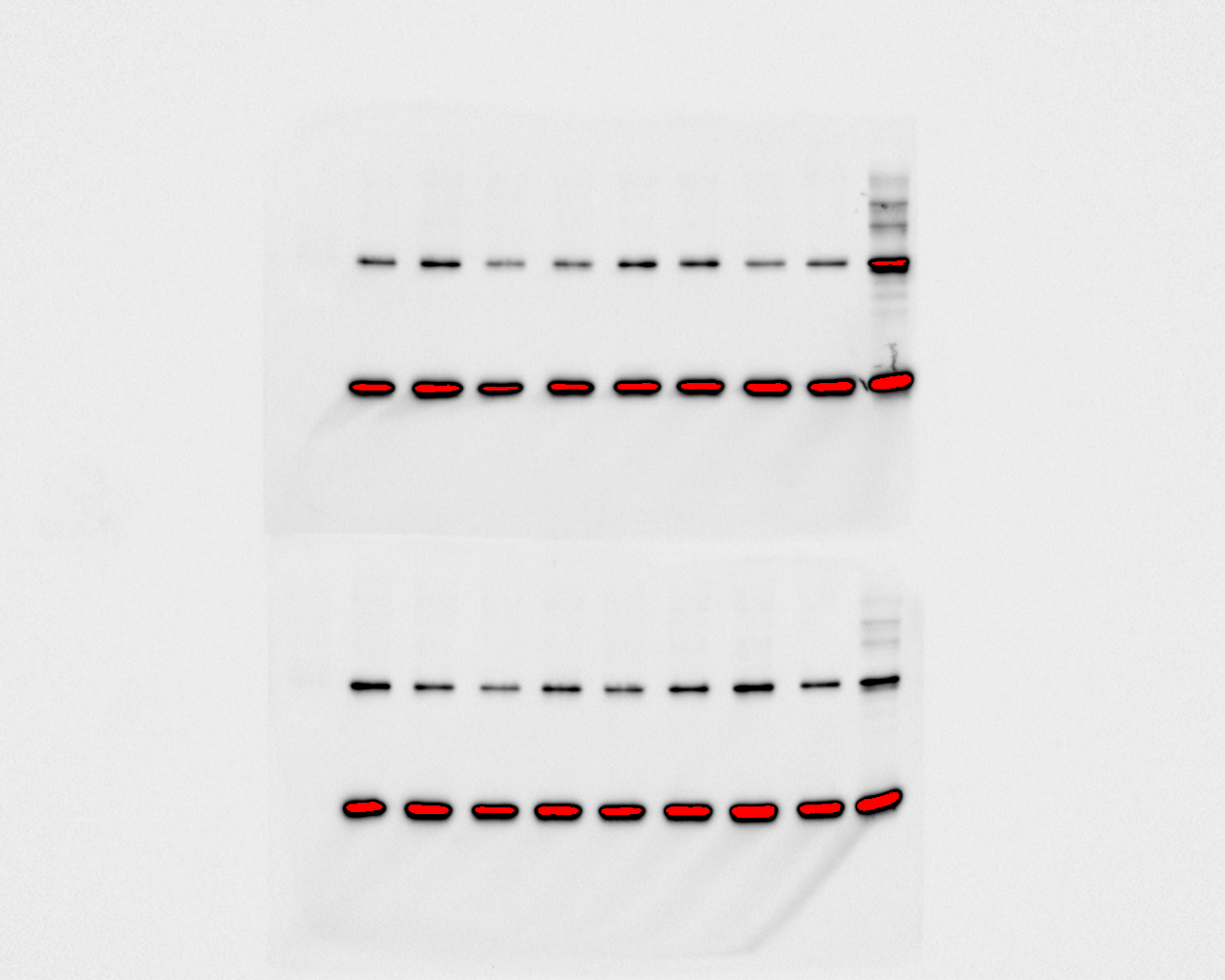

**2.2. Evaluation of CSF instability – 2nd set of blots (12 blots)**

**Blot 1 and 2**

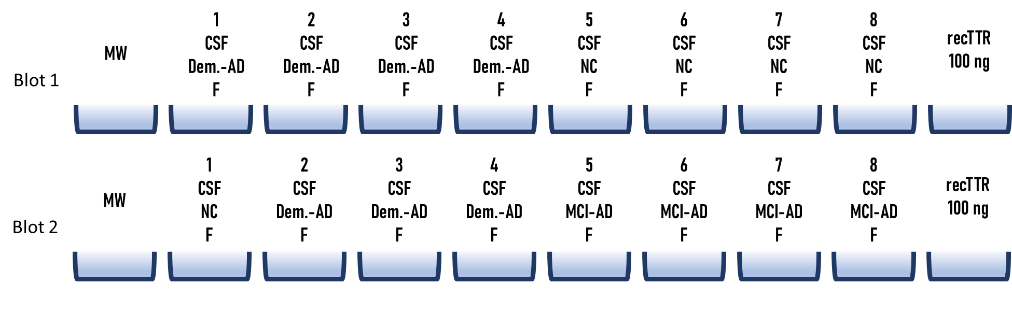

Exposition time = 1 second

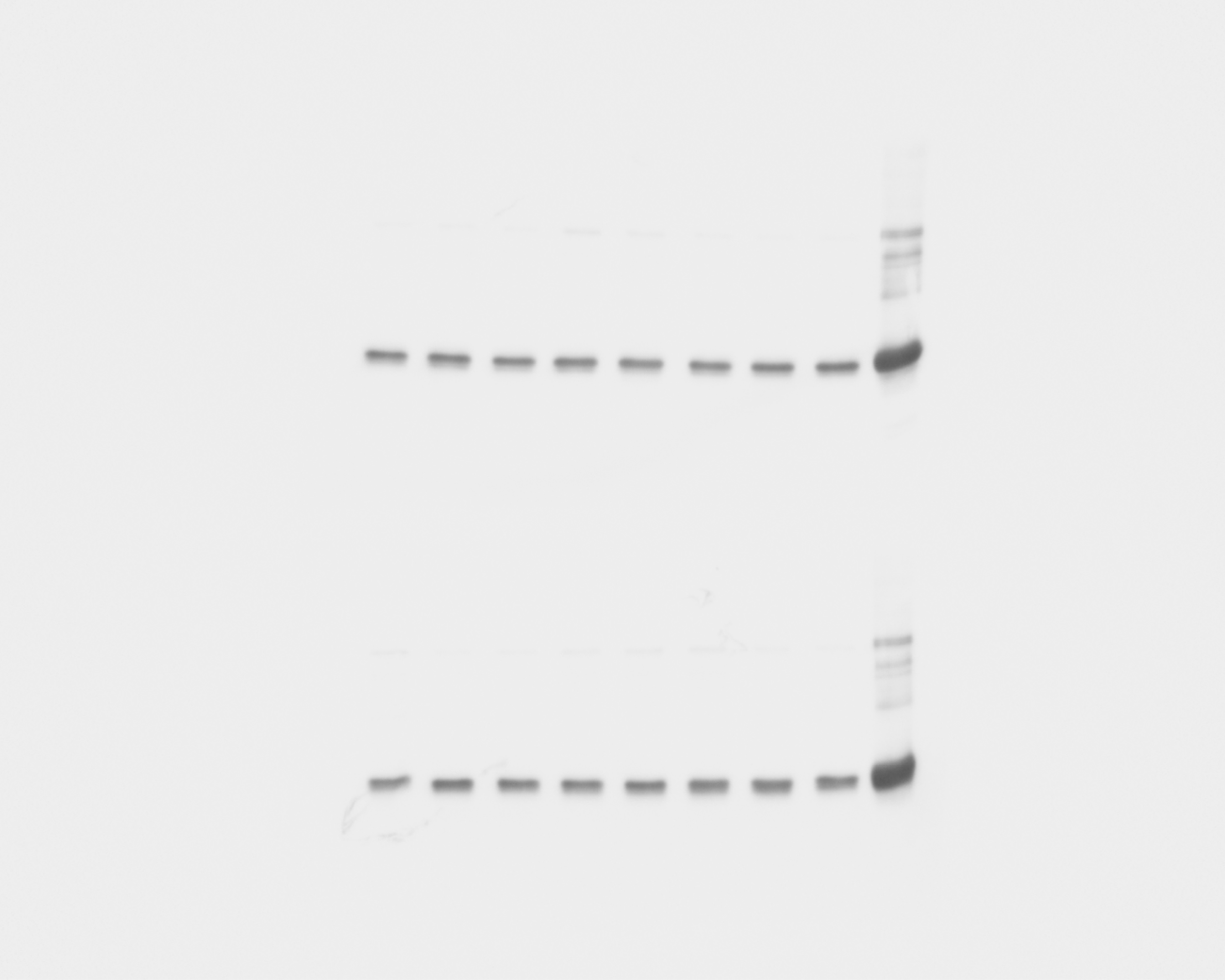

Exposition time = 11 seconds

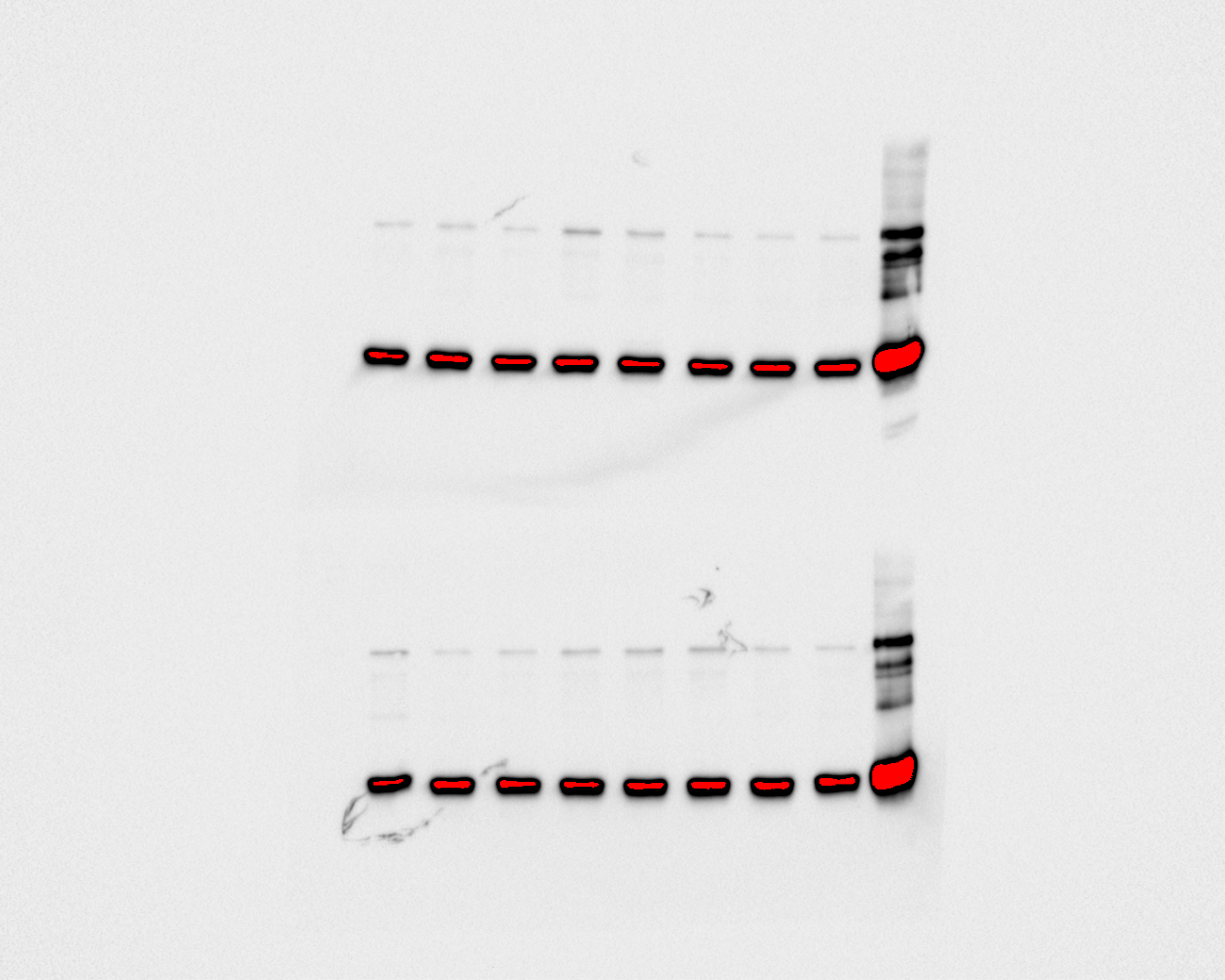

**Blot 3 and 4**

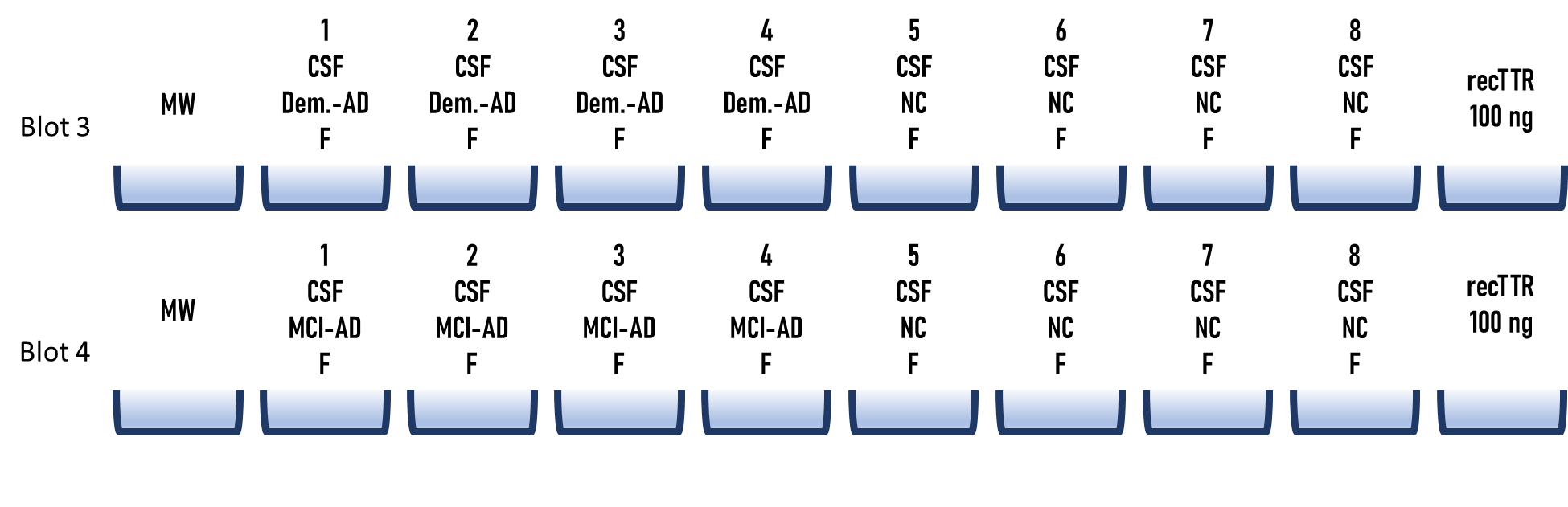

Exposition time = 1 second

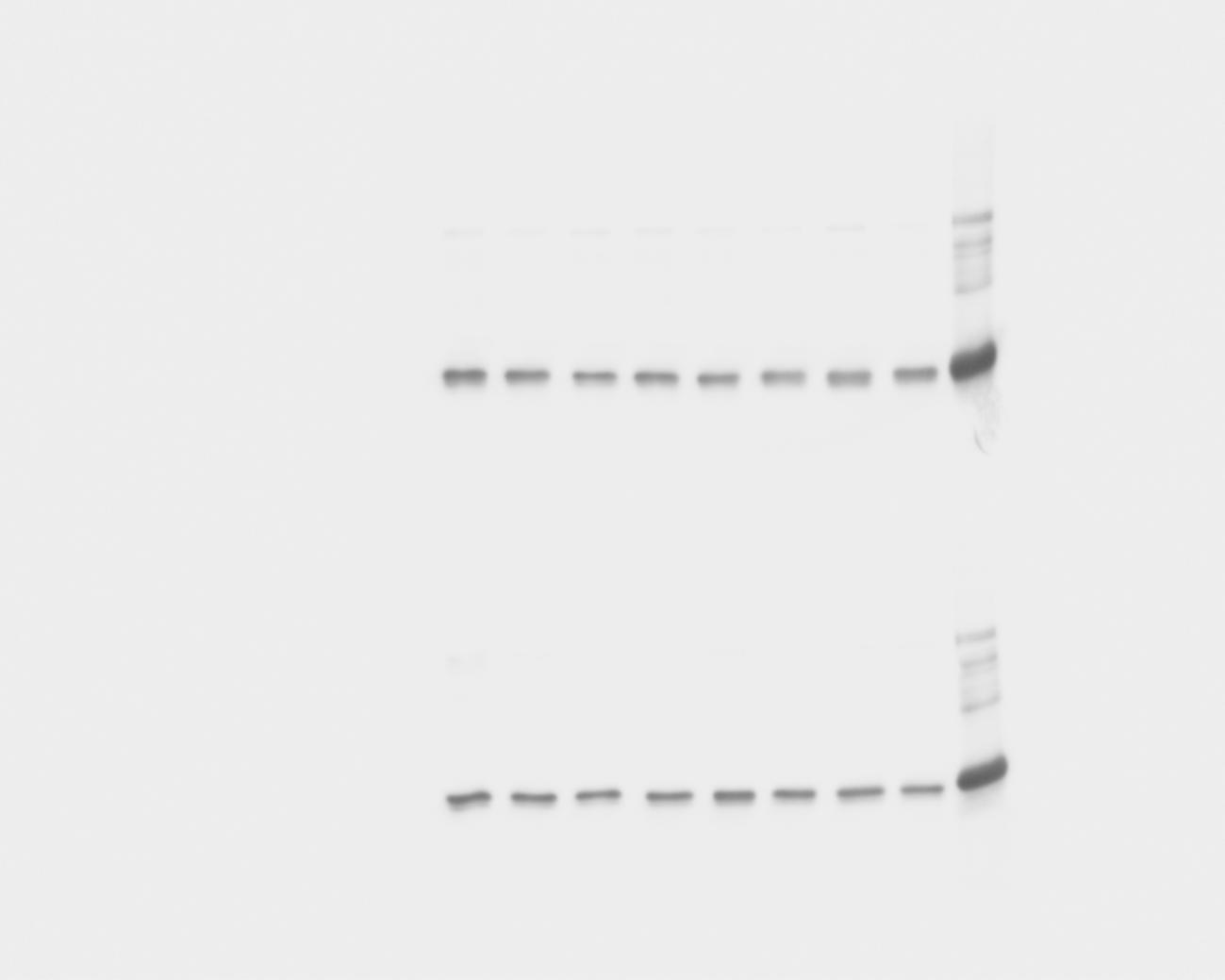

Exposition time = 11 seconds

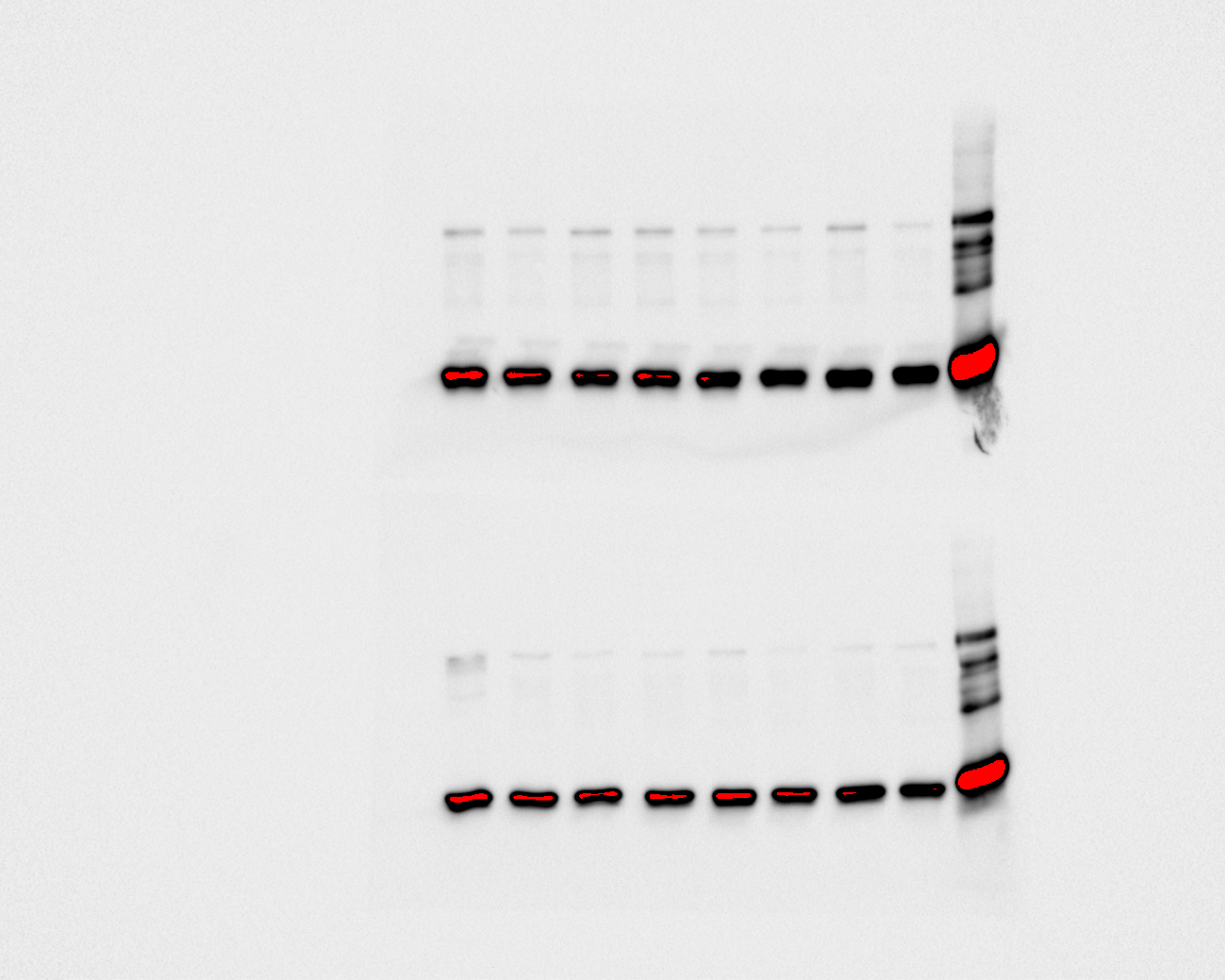

**Blot 5 and 6**

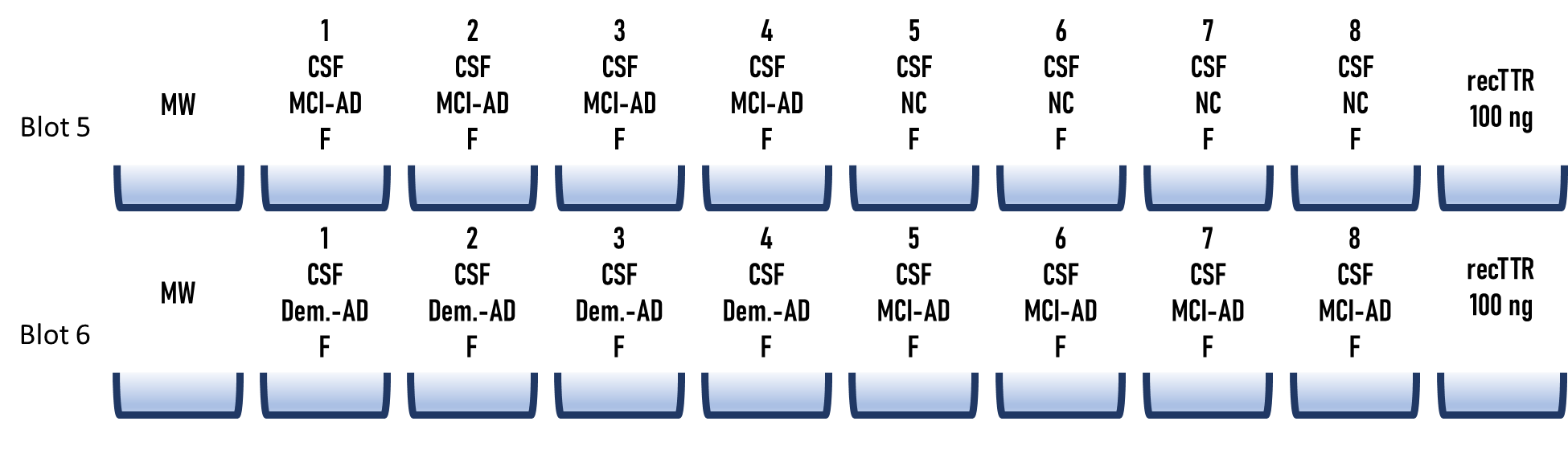

Exposition time = 1 second

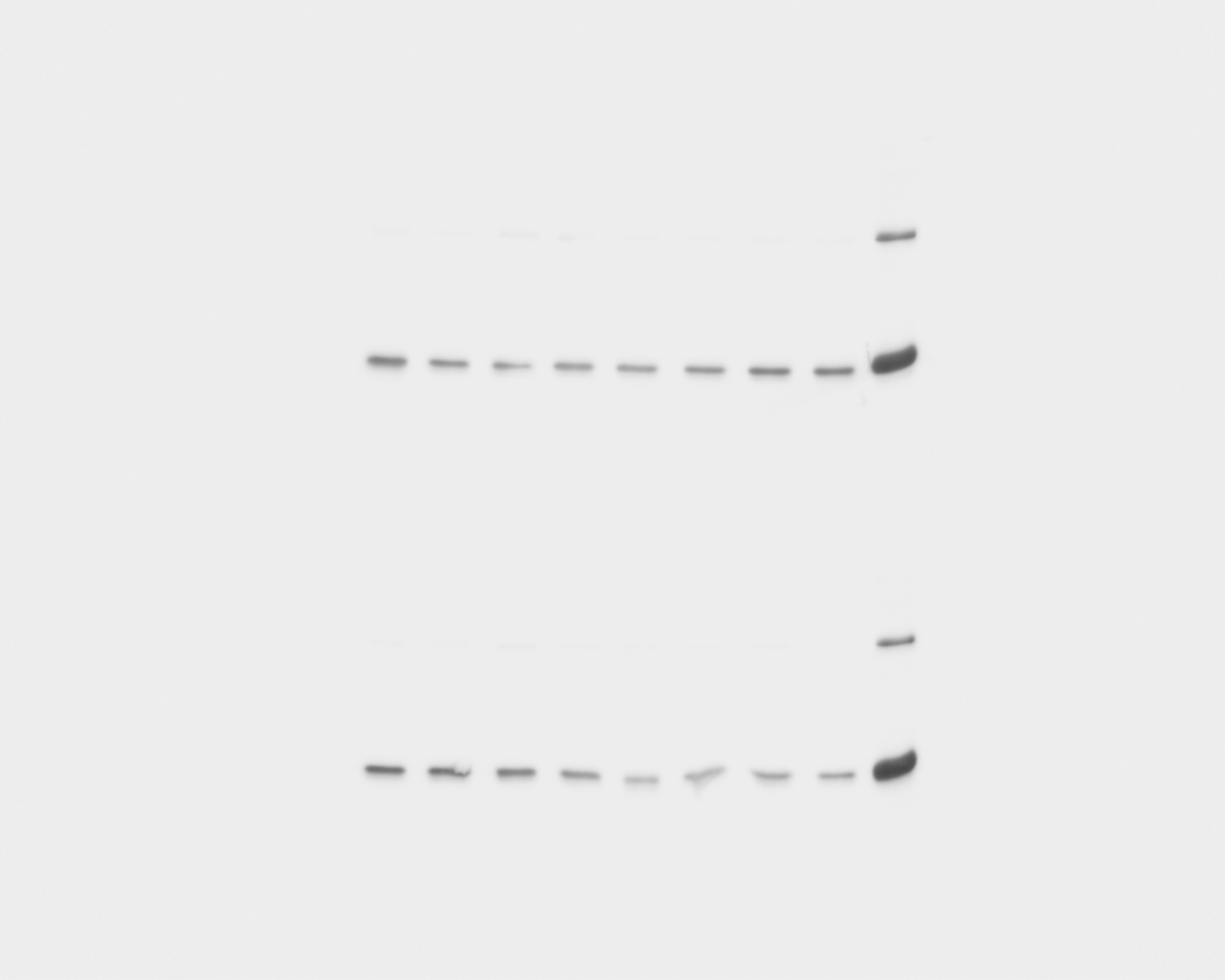

Exposition time = 11 seconds

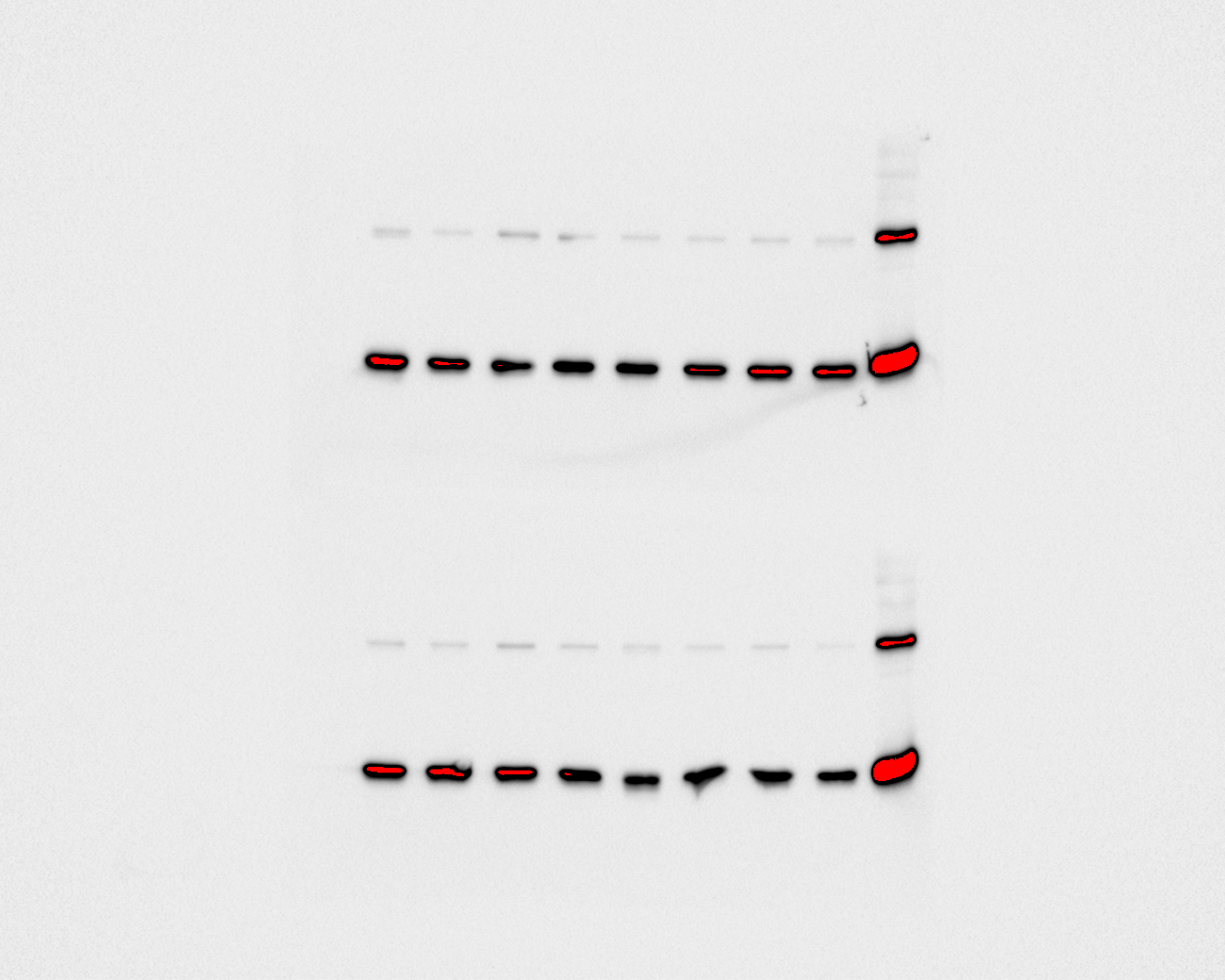

**Blot 7 and 8**

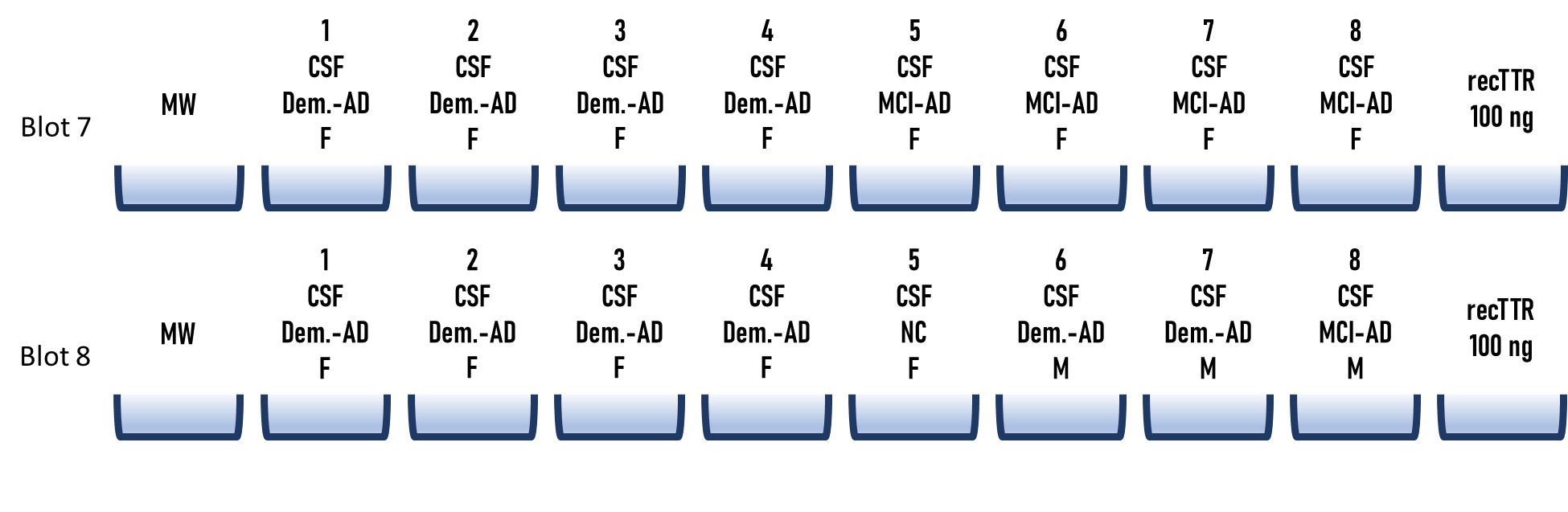

Exposition time = 1 second

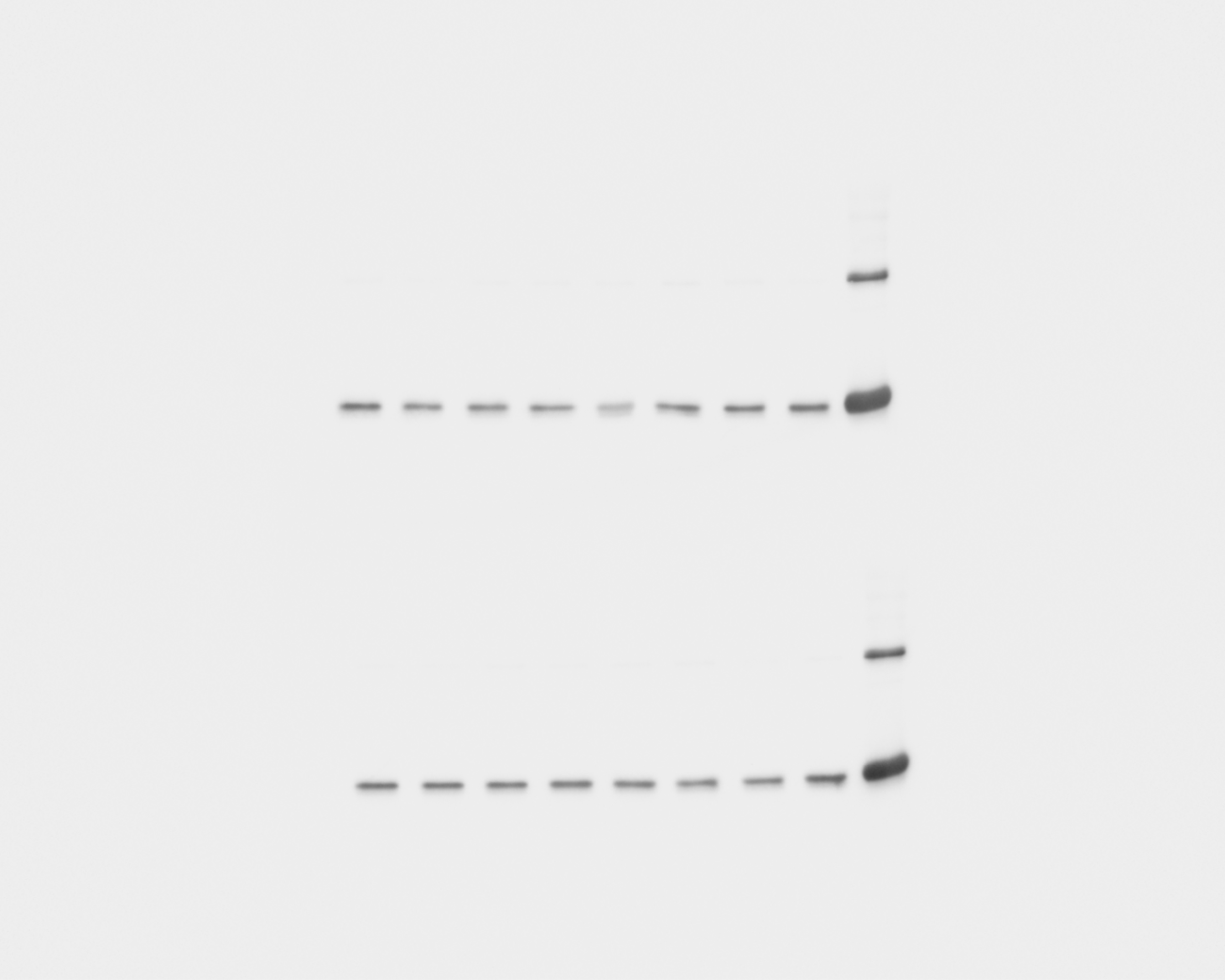

Exposition time = 11 seconds

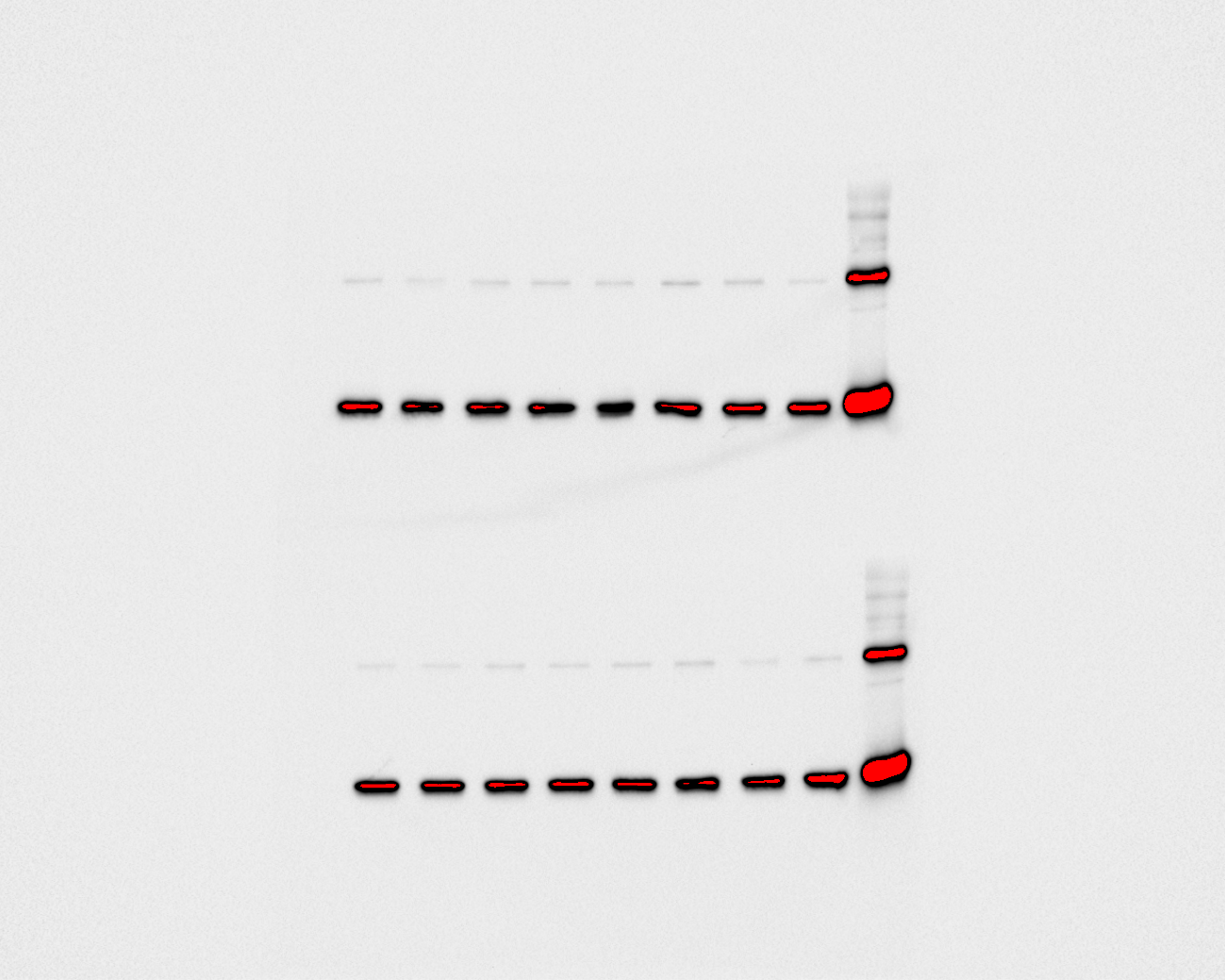

**Blot 9 and 10**

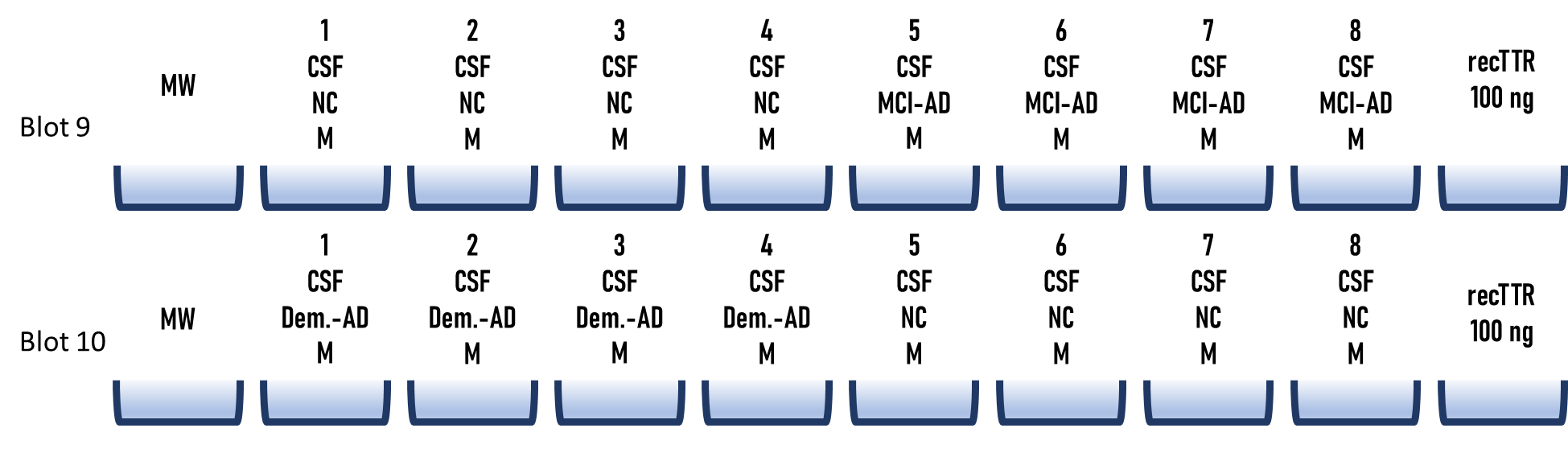

Exposition time = 1 second

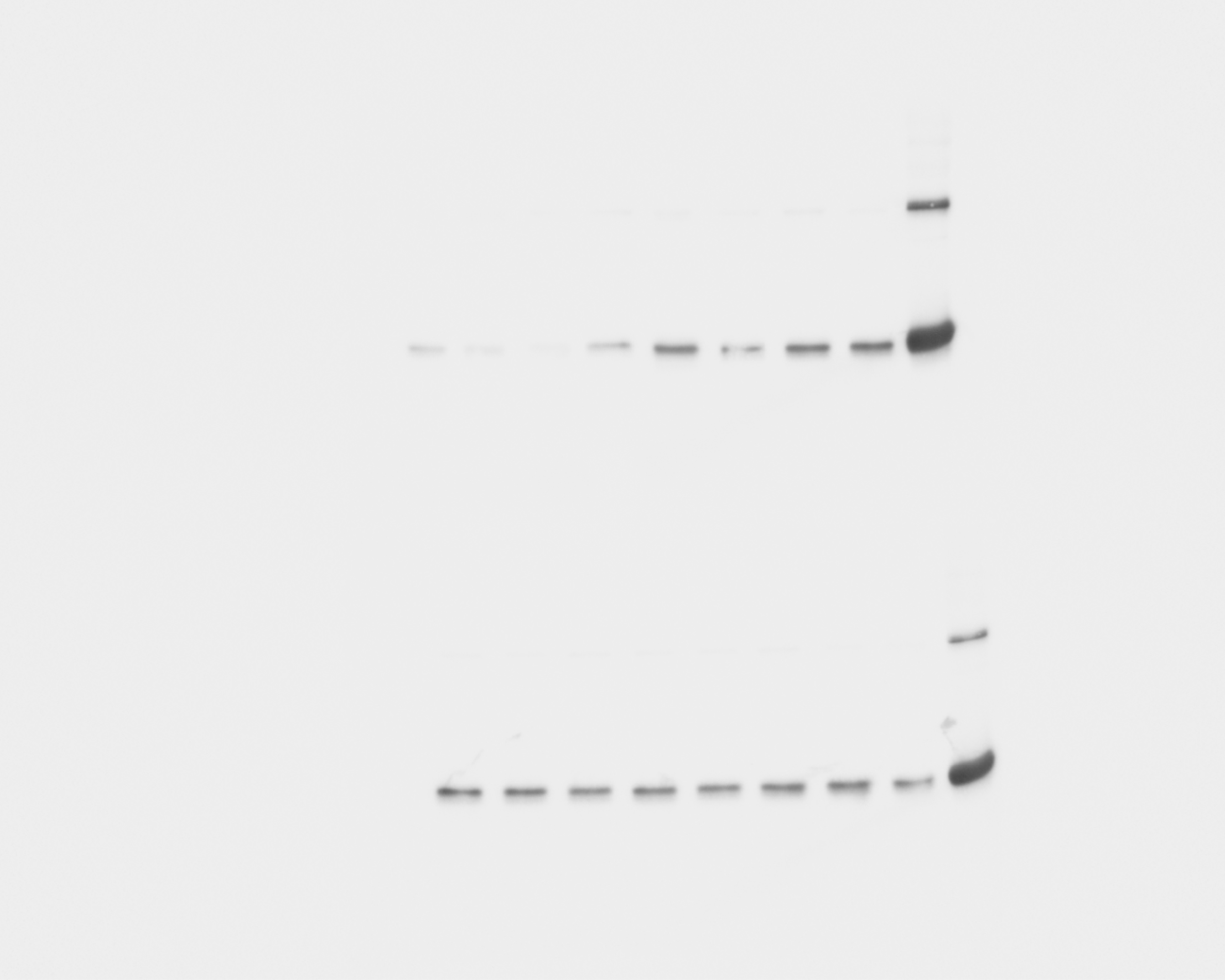

Exposition time = 11 seconds

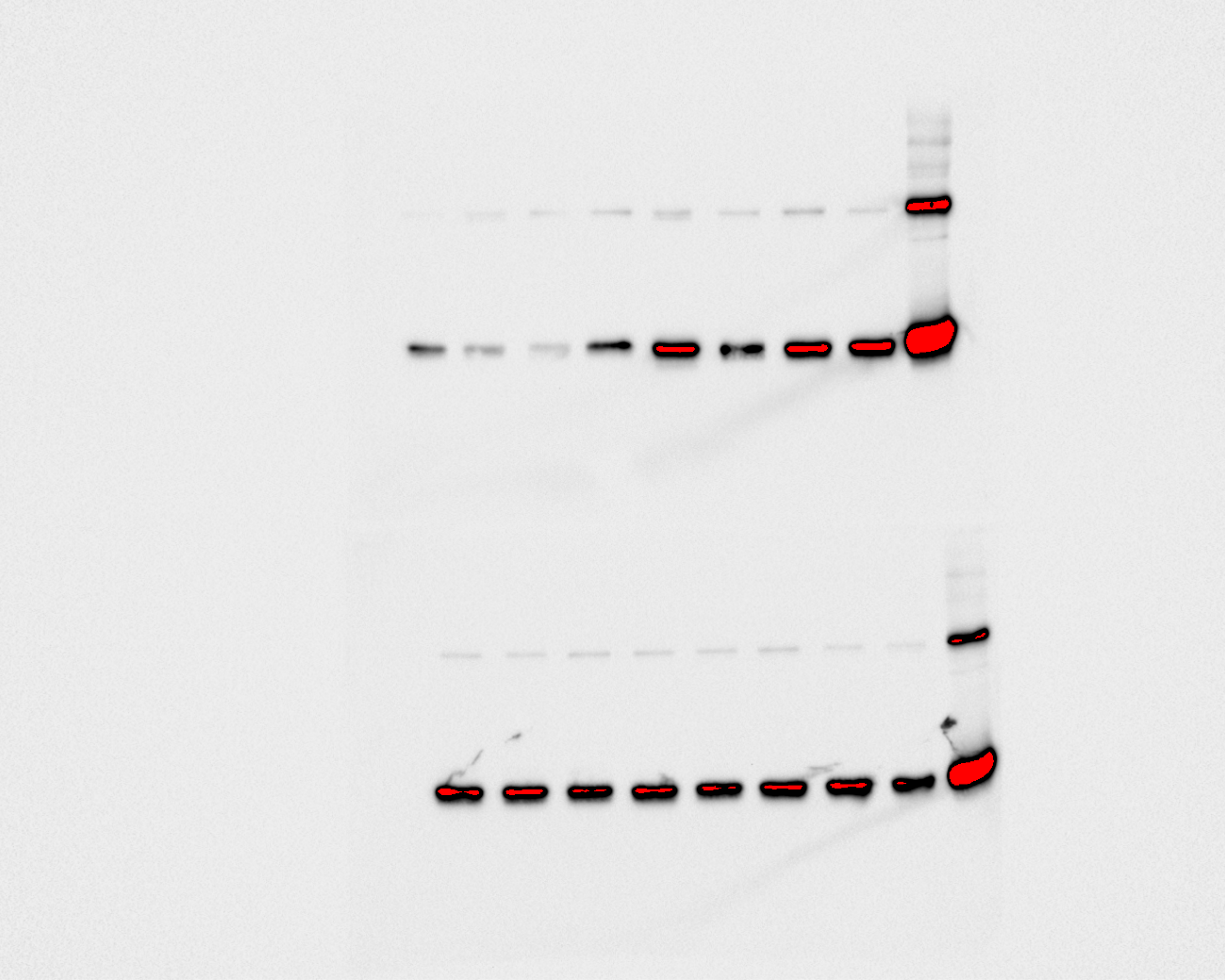

**Blot 11 and 12**

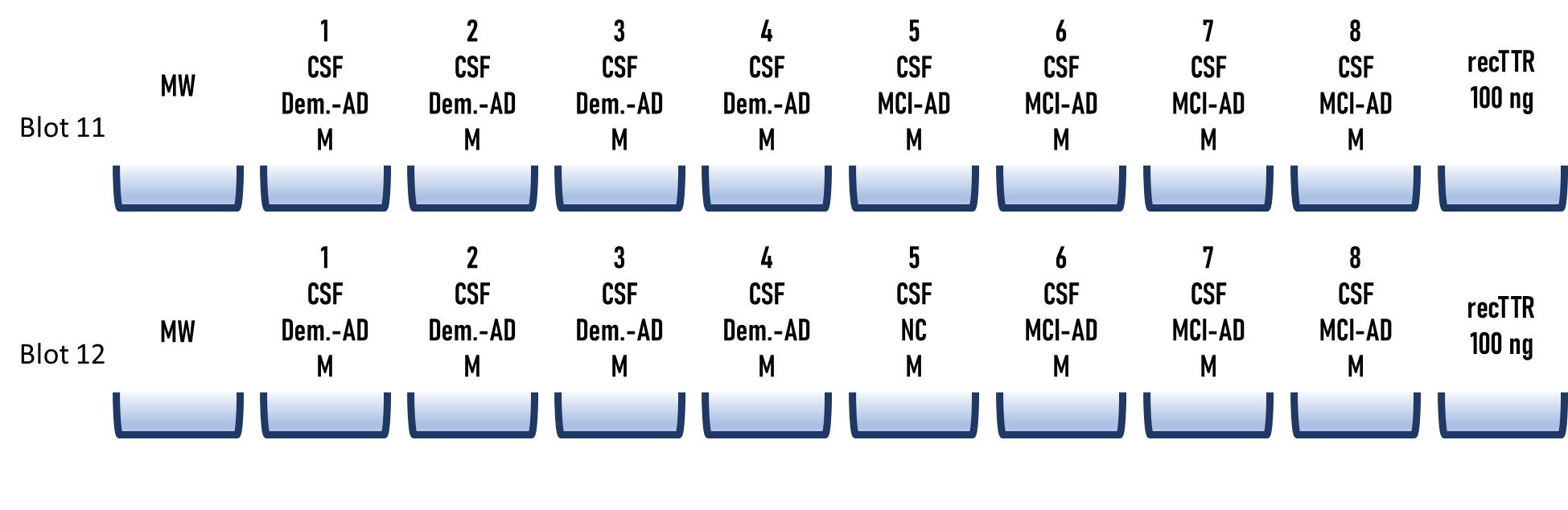

Exposition time = 1 second

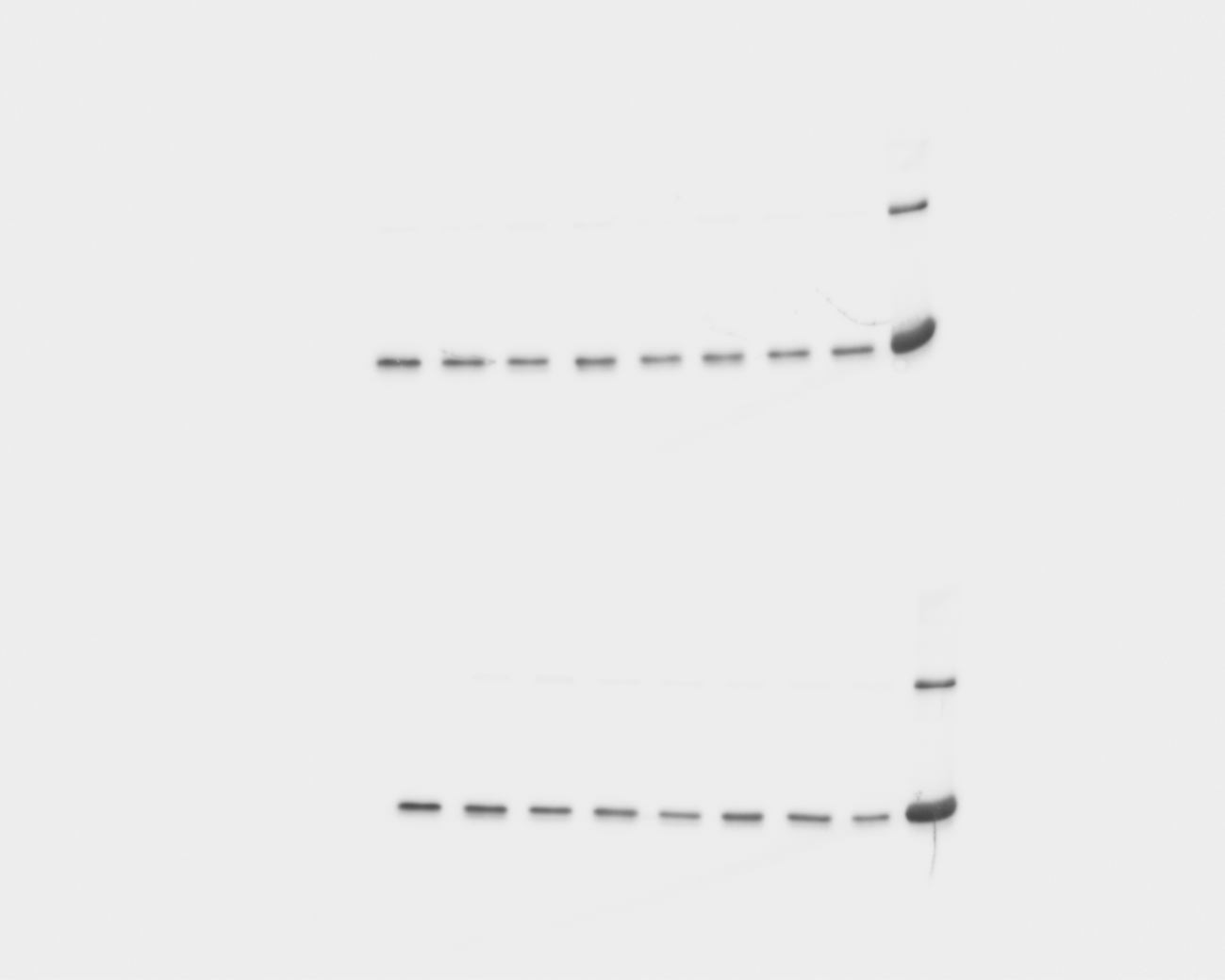

Exposition time = 11 seconds

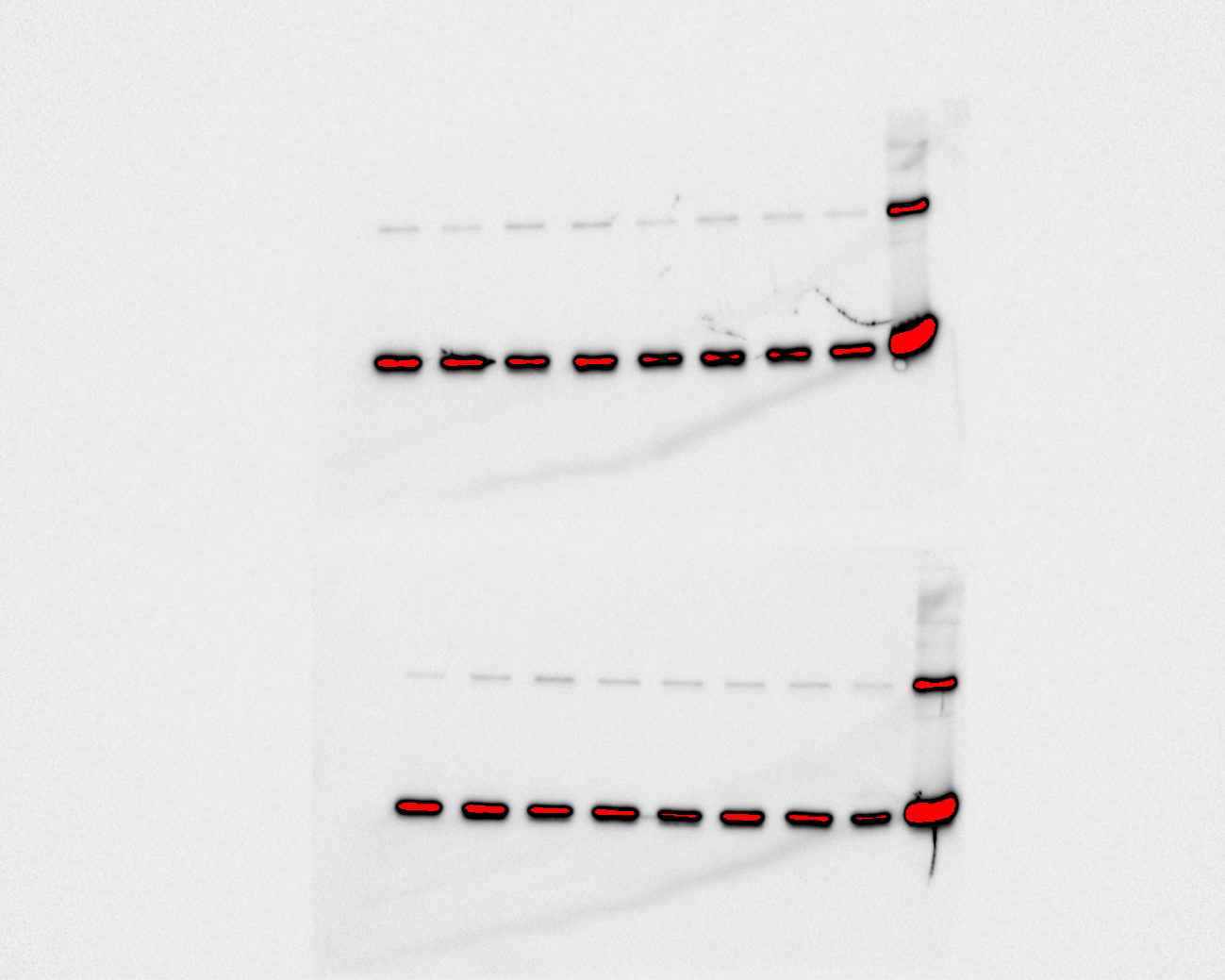

3. **Evaluation of the effect of TTR binding to Aβ42 peptide**

The image shows the uncropped blot presented in Figure 3
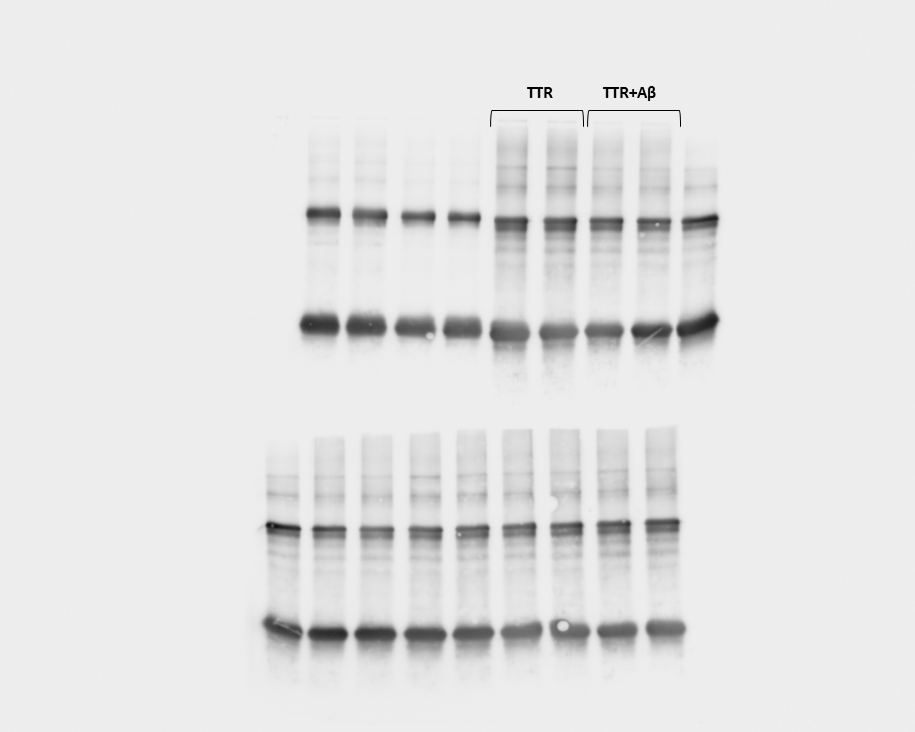
